# Complete genomes of a multi-generational pedigree to expand studies of genetic and epigenetic inheritance

**DOI:** 10.64898/2025.12.14.693655

**Authors:** Monika Cechova, Tamara A. Potapova, Andreas Rechtsteiner, Glenn Hickey, Rebecca Serra Mari, Mira Mastoras, Julian Menendez, Nikol Poláková, Prajna Hebbar, Fedor Ryabov, Hailey Loucks, Aljona Groot, Tomáš Pavlík, Mobin Asri, Shihua Dong, Stephanie M. Yan, Julian K. Lucas, Steven J. Solar, Matthew Borchers, Mark Mattingly, Sean McKinney, Marie Krátká, Catherine Mikhailova, Ondřej Hanák, Sohinee Tiffany Saha, Emily Xu, Dmitry Antipov, Sergey Koren, Jordan M. Eizenga, Brandy McNulty, Joshua M.V. Gardner, Todd Hillaker, Ivo Violich, Christopher Markovic, Semyon Kruglyak, Shawn Levy, Trevor Wolf, Matthew W. Mitchell, Laura Scheinfeldt, Haoyu Cheng, Ivan A. Alexandrov, Rajiv C. McCoy, Benedict Paten, Adam M. Phillippy, Justin M. Zook, Jennifer L. Gerton, Robert S. Fulton, Nathan O. Stitziel, Ting Wang, Tobias Marschall, Carol W. Greider, Karen H. Miga

## Abstract

Pedigree analysis remains the gold standard for rare disease diagnostics, yet whole genome sequencing studies typically omit critical regions like centromeres, telomeres, and acrocentric chromosome p-arms. Here, we present telomere-to-telomere (T2T) reference genomes for four self-identified African American individuals of admixed ancestry spanning three generations. Our parent-of-origin assigned, chromosome-level assemblies revealed precise meiotic recombination breakpoints in previously inaccessible regions, including recombination events across acrocentric and subtelomeric sequences. Centromeric regions were highly stable, with multi-megabase arrays inherited intact across three generations, while the position of kinetochore assembly sites remained consistent and predominantly associated with the p-arm proximal region. The relative lengths of telomeres on individual chromosomes were maintained across generations. Using a targeted rDNA assembly approach, we reconstructed a complete megabase-scale ribosomal DNA (rDNA) array corresponding to the paternal chromosome 14. This openly available pedigree provides a benchmark dataset for studying recombination and genetic and epigenetic variation across the complete genome.

## Introduction

Repeat-rich heterochromatic regions, including centromeres, telomeres, and the acrocentric short arms, are essential for cell viability yet remain among the least understood parts of the genome. These loci have critical roles in chromosome segregation, cellular aging, and ribosome biogenesis (Verdaasdonk and Bloom 2011; Aubert and Lansdorp 2008; McStay 2023), and their dysregulation has distinct clinical consequences. Centromere dysfunction can lead to aneuploidy, a leading driver of pregnancy loss and cancer (Barra and Fachinetti 2018; Essers et al. 2023). Progressive telomere erosion imposes replicative limits that drive cellular senescence and tumor evolution; malignant cells overcome these limits through telomerase reactivation or alternative lengthening pathways (Harley et al. 1990; Sieverling et al. 2020). The acrocentric short arms contain megabase-scale rDNA arrays required for protein synthesis, where variation is linked to renal function, hematological and other traits (Rodriguez-Algarra et al. 2024, 2025). Beyond individual loci, these regions are also prone to large-scale structural instability: for example, repeat classes on acrocentric chromosomes present hotspots for Robertsonian translocations, which are among the most common human chromosomal rearrangements (de Lima et al. 2025; Guarracino et al. 2023; Gerton 2024). Despite their biological and clinical relevance, these regions have remained excluded from genomic analysis due to their highly repetitive sequence composition, leaving a critical gap in our understanding of how they evolve and are transmitted across generations.

Telomere-to-telomere (T2T) assemblies now provide high-resolution access to previously inaccessible repeat-rich regions, enabling direct study of their roles in genome stability, cellular function, and disease. The T2T-CHM13 assembly provided the first fully complete haploid reference (Nurk et al. 2022), revealing approximately 200 million base pairs of previously unresolved sequence including complete centromeres, segmental duplications, and p-arms of acrocentric chromosomes. Subsequent efforts achieved a complete diploid benchmark (HG002-Q100) with near-perfect accuracy across both haplotypes, adding approximately 700 Mb of confidently resolved autosomal sequence to this widely used benchmark (Hansen et al. 2025). However, producing fully complete, error-free T2T assemblies requires significant expertise, costly multi-platform data, and considerable manual oversight. Consequently, large-scale projects have adopted near-T2T assemblies that, while not yet matching the completeness of manually curated reference genomes, represent an advance in contiguity and overall assembly quality. For example, the Platinum Pedigree study (Porubsky et al. 2025; Kronenberg et al. 2025) established that near-T2T methods can resolve inheritance and de novo mutation rates across four generations, despite incomplete coverage of regions such as the acrocentric p-arms. Similarly, the Human Genome Structural Variation Consortium (HGSVC) (Logsdon, Ebert, et al. 2024) produced hundreds of nearly complete genomes at population scale, greatly expanding the catalog of complex variation. Nonetheless, centromeric satellites, subtelomeres, and the acrocentric short arms remain among the most challenging regions to assemble, and are consistently underrepresented. While technical challenges remain for specific genomic regions, the field has now reached a point where high-quality T2T assemblies can be applied to answer fundamental biological questions. A critical next step is extending fully curated T2T assemblies to pedigrees, which enables direct measurement of Mendelian inheritance and *de novo* mutation rates across both single-nucleotide and structural variants. In this work, we present complete T2T assemblies for four family members spanning three generations, providing a comprehensive view of genetic inheritance across the entire genome.

Centromeric satellite arrays are among the most repetitive and structurally complex regions of the human genome, where sequence turnover and epigenetic specification jointly support kinetochore function (Henikoff et al. 2001; Altemose et al. 2022). Population-scale comparisons reveal extensive structural variation: arrays differ by megabases between individuals and evolve rapidly compared to chromosome arms through unequal crossover (Vincenten et al. 2015), non-allelic homologous recombination (Onaka et al. 2016), and break-induced replication (Miga 2019; Logsdon, Rozanski, et al. 2024; Logsdon, Ebert, et al. 2024; Showman et al. 2024). Family-based sequencing studies demonstrate that, despite large-scale population variation, centromeric arrays remain largely unchanged across meioses. Early work tracking the DXZ1 array on the X chromosome across 84 meiotic events and ∼191 Mb of alpha satellite DNA detected no recombination within centromeric arrays across three-generation families (Wevrick and Willard 1989), and recent analysis confirms this pattern, showing no observable centromere array length changes across transmissions, with only ∼4.4 *de novo* mutations per meiosis (Porubsky et al. 2025). This intergenerational stability contrasts sharply with the arrays’ role in aneuploidy and cancer, where centromere dysfunction drives chromosome missegregation (Barra and Fachinetti 2018). Understanding how centromeric arrays maintain stability during normal inheritance while contributing to genomic instability in disease requires following complete, fully resolved arrays through multiple generations.

The acrocentric short arms (chromosomes 13, 14, 15, 21, and 22) contain large, megabase-sized rDNA arrays, which are essential for ribosome biogenesis, where copy number variation associates with hematological traits and renal function (Rodriguez-Algarra, Evans, and Rakyan 2024). Population studies reveal considerable variation in both rDNA copy number and array organization, yet pedigree analyses show family-specific profiles that segregate as stable Mendelian loci with suppressed recombination (Petes and Botstein 1977; Gibbons et al. 2015; Potapova et al. 2025). The homology among acrocentric short arms enables non-homologous exchanges, creating pseudo-homologous regions capable of ongoing recombination (Guarracino et al. 2023; Ramos et al. 2019). Homology between these chromosomes supports polymorphic variation, exemplified by the D15Z1 satellites detected on non-homologous acrocentrics in ∼20% of individuals (Guarracino et al. 2023; Ramos et al. 2019). Robertsonian translocations occur in approximately 1 in 800 births (Hamerton et al. 1975; Nielsen and Wohlert 1991; Zhao et al. 2015). While they represent only ∼3-4% of Down syndrome cases, maternal balanced carriers face recurrence risks as high as 10-15%, highlighting the clinical impact of acrocentric short arm instability (Kolgeci et al. 2012; Wilch and Morton 2018; Mutton et al. 1996). Despite their clinical importance and known population-level variation, the transmission dynamics of complete acrocentric short arms, including rDNA array organization, satellite composition, and inter-chromosomal exchange, have never been directly observed at base-pair resolution across generations.

The transmission dynamics of subtelomeric regions and telomeres have remained poorly characterized despite the established role of telomere length in cellular senescence. Subtelomeres are enriched in segmental duplications and act as recombination hotspots, driving substantial structural variation and expansion–contraction across generations (Mefford and Trask 2002; Linardopoulou et al. 2005; Young et al. 2020). Rearrangements in these regions can disrupt dosage-sensitive genes and lead to developmental disorders (Sharp et al. 2005). Telomere length itself is strongly heritable (∼70%) (Slagboom et al. 1994; Broer et al. 2013; Honig et al. 2015) and influenced by parental origin effects, including offspring of older fathers having longer telomeres (Nordfjäll et al. 2005; Hjelmborg et al. 2015), but most studies have relied on bulk estimates that obscure chromosome-specific patterns. Recent work revealed that individual chromosome ends carry distinct chromosome-specific length profiles (Karimian et al. 2024), raising fundamental questions about how these profiles are established and inherited. Complete T2T assemblies now enable chromosome-by-chromosome analysis of telomere-subtelomere inheritance, making it possible to dissect maternal versus paternal contributions and track relative length rankings across generations, which bulk methods miss. Resolving these patterns is essential for understanding how telomere dynamics contribute to cellular senescence, and disease risk.

Here, we present the first complete telomere-to-telomere pedigree: four T2T genomes from a three-generation family who identify as African American, which resolves centromeres, telomeres, satellite sequences, and a subset of rDNA arrays across generations. This resource enables direct comparison of inherited haplotypes to quantify mutation rates, assess structural stability, and trace the inheritance of previously intractable medically relevant genomic features. By following these complex regions through Mendelian inheritance, we provide fundamental insights into their contribution to human genome evolution and establish a new paradigm for studying repeat-rich genomic regions. Our assemblies are released as an openly accessible community resource with matched cell line materials under broad consent, enabling population-scale studies of the complete human genome.

## Results

### Complete, highly accurate T2T assemblies of a multi-generational pedigree

We report a three-generation pedigree of family of admixed ancestry recruited in St. Louis, Missouri, to expand genomic reference resources from underrepresented populations. All participants self-identified as African American adults, and provided informed consent for open data sharing, broad research use, and iPSC derivation (Supplementary Files: Consent). The pedigree consists of the grandmother (HG06803; MGISTL-PAN010), grandfather (HG06804; MGISTL-PAN011), their progeny, here referred to as mother (HG06807; MGISTL-PAN027), and granddaughter (HG06808; MGISTL-PAN028). Lymphoblastoid cell lines (LCLs) were established for all four individuals and are available through the NHGRI Sample Repository for Human Genetic Research (NHGRI Repository) at the Coriell Institute for Medical Research, while induced pluripotent stem cell (iPSC) lines were so far successfully derived from three participants (HG06803, grandmother; HG06804, grandfather; and HG06807, mother) that are available through the NHGRI Repository. Karyotype analysis confirmed the diploid nature of samples (ISCN 46, XX or XY) except for the granddaughter (ISCN 45,X[5]/46,XX[15]), who exhibited mosaic Turner syndrome (Supplementary Files: Cytogenomics Report).

For this pedigree, we generated long- and short-read sequencing datasets from both blood-derived DNA and immortalized cell lines (Figure 1D, Supplementary Table 1 “Sequencing data”). PacBio high-fidelity (HiFi) sequencing from blood was obtained for all four individuals, ranging from 44× to 70× coverage. In addition, a matched blood and cell line HiFi dataset was produced for the mother (70× and 64×, respectively), providing an opportunity to directly compare sequencing performance across sample types. Oxford Nanopore Technology (ONT) Ultra-long nanopore datasets (ONT-UL) were generated at very high depth (169×–191× total coverage, including 70×–73× of reads ≥100 kb), providing continuity for assembly of highly repetitive regions. Additionally, ONT duplex sequencing from cell lines was performed with coverage between 18× and 25×. Chromatin conformation capture data were also produced, including ONT Pore-C (24.8×–31.5×) and Hi-C (Dovetail OmniC, 56×–100×) from cell lines, enabling haplotype phasing and scaffolding. Illumina whole-genome sequencing (WGS) short-read sequencing (150bp PE) was performed on both blood and cell line DNA for all individuals, with coverage ranging from 7× to 11× for blood and 36× to 43× for cell lines. Additionally, we included Element AVITI Biosciences (Element) WGS data (88×–105×, derived from LCLs) to improve error correction and polishing.

**Figure 1.**
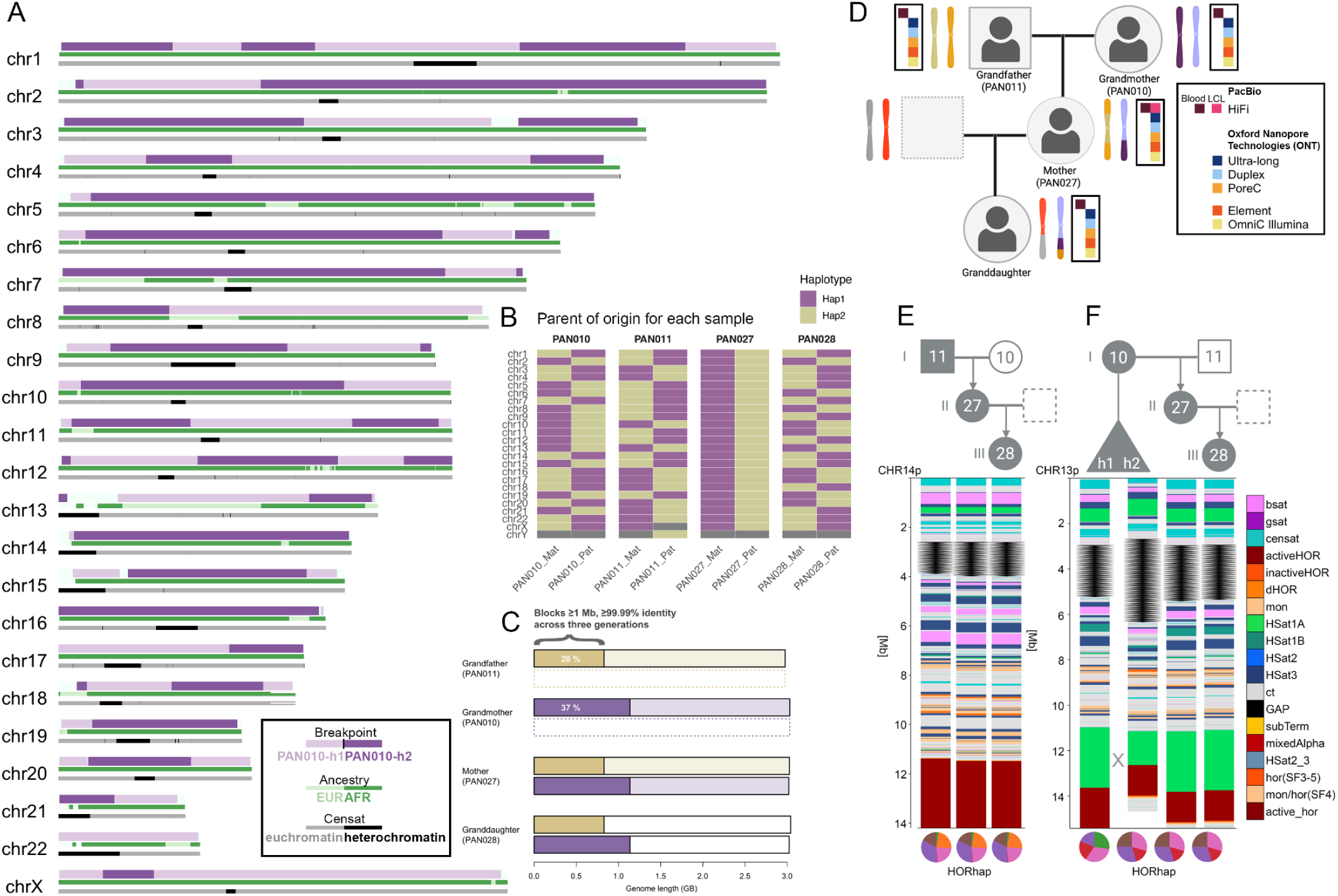
The data overview and the simplified workflow of the assembly process and patching. Maternal assembly. (A) The complete annotated maternal haplotype of the Mother (PAN027). The first layer (top) represents meiotic recombination breakpoints, switching between the two haplotypes of the Grandmother (PAN010), represented by alternating dark and light purple color. The second layer represents local ancestry inference, with dark green color representing genetic similarity to 1KGP African reference samples and light green color representing genetic similarity to 1KGP European samples. The bottom layer represents the satellite annotation within the chromosome, with the centromeric and satellite (cenSat) annotation in black. Unavailable information is represented by the dark white background. **Parent of origin**. (B) We derived the parent of origin for each of the chromosomes in each assembly using imprinted loci, including those scaffolded with HiC (PAN010, PAN011, PAN028). The plot represents the initial chromosome assortment. **Transmitted blocks.** (C) The proportion of the assembly simultaneously transmitted in highly accurate and highly contiguous blocks across the three generations. **Available datasets.** (D) The schematics of the pedigree showing family relationships, together with the available sequencing data types. **Inter-generational changes on acrocentric chromosomes.** Black arrows represent the copy number of rDNA arrays, and the HORhap pie charts show the composition of the individual centromeres based on the “centromeric haplotypes” derived from the structural variation within alpha satellites. (E) Chromosome 14, showing no gross rearrangements across the generations. (F) Meiotic recombination breakpoint as detected on chromosome 13, swapping a large portion of the satellite-rich acrocentric arm between homologous chromosomes.

Assemblies were generated using high-accuracy PacBio HiFi and ultra-long ONT reads, phased with Hi-C, and assigned chromosomes to maternal or paternal haplotypes using either parental k-mers (for PAN027) or imprinted, differentially methylated genes (Figure 1B, Supplemental Note 1 “Assembly generation”, and Supplementary Figure 1 “Parent of origin analysis”). Primary assemblies were produced with both Verkko (Rautiainen et al. 2023; Antipov et al. 2025) and hifiasm (UL) (Cheng et al. 2021), and we further explored alternative data types (duplex, R10.4.1/E8.2.1 super accuracy (Q27), HERRO-corrected reads) and assembly strategies (Supplemental Note 1 “Assembly generation”). Across datasets, ∼32 chromosomes assembled T2T in Verkko-based assemblies and up to ∼39 in duplex-based assemblies directly from the assembler, with the remainder typically breaking into 2–3 large scaffolds. While both Verkko and hifiasm (UL) were able to produce near-complete T2T assemblies from the same sequence datasets, chromosomes frequently broke in different locations depending on the assembler, reflecting methodological differences in handling complex or repetitive sequence. To systematically resolve these inconsistencies, we first established a carefully curated, manually corrected set of sites (Supplementary Table 2), and then used this benchmark to guide the development of *panpatch*, an automated, graph-based tool for pangenome-aware patching of near-T2T scaffolds into gapless chromosome-scale assemblies (Supplemental Note 2: Panpatch, Supplementary Figure 2, Supplementary Table 3). We benchmarked *panpatch* performance in PAN027 against a high-quality set of manually curated sites, confirming its accuracy in resolving challenging loci. Running *panpatch* as the last step, we generated four diploid genomes comprising 46 resolved chromosomes, including complete acrocentric p-arms (Supplementary Table 3), which were consistently transmitted across generations (as shown for chromosome 14, Figure 1EF). To evaluate scalability beyond the pedigree, we also applied *panpatch* to nine 1000 Genomes samples from the Human Pangenome Reference Consortium (HPRC, (Wang et al. 2022)), where both Verkko and HPRC Release 2 (hifiasm UL) assemblies were available, demonstrating efficiency across diverse samples (Supplemental Note 2: Panpatch, Supplementary Figure 2).

High-confidence variant calls from samples from African (AFR) and European (EUR) populations from the 1000 Genomes Project (see Methods) (1000 Genomes Project Consortium et al. 2015; Byrska-Bishop et al. 2022), aligned to CHM13 v2.0, were used as references for local ancestry inference. The majority of haplotypes in all individuals exhibited greatest similarity to 1KGP AFR reference samples: 91.4% AFR for the grandmother (PAN010), 61.7% AFR for the grandfather (PAN011), 79.7% AFR for the mother (PAN027), and 82.4% AFR for the granddaughter (PAN028). Notably, the mother exhibited the most difference in ancestry composition of her maternal and paternal complement, with ∼67% of her paternal haplotypes versus ∼94% of her maternal haplotypes exhibiting maximum similarity to 1KGP AFR reference samples (Supplementary Table 4 “Ancestry Table”). Local ancestry visualization across autosomes (Figure 1A, MGISTL-PAN027 or “PAN027” maternal) revealed distinct ancestry blocks that trace inheritance through the multi-generational pedigree. This pedigree provides high-quality genomic data with matched cell line resources for benchmarking analysis methods while contributing African ancestry haplotypes to reference genome databases.

We comprehensively annotated the genomes of all four family members using the Comparative Annotation Toolkit (CAT2.0) (improving on (Fiddes et al. 2018)), which integrates *transMap*, *liftoff*, and *miniprot* modules to project the JHU RefSeq v110 + Liftoff v5.2 gene annotation set onto each assembly using CHM13 as the reference. This approach confirmed the presence of the full complement of expected human genes, including both protein-coding and pseudogenes, and allowed us to assess transcript models, frameshifts, and premature stop codons across haplotypes. Gene counts across family assemblies were highly consistent with CHM13, with each haplotype containing ∼57,500–58,000 genes and ∼175,000 transcripts, including ∼20,000 protein-coding genes and ∼17,000 pseudogenes (Supplementary Table 5: Gene Annotations). These results demonstrate both the completeness of the assemblies and their concordance with the current human reference. Repeat content, including microsatellites, satellites, tandem repeats, and segmental duplications, was systematically characterized alongside key genomic regions such as centromeres and telomeres, and integrated with genome-wide methylation profiles (see Data and Code Availability). We annotated repetitive sequences across maternal and paternal haplotypes and compared genome-wide repeat content with CHM13. Together, these results establish end-to-end, fully annotated assemblies for a multi-generational pedigree of African American individuals, providing one of the most accurate resources to date for studying inheritance, repeat-rich regions, and reference diversity.

### Pedigree-Guided Recombination Mapping and Polishing

We tested assembly accuracy through inheritance mapping, revealing crossover breakpoints and de novo changes. To identify meiotic recombination breakpoints, we combined variant-based phasing and a shared-node mapping approach (Supplementary Tables 6 and 7: Breakpoint Analysis Shared-Node and Variant Based). Variant calling with DeepVariant (Shafin et al. 2021) followed by phasing with WhatsHap (Martin et al. 2016) yielded millions of heterozygous variants, from which phase switches were used to infer candidate recombination sites. In parallel, the shared-node method traced haplotype-specific unitigs (or a maximal contiguous sequence that has no branches or alternative paths in the assembly graph) transmitted from parent to child, providing a more conservative set of breakpoint calls. Manual curation further confirmed several ambiguous sites. In the maternal haplotype (PAN027), 46 breakpoints were detected by shared-node mapping, 45 of which were independently supported by variant calling, with additional 43 variant-only candidates enriched in pericentromeric regions such as chromosome 16 (Supplementary Figure 3). In the paternal haplotype, 25 breakpoints were detected by shared-node mapping, and all but one were validated by variant calling, alongside 29 additional variant-only candidates (Supplementary Figure 3, Supplementary Table 7 “Breakpoint Analysis Variant-Based”). Notably, the paternal chromosome 3 was inherited without detectable recombination, consistent with prior reports of non-recombinant chromosomes in a four-generation pedigree (Porubsky et al. 2025). Overall, this analysis revealed high-confidence recombination breakpoints across both haplotypes (PAN027 maternal haplotype shown in Figure 1A), with conservative and variant-based methods concordant and complementary in resolving crossover positions. As anticipated, recombination breakpoints frequently aligned with local ancestry switches (Figure 1A, Supplementary Figure 3 “Meiotic recombination breakpoints in PAN027”).

High-confidence recombination breakpoints offered accurate haplotype resolution, enabling inheritance-aware polishing to further refine base-level accuracy. Assemblies were first corrected using blood-based haplotype-aware PacBio HiFi DeepVariant calls (e.g., 2,558 edits in the mother), followed by incorporation of highly accurate short reads consistent with the Mendelian inheritance from Element Biosciences, which are particularly effective at resolving homopolymer-associated errors and contributed an additional 6,231 edits (Supplementary Figure 4, Supplementary Table 8 “Polishing Edits”). These technologies were complementary: PacBio HiFi long reads (>20 kb) span regions inaccessible to short reads but retain slight biases in homopolymers and short tandem repeats, whereas Element short reads (∼150bp PE) cannot resolve large repeat structures or structural variants but achieve high accuracy in these short tandem repeat contexts. Consistent with this, overlap between the two callsets was minimal (244 edits), and 55% of HiFi calls and 95% of Element calls intersected with our microsatellite annotation track (mono- and di-nucleotide repeats of at least 10bps). After polishing, we used Merqury (Rhie et al. 2020) to report QV for PAN027 increased from Q65.64 to Q66.33 using k=31-mers and a hybrid k-mer database derived from Pacbio HiFi and Element reads (Q69.73 using 21-mers, Supplementary Table 9: “Polishing QV”). To systematically assess misassemblies, we applied Flagger (Liao et al. 2023), a read-based method that detects coverage anomalies in diploid assemblies and classifies regions as erroneous, falsely duplicated, correctly assembled (haploid), or collapsed. Flagger analysis on the polished assemblies (v1.1) confirmed high assembly accuracy, with only 0.05–0.12% collapsed regions and 0.03–0.06% duplicated regions in HiFi assemblies; ONT assemblies showed slightly higher collapse rates but similar duplication levels (Supplementary Table 10: Flagger, Supplementary Figure 5). Finally, pedigree-aware polishing, that is, using inheritance patterns to validate or reject assembly variants, allowed us to further refine accuracy in otherwise ambiguous contexts, a key advantage of this resource. Importantly, pedigree-aware polishing used inheritance patterns (Supplementary Table 11: “Transmitted blocks”) to remove variants seen only in the mother but absent in both grandparents and the granddaughter, thereby eliminating somatic artifacts and residual assembly errors and refining accuracy in otherwise ambiguous regions. This step would not influence our de novo mutation rates, as, unless stated otherwise, we define those as variants observed for the first time in the mother (PAN027), and successfully passed to the granddaughter (PAN028). In summary, we generated complete, highly accurate T2T assemblies across a three-generation pedigree, integrating deep sequencing, pedigree-aware assembly, and novel pedigree-based polishing.

The genomes show near-complete transmission fidelity across three generations. Alignment of each grandparent haplotype to the mother’s genome and each granddaughter’s haplotype back to the mother demonstrated that 74% and 73% of three-generational inherited segments were transmitted simultaneously as aligned blocks >=1 Mb in length with >=99.99% identity for maternal and paternal lineages, respectively. This enabled tracking of 1,136 Mbs of the maternal genome and 829 Mbs of the paternal genome across the three generations (Figure 1C). Gene annotations revealed limited variation in coding regions: most genes were transmitted without change, with only rare frameshifts (0.40%-0.47%) or early stop codons (0.26%-0.31%) (Supplementary Table 5 “Gene Annotations”). The magnitude of these changes aligns with expected rates of rare functional variants per gene and supports population-based estimates indicating that most genes tolerate very few loss-of-function mutations (Karczewski et al. 2020; Jónsson et al. 2017; Rahbari et al. 2016). Copy numbers of known gene families were stable across nearly all loci, with the exception of the USP17L cluster on chromosome 4p16.1 (Burrows et al. 2010), which exhibited shifts of 2–4 units between generations. Moreover, the challenging subtelomeric locus D4Z4 associated with Facioscapulohumeral Muscular Dystrophy (FSHD) (van der Maarel and Frants 2005) was stably transmitted from both grandparents to the mother (at 27 and 34 copies for the maternal and paternal lineage, respectively, see Supplementary Figure 6 “D4Z4”). A homologous D4Z4 array occurs at chromosome 10q, sharing ∼99% sequence identity with its 4q counterpart but lacking the functional DUX4 polyadenylation signal that confers FSHD pathogenicity (van Geel et al. 2002; Clapp et al. 2007; Arends et al. 2025). At this locus, transmission from the paternal grandfather (PAN011) to the mother (PAN027) involved a copy number shift: the grandfather carried 42 and 9 repeat units across his two haplotypes, whereas the mother inherited 21 copies of the 10q D4Z4 repeat from her father. These structural changes were supported by ONT-UL reads in 15/16 studied haplotypes, while posing a challenge even for HiFi sequencing. Thus, genome-wide transmission was highly stable, with structural variation limited to specific repetitive loci known for instability. Analysis of de novo mutations within transmitted blocks revealed mutation rates for combined single nucleotide variant (SNV) and insertion/deletions (INDEL) of 5.37 x10^-6^ and 9.89 x10^-6^ per base pair and generation for the maternal and paternal lineage (0.196 x10^-6^ and 0.143 x10^-6^ not present in any of the difficult regions as defined by the Genome in a Bottle (GIAB) Consortium (Dwarshuis et al. 2024)); Supplementary Figure 7-9, Supplementary Table 12 “Genome-wide Mutation Rates”), with enrichment at satellite sequences and meiotic recombination breakpoints but no evidence of de novo transposable element insertions (Supplemental Note 3). Thus, genome-wide transmission was highly stable, with variation limited to specific repetitive loci known for instability and de novo point mutations occurring at expected rates.

The accurate assembly of an acrocentric short arm is a significant technical achievement, as these regions span megabases of satellites, segmental duplications, and rDNA arrays, repetitive elements that pose substantial assembly challenges. In particular, the highly homogeneous rDNA arrays have been the most difficult to resolve in T2T assembly efforts and are often modeled or excluded. Despite the extreme structural complexity of acrocentric short arms, we observed remarkable generational stability. Across five transmissions spanning two generations, we detected only a single meiotic recombination event on the short arm of an acrocentric, occurring between haplotypes of chromosome 13 in the grandmother (Figure 1F, Supplemental Note 4 “13p recombination event”). We localized the putative meiotic recombination breakpoint to a ∼800 bp segment at the 3′ end of the HSat1A array, directly adjacent to the centromere, where the two PAN010 haplotypes are nearly identical (2 differences per 1,000 sites). Examination of adjacent higher-order repeat haplotypes (HORhaps), that is, clusters of higher-order repeats sharing characteristic sequence or structural variants that define local haplotype structure within centromeric arrays (Altemose et al. 2022; Miga and Alexandrov 2021), provided further support for a transition to the PAN010 haplotype 2. Within this terminal ∼800 bp segment of the HSat1A array, we observed a sharp increase in de novo variation, including 21 novel variants (two of which were indels). All other acrocentric regions were transmitted intact without major structural variation. Comparison of satellite annotations between inherited acrocentric short arms revealed predominantly exact transmission of arrays (Supplementary Figure 10 “Scaffolding of the rDNA distal and rDNA proximal sequences on acrocentric chromosomes”), with a notable exception in the ACRO1 satellite family. This satellite array includes a putative protein coding gene (Boots et al. 2020), on the acrocentric short arms (Supplementary Figure 11 “The most variable genes in the transmitted regions”). Copy number variation in ACRO1 was observed on chromosome 13 (loss of 8 copies between grandmother PAN010 and mother PAN027) and chromosome 15 (loss of 4 copies between mother PAN027 and granddaughter PAN028). Detailed analysis of pseudogenes on the acrocentric short arms showed stable copy number transmission with limited variation (Supplementary Table 13). These results indicate that acrocentric short arms, among the most difficult regions to assemble, are transmitted with notable stability and accuracy, underscoring the quality of this multi-generational reference resource.

### Resolving rDNA Array Organization and Activity Across Generations

Ribosomal DNA (rDNA) arrays remain among the most challenging regions of the human genome to assemble. Despite their complexity, they are biologically essential, encoding the ribosomal RNA genes required for protein synthesis and genome stability (Hori et al. 2023; Salim and Gerton 2019). To investigate rDNA copy number and transmission across the three-generation pedigree, we estimated chromosome-specific copy numbers of the ∼45kb rDNA units by integrating fluorescence in situ hybridization (FISH) signal intensities (Supplementary Figure 12), digital droplet PCR (ddPCR), and k-mer–based estimates from whole-genome sequencing. These complementary approaches produced highly consistent results (Methods; Supplementary Table 14 “rDNA size estimates”), with copy number estimates from blood and derived cell lines (PAN027) showing strong concordance. To aid chromosome-specific assignment, we additionally incorporated WaluSat, an acrocentric p-arm distal satellite present on a subset of acrocentric chromosomes (Potapova et al. 2025). Because WaluSat varies in presence and size among individuals, it provides a useful marker for distinguishing acrocentric haplotypes in copy-number assays. Our assemblies confirmed the accurate sizing of the WaluSat array, and notably, these arrays were transmitted with identical lengths across three generations (Supplementary Figure 10). Total rDNA copy number ranged from 625 units in the grandmother to 554 units in the granddaughter, consistent with previously reported genome-wide copy number (Potapova et al. 2025). Array sizes were assigned to one of four categories: near-absent (<1 unit), small (20–50 units), intermediate (50–90 units), and very large (>90 units, up to ∼132 units/∼6 Mb) Supplementary Table 14. Arrays in the range of 1-20 units not observed and therefore not classified as a separate category. Two near-absent arrays were detected, a partial array on chromosome 15 transmitted paternally and another on chromosome 21 transmitted maternally. Across all individuals, most arrays were intermediate (45%, 18/40), followed by small (25%, 10/40), very large (17.5%, 7/40), and near-absent (12.5%, 5/40). Detailed study of inheritance of copy number across the multi-generational pedigree demonstrated overall stability of inheritance across generations (r = 0.78; Figure 2A, Supplementary Table 14 “rDNA Size Estimates”). Collectively, these findings suggest that rDNA arrays exhibit intergenerational stability (in support with prior studies (Potapova et al. 2025)), with minor copy-number changes occurring predominantly in the largest arrays, where measurement uncertainty is greatest, and minimal variation observed in the smaller arrays.

**Figure 2.**
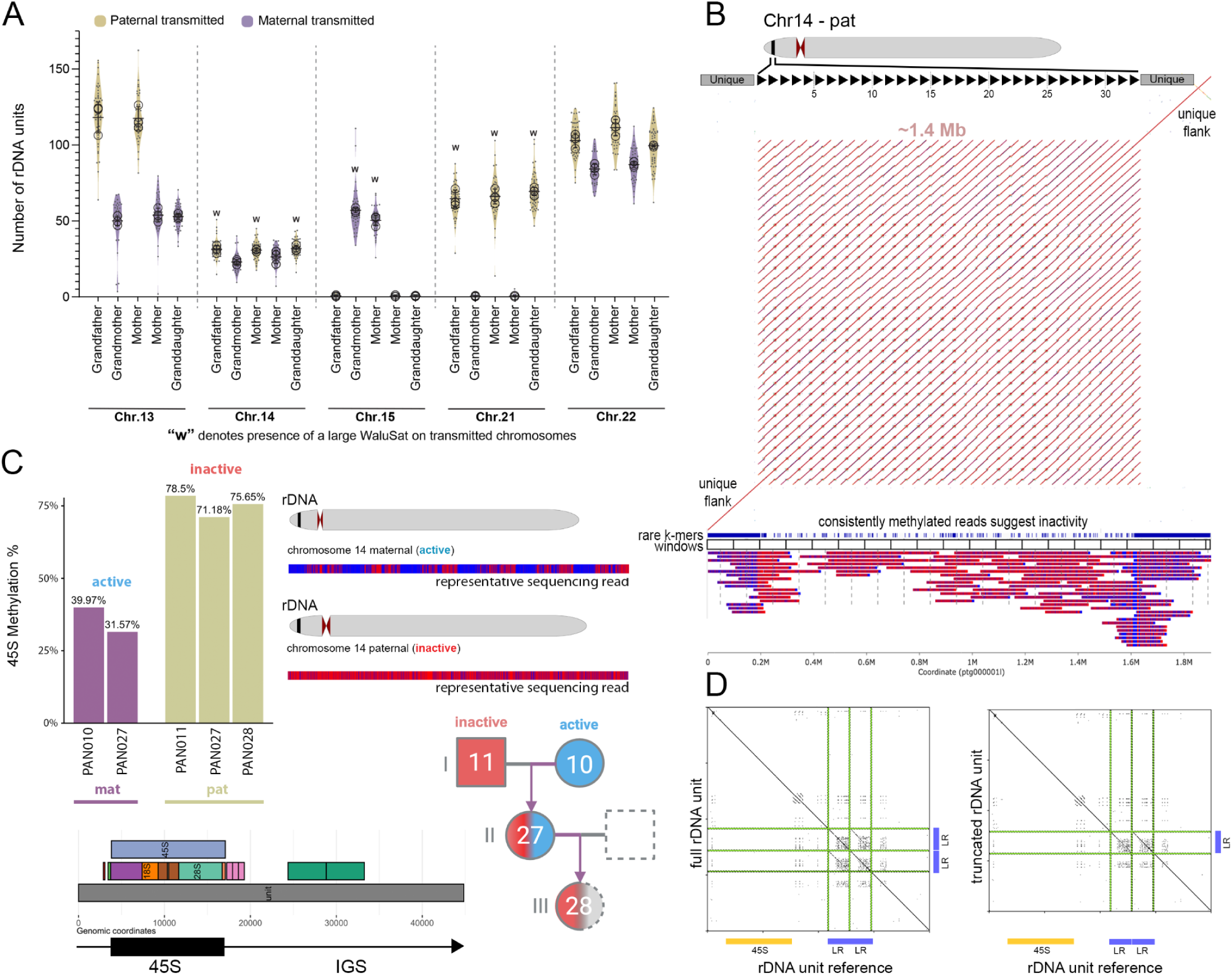
(A) The stable transmission of the rDNA arrays across generation for the paternal (gold) and maternal (purple) lineages and the five acrocentric chromosomes. The “w” denotes the presence of the walusat, a satellite array present in the subset of acrocentric chromosomes, validating the correct tracing. (B) The assembled rDNA array of the chromosome 14 paternal, spanning 33 units, with the unique flanks on each side, visualized with moddotplot. The array is methylated throughout its full length, suggesting inactivity, as visualized with modbamtools. (C) The read analysis of the rDNA reads aligned to the flanks of the original assembly estimates the 45S methylation status at 32% and 71% for the maternal (purple) and the paternal (gold) rDNA array, respectively, with consistent values across generations. (D) Structurally, paternal rDNA units on the chromosome 14 exist in two versions: full (containing all expected reference annotations, including two LR repeats), and truncated (containing all expected reference annotations, but only a single LR repeat).

While copy-number analyses offer insight into inheritance patterns, assembly provides the most direct means to examine rDNA transmission at base-pair resolution. Due to their large size and limited sequence variation, rDNA arrays require deep coverage of long, accurate reads to confidently resolve the ordering of individual repeats. To increase rDNA coverage, we generated a comprehensive dataset for the mother (PAN027) by combining ONT ultra-long whole-genome sequencing (14,922 rDNA reads, N50 = 91 kb) with adaptive sampling (Weilguny et al. 2023) (13,774 reads, N50 = 121 kb), which further enriched coverage of reads >100 kb across each rDNA region. In this work, adaptive sampling enriches for reads from repetitive, low-mappability regions by rejecting reads that are in regions known to map uniquely (determined in the assembly itself), thereby allowing reads from repetitive, low-mappability sequences to proceed through the pores for sequencing, including sequences from all rDNA arrays. This strategy targeted 12% of the genome and achieved up to threefold enrichment in rDNA coverage while capturing additional sequences from previously difficult-to-resolve genomic regions (Supplementary Table 15 “Adaptive Sampling”). The combined dataset was basecalled with Oxford Nanopore Technology’s experimental hyperbasecalling model, which improved the read accuracy and reduced indel errors compared with standard basecalling (Supplementary Table 16 “Hyperbasecalling”). The most frequent differences between the two models involved A and G substitutions and single basepair deletions of the same nucleotides, and the pairwise alignment of corresponding reads was at >99.12% identity. Compared to the super accuracy mode, the proportion of the >Q25 reads raised from 31.4% to 46.5% with hyperbasecalling (Supplementary Figure 13 “rDNA sequence quality improvements with hyperbasecalling”). This yielded the final rDNA coverage of 57x, and ∼30x in the >100kb+ category.

Having generated a high-quality, ONT-UL dataset, we assembled the rDNA-containing reads (hifiasm (ONT)) to reconstruct complete rDNA arrays from the mother’s genome. To maximize assembly accuracy, we restricted the dataset to reads exceeding a single rDNA repeat unit (≥52 kb) and set the expected number of haplotypes to eight (see Methods), corresponding to the expected number of rDNA-bearing acrocentrics after excluding the near-absent 15p-pat and 21p-mat arrays. Complete reconstruction of all rDNA arrays remains technically challenging; however, we consistently resolved the paternal chromosome 14 array, comprising 33 full rDNA units (∼1.4 Mb), together with sequences mapping to the 14p flanking regions. This assembly was consistent with experimental copy-number estimates (Supplementary Table 14 “rDNA Size Estimates”). To validate the accuracy of the assembly, we applied a stringent rare k-mer–based mapping approach (Methods) that ensured confident read support across the array, with every 100 kb window fully spanned and tiled (Figure 2BC, Methods “rDNA assembly validation”). Although we cannot claim with certainty that the paternal chr14 rDNA array is free from error, no current data or analyses suggest that it is incorrect.

The high-quality assembly of the paternal chromosome 14 array enabled detailed methylation profiling across all 33 individual 45S units, revealing uniform hypermethylation consistent with transcriptional silencing and supporting prior work linking DNA methylation to rRNA gene inactivity (Potapova et al., 2024). This result was further corroborated by a read-based analysis of full-length rDNA repeat units, which confirmed extensive methylation in the 45S region of each unit. Consistent with these findings, Potapova et al. (Potapova et al. 2025) reported a UBF-negative rDNA array on chromosome 14 transmitted from grandfather to granddaughter, where the absence of UBF binding (a factor required for transcription initiation) marks transcriptional silencing. In contrast to the fully methylated paternal chromosome 14 array, studies of the preliminary draft assembly of the maternal chromosome 14 array (four windows or ∼400kb were not confidently validated using our tiling approach) displayed reduced methylation and alternating accessibility patterns characteristic of active transcription (Figure 2C). To extend these observations, we profiled methylation of the 45S gene and its promoter in all chromosome- and haplotype-assigned rDNA reads by averaging across units from each chromosome independently (Figure 2C, Supplementary Table 17 “rDNA methylation” and Supplementary Table 18 “Assembled rDNA”). Active rDNA units were characterized by hypomethylation and periodic accessibility, whereas inactive units were hypermethylated throughout their length (Figure 2BC). The per-haplotype methylation status suggested activity or inactivity (high or low methylation in all three generations), although the profiles of sequencing reads were a more reliable indicator, as the mid-range numbers, such as those observed for chromosome 13, could potentially suggest switches in activity within an array. Still, methylation states were generally preserved across generations (Supplementary Table 17 “rDNA methylation”, Supplementary Figure 14 “rDNA methylation analysis comparing the 45S gene and the promoter”), indicating that nucleolar organizer activity can be faithfully maintained through inheritance.

To investigate intra-array sequence diversity, we compared individual rDNA repeat units within the assembled paternal and maternal chromosome 14 arrays, focusing on structural and local sequence variation. This analysis revealed structural variation at the rDNA unit scale, including a truncated Long Repeat (LR) consisting of a single copy as opposed to LR1 and LR2 present in the reference (Figure 2D). The first eight rDNA units carried this truncated version, followed by five full units, three truncated units, sixteen full units, and finally a truncated unit. For the maternal chromosome 14 array, the main structural variation was the number of TR repeats, with the majority of arrays matching the rDNA reference at 3 units, and a subset with only two TR repeats. Consistent with this, we observed a higher percentage of conserved positions in the sequence alignment of the individual units in the maternal array (92.18%), compared with the paternal array (86.73%); the combined dataset had 81.48% conserved positions. The variation was predominantly restricted to variable blocks with various di-nucleotide motifs. The median length of the blocks was only 8 basepairs and dominated by CT nucleotides (followed by CG and T), with no difference in length distribution between the maternal and the paternal arrays. However, the maternal array contained more than twice as many variable CT blocks. Interestingly, we found that G-quadruplexes, specifically those with high prediction scores, were frequently localized to variable blocks (Supplementary Figure 15 “G4s in rDNA”), suggesting that the variation in indels could potentially be modulating G-quadruplex formation.

### Centromere stability across generations

Centromeric regions consist of diverse satellite DNA classes, typically organized into megabase-scale tandem arrays that remain challenging to resolve in standard reference assemblies. As part of quality assessment, we evaluated the genomic distribution and abundance of pericentromeric and centromeric repeat families (αSat, βSat, γSat, HSat1–3) across the four pedigree assemblies (Supplementary Files “CenSat annotations”). Overall, the satellite arrays demonstrated high assembly accuracy with minimal errors and remarkable transmission stability, with only 5 of over 100 annotated arrays showing evidence of length variation across all three generations. Similarly, satellite classes were transmitted with remarkable accuracy, with sparse de novo mutation candidates in blocks transmitted across the three generations (Supplementary Table 19 “Centromere Transmissions”, Supplementary Table 20 “Centromeres QC”). We next focused specifically on the active higher-order repeat (HOR) arrays of alpha satellite DNA, which underlie kinetochore assembly. Taking advantage of the completeness of our assemblies, we evaluated the quality of each active array across three generations (23 arrays × 3 individuals = 69 total arrays). Of the 69 arrays examined, 62 showed no evidence of assembly error with Flagger Pacbio or ONT; notably, 18 of 23 arrays were transmitted without such issues across all three individuals in the pedigree. The four showed potential issues in only a single family member, consistent with overall high assembly quality (Supplementary Table 20 ‘Centromeres QC’). Ten centromeres exhibited no germline variation across several megabases of active higher-order repeats, and chromosomes 6, 10, 12, and 15 were assembled identically in all three individuals (Figure 3A), highlighting both the accuracy of reconstruction and the faithful transmission of centromeric arrays across generations. In line with our expectations from previous literature, centromeres were AT-rich and enriched for non-B DNA for a subset of chromosomes (Smeds et al. 2025) (Supplementary Figure 16 “Centromere characterization”), with Z-DNA being sporadically located in multiple locations across the active array, and A-phased repeats consistently being present either for the whole length of the array, or none.

**Figure 3.**
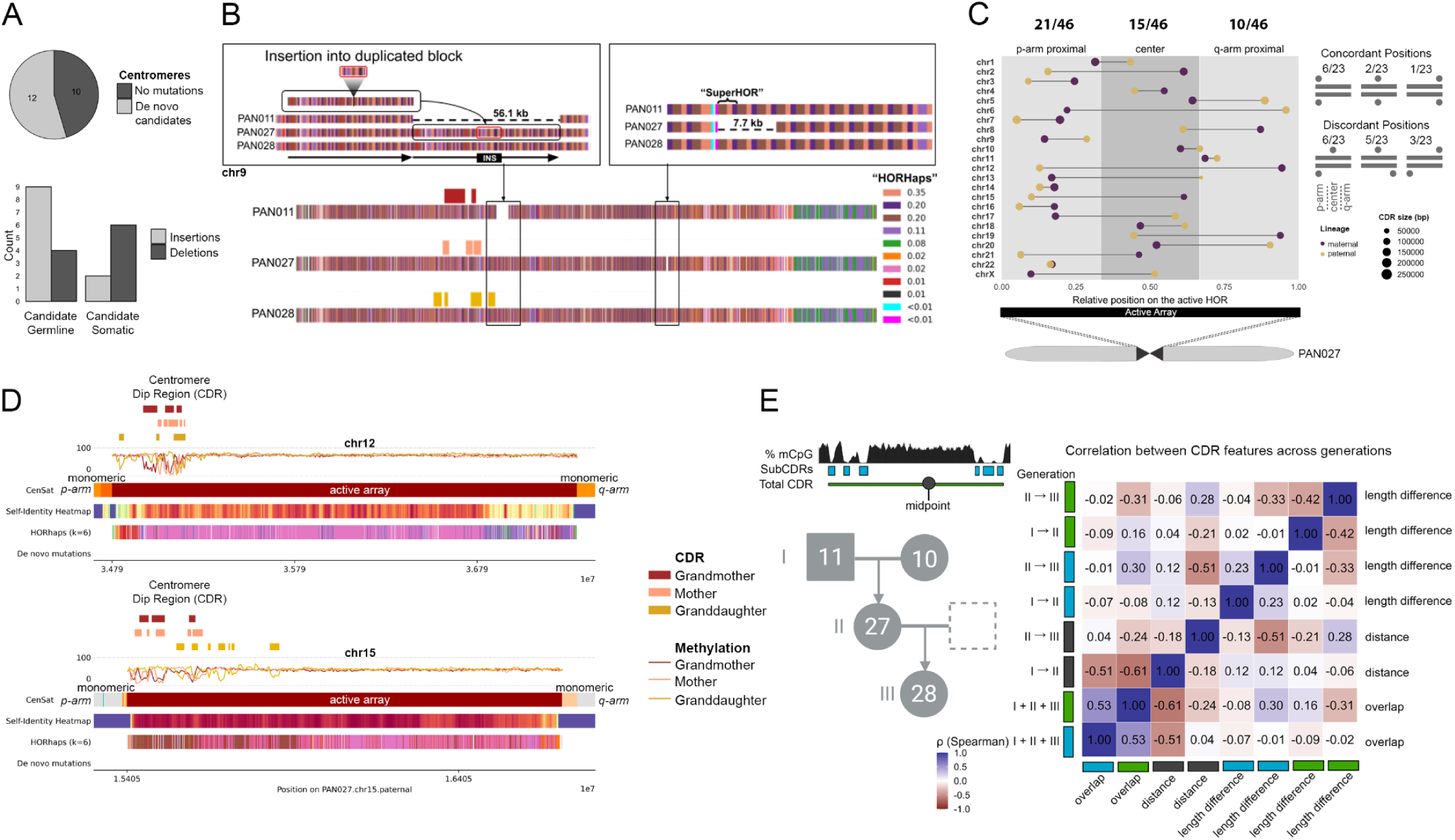
Centromeres. De novo mutations in centromeres. (A) De novo mutation candidates combining SNVs and indels were present in about half (12/22) of the studied centromeres. Longer SVs were enriched for insertions for the candidate germline mutations, and deletions for somatic variants. (B) Structural variation on the chromosome 9, with the largest observed change in the pedigree. The colored banding represents the individual HORhaps (HOR-haplotypes). The colors represent the proportion of HOR-haplotypes (HORhaps) within the PAN027 chromosome 9 active α-satellite array. (C) **Centromere Dip Region in the diploid individual.** The position of the CDR within the active array can be either p-arm proximal, central, or q-arm proximal; we display the paired CDR locations for each chromosome, and indicate the size of the CDR length (defined here as sum of subCDRs), and the parent of origin (gold for paternal and purple for maternal). We observe over twice as many p-arm locations as q-arm locations. (D) **Identical centromeric sequences across the three generations.** Chromosomes 12 and 15 represent an example of 100% identical centromeric sequences, transmitted across the three generations. The variation in the Centromere Dip Region positioning is displayed, as well as first the annotation of the monomers and active higher order repeats (top track), second, self-identity heatmap with the highest identity represented by the dark red, followed by light red, yellow, and blue (the lowest identity), third, HORhaplotypes (HORhaps) assigning each HOR copy to one of six variants. E) **Centromere Dip Region variability across the three generations.** CDR positioning was globally preserved across studied three-generational centromere transmissions. We studied the correlations between the subCDR (light blue), total CDR (green), and midpoints across the three generations (gray); studying the overlaps across the three generations, distance between the midpoints, and the total length difference.

Next, we identified single nucleotide de novo mutation candidates in all centromeres (after removing chromosome four based on QC assessments (Supplementary Table 20)), defined as variants observed for the first time in the mother, and subsequently passed to the granddaughter. This resulted in 79 candidate variants originating from 11 chromosomes (Figure 3A); interestingly these candidates had higher than expected rates of G→C transversions, despite G and C nucleotides not being prevalent in centromeres. We validated the sites of mutations using HiFi reads derived from blood, at the corresponding matching locations in each assembly. We chose this strategy to increase the accuracy of the alignment (Supplementary Table 21 “CDR Mapper Dependency”), as using this strategy, we are only aligning sample-specific reads to sample-specific references, noting that even grandparents to mother read mapping can be challenging in centromeres (Supplementary Table 22 “Centromere Validations SNVs”). The validations further supported the extraordinary faithfulness of centromere transmissions, as only four SNVs were validated (G→C, T→C, A→G, T→A) (Supplementary Figure 17 “Validations of de novo mutation candidates in the centromeres”), in line with the previous estimates (Porubsky et al. 2025), while nearly all of the remaining candidates were indicative of somatic variants.

To enable finer-scale analyses within these arrays, we developed a new method to define higher-order repeat haplotypes, or HORhaps (Supplementary Files ‘HORhap annotations’, Supplementary Table 23 “Centromere Validations indels”). HORhaps are sets of HORs within an array that share positional variants, i.e. sequence or structural differences relative to the consensus, that distinguish one group of HORs from another, thereby enabling resolution of centromeric variation at the haplotype level. This approach revealed a distinct signature for each centromere, with three-generation transmissions clearly visible, and allowed us to pinpoint precise sites of structural variation across arrays (Supplementary Table 24 “Centromeric Indels”). Variants arising in the first transmission were classified as somatic (not retained) or germline (persisting), yielding 10 somatic and 14 germline events. Somatic variants were predominantly deletions, whereas germline events were more often insertions, indicating distinct mutational profiles by origin; although not statistically significant, the odds of observing a deletion were estimated to be 6.75-fold higher in somatic than in germline events. Most germline events were modest (deletions of ∼1.3–4.9 kb; insertions of ∼1.2–7.5 kb), except for a large 56 kb insertion on chromosome 9 encompassing 69 HORs (Supplementary Figure 18 “Structural changes in chromosome 9 centromere”), characterized by a tandem duplication of 62 repeats interrupted by a 7-HOR insertion (highlighted in Figure 3B). Overall, we identified 9 maternally and 14 paternally inherited centromeres transmitted from grandparents to mother, and subsequently passed to the granddaughter, and, combining SNVs and indel de novo mutation candidates, observed 3 and 14 de novo mutation candidates in the maternal and paternal lineage, respectively (Supplementary Tables 22 and 23).

Finally, we studied changes in the Centromere Dip Region across generations. CDR is the methylation drop coinciding with the CENP-A protein binding, typically represented by multiple consecutive subCDRs separated by methylated spacers. The CDR region was generally located towards the 5’ end of the chromosome and preserved across generations, with the midpoint moving only by the median of 55 and 46 kb between generations (Supplementary Table 25 “CDR Variability”). The median overlap of CDRs across generations was 41% (calculated as intersect divided by the union of total length), while the individual subCDRs fluctuated within the total CDR span, with the median intersect of 8%. The total subCDR length was more preserved than the CDR length (median difference of 12-30 kb and 85-99 kb for the first and the second generational transmission). While our pedigree generally preserves CDR location generationally, we also observe general CDR positioning variation consistent with that reported across a panel of diverse haplotypes (Altemose et al. 2022); for example, maternal and paternal copies of the chromosome X centromere are in the 5’ distal end, and the middle, respectively.

### Telomere Length Inheritance Across Generations

Telomere length plays a critical role in age-related disease and cancer. While previous work using FISH suggested telomere lengths are inherited (Graakjaer et al. 2004, 2006), long read sequence methods now allow unprecedented accuracy in length determination (Karimian 2024). Leveraging complete assemblies across our three-generation pedigree, we reconstructed subtelomeric regions with near-perfect accuracy, including the segmental duplications that typically confound assembly (Supplementary Figure 19 “Segmental duplications”), and used the 200kb directly adjacent to the telomeric repeat for the chromosome-end tracing. Analysis of the 5 Mb terminal regions of each chromosome revealed that subtelomeric sequence organization was strikingly stable across generations, with conserved block structure and spacing. The analysis of subtelomeric regions jointly with the flagger and nucflag annotations revealed that the proportion of QC-flagged regions (Supplementary Table 26 “Subtelomeric Regions”), while higher than genome-wide estimates, were predominantly driven by the p-arms of acrocentric chromosomes, for which these algorithms have not been optimized and are likely to overcall (Supplementary Figure 5 “Flagger Quality Control”). Nonetheless, we identified evidence of allelic recombination at multiple subtelomeric loci (chr11p, chr12p, chr7q, chr11q, chr13q maternal; chr8p, chr14p chr18p, chr20p, chr4q, chr6q, chr11q, chr15q, chr18q, chr19q paternal), Supplementary Table 7 “Breakpoint Analysis Variant-Based”), consistent with rare but measurable sites of subtelomeric exchange. The stability of subtelomeric assemblies enabled confident mapping of reads for precise chromosome specific length measurement.

Accurate estimation of telomere length requires careful consideration of sequencing source and technology, which we evaluated in our pedigree datasets (Supplemental Note 5 “Telomere Length Differences in LCLs and Blood”) EBV-transformed lymphoblastoid cell lines (LCLs) are known to develop abnormal telomere profiles because most cells senesce after transformation, and surviving clones maintain growth by activating telomerase (Counter et al. 1994) or alternative lengthening of telomeres (ALT) (Bryan et al. 1995), resulting in unusually long telomeres. Consistent with this, our pedigree LCL-derived reads, whether sequenced with PacBio HiFi or ONT ultra-long, produced length estimates exceeding those typically reported for human blood (typical median 8-10 kb). In contrast, blood-derived HiFi data had telomere length within the expected range (Karimian et al. 2024) (Figure 4A, Supplementary Figure 20), was reproducible across individuals, and displayed the expected k-mer profiles. We examined telomere length at each chromosome end in all members of the pedigree using high-quality reference assemblies and deep blood-derived PacBio HiFi data (44×–70× coverage). Telomere lengths were estimated using TeloNP (Karimian et al. 2024), after assigning the corresponding reads to subtelomeric regions. Each chromosome end exhibited a distinct length distribution (Supplementary Table 27 “Telomere Lengths Per Chromosome”). The longest and shortest p- and q-arm chromosome end telomeres for all family members correlated with prior reports (Rho 0.32-0.41 and p-value 0.011-0.052) (Karimian et al. 2024) (Supplementary Figure 21). Telomere lengths across all chromosome ends with sufficient read support (≥5 reads per chromosome end) are shown in Figure 4B. These findings confirm that human telomere length is chromosome end–specific, and this haplotype-level resolution provided the foundation to directly trace telomere length transmission across generations.

**Figure 4.**
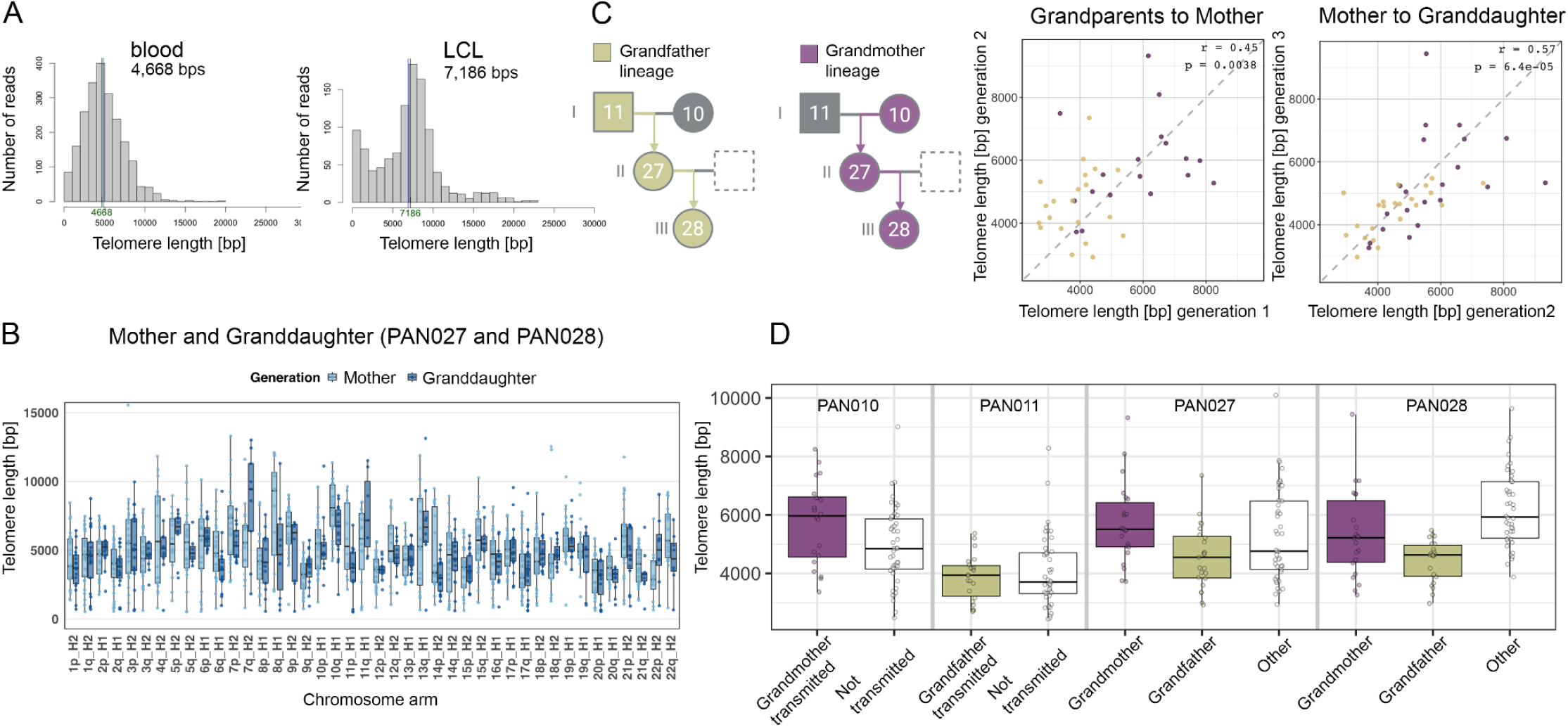
The telomeric length transmissions in the pedigree when tracing haplotypes using fully assembled subtelomeric sequences. (A) The expected telomere length distribution in the blood (PacBio) versus excessively long telomeres derived from LCL (PacBio) in the mother indicate the importance of using blood-based data for the telomere length estimation. (B) The telomere lengths for the two most recent generations (PAN027 and PAN028), and for individual chromosome arms. (C) The correlation of telomeric lengths across generations, specifically first and second generation, and second and third generation, with the telomeres originally inherited from grandmother in purple and grandfather in gold. (D) The telomere lengths of transmitted and not transmitted telomeres for the maternal lineage (purple) and the paternal lineage (gold).

Having established reliable telomere length estimates from blood-derived HiFi data, we next examined their transmission across the pedigree. Telomere lengths showed clear inheritance patterns across generations. For each chromosome in the mother, we identified its parental origin and compared telomere lengths at corresponding chromosome ends between the mother and the source grandparent. These parent-offspring comparisons revealed strong correlations (grandmother/grandfather to mother, r = 0.45), which persisted in the subsequent generation (mother to granddaughter, r = 0.57) (Figure 4C). Direct grandparent-granddaughter correlations were weaker than consecutive parent-offspring pairs, consistent with age-related telomere shortening. Critically, telomeres showed greater similarity when comparing the same inherited haplotype across generations than when comparing different haplotypes within an individual, consistent with the large difference in maternal and paternal telomere lengths for some chromosome ends previously described (Karimian 2024). For example, the shortest telomeres inherited from the grandfather remained among the shortest in both the mother and granddaughter across all three generations, despite modest length changes (Figure 4D). To test robustness, we permuted haplotype assignments and observed only a residual chromosome-specific correlation (r = 0.19; Supplementary Figure 22 “Haplotype- and chromosome-specific telomere lengths”), confirming that the observed signal reflects haplotype-driven inheritance rather than chromosome-specific length alone. Together, these findings demonstrate that telomere length is inherited in a haplotype-specific manner and that intergenerational transmission preserves relative chromosome-end ranking, even as absolute lengths change.

## Discussion

Here we present the first complete telomere-to-telomere assemblies of a multi-generational pedigree, enabling direct observation of Mendelian inheritance at base-pair resolution across entire human genomes, including previously inaccessible repetitive regions critical for cell division, chromosomal stability, and ribosome biogenesis. Analysis of inheritance across the three generations reveals remarkable transmission fidelity, with 73–74% of the transmitted genome passed as ≥1 Mb haplotype blocks showing ≥99.99% sequence identity, totaling 1,136 Mb of maternal and 829 Mb of paternal sequence. Notably, one chromosome (chr 3) was transmitted without detectable recombination, underscoring the exceptional preservation of long-range haplotypes across generations. In addition, the future work could in study non-crossover, gene conversion events. Overall, we provide scalable, automated methods that address long-standing barriers requiring manual intervention. Our graph-based assembly completion approach (*panpatch*), combined with pedigree-aware polishing that leverages Mendelian inheritance to filter artifacts, achieves Q66–Q70 consensus quality and establishes a generalizable framework for high-fidelity genome finishing. These assemblies include African American individuals, of admixed ancestry with both LCL and iPSC lines available, creating a valuable resource for integrating genomic and functional studies across diverse haplotype structures. This open-access resource enables direct examination of intergenerational stability at the genome’s most complex loci, revealing inheritance mechanisms in repeat-rich regions implicated in human disease.

The complete assembly of all acrocentric short arms represents a technical advance for studying regions that have remained largely unassembled in previous genome references. Following the transmission of the five acrocentric short arms across two generations, only a single crossover recombination event was detected, suggesting substantial stability despite megabase-scale arrays of satellites, segmental duplications, and rDNA repeats. Copy number estimates across all acrocentric rDNA arrays demonstrated stable transmission across generations (r=0.78), with array sizes ranging from near-absent sites (less than 1 unit) on chromosomes 15 and 21 to very large arrays exceeding 100 units. Achieving complete, base-level assemblies of rDNA arrays addresses one of the most persistent unresolved regions of the human genome, opening the way for mechanistic studies of their variation and inheritance. The complete assembly of an individual rDNA array on chromosome 14 (33 full units, 1.4 Mb) enabled functional characterization at base-level resolution. The observed intra-array structural variation, including truncated LR repeats, variable tandem repeat numbers, and G-quadruplex structures localized to hypervariable blocks, provide mechanistic candidates for understanding rDNA regulation and recombination susceptibility. The paternal array exhibited uniform hypermethylation across all 33 units, consistent with transcriptional silencing and corroborated by the UBF-negative status reported by Potapova et al. (Potapova et al. 2025). The maternal array displayed reduced methylation and alternating accessibility patterns characteristic of active transcription. These methylation states were maintained across three generations, indicating that nucleolar organizer activity persists through meiosis rather than resetting each generation (Potapova et al. 2025). Interestingly, the pattern of silent inheritance has so far been documented in two cases of paternal transmission (Potapova et al. 2025). Collectively, these findings establish that acrocentric short arms are transmitted with high fidelity and that both rDNA copy number and epigenetic activity states are coordinately maintained across human generations.

Active centromeric arrays showed strong transmission fidelity across three generations, with ten centromeres exhibiting no detectable germline variation across megabase-scale higher order repeat regions. Four chromosomes (6, 10, 12, and 15) were assembled identically across all transmissions. Within these arrays, we characterized groups of phylogenetically similar higher order repeats, termed HORhaps. This HOR level annotation was traceable across three generations and enabled localization of structural variants within and between arrays. In contrast, the active arrays on the homologous chromosomes that were not inherited through the pedigree showed markedly different HORhap annotations, with substantial shifts in repeat type and frequency. In some cases, such as on chromosome 13, the array composition was completely distinct (Supplementary Figure 23 “HORhap variants”). Somatic variants were mainly deletions, whereas germline events more often involved insertions. Across all arrays, only four single nucleotide germline variants were detected, consistent with prior studies (Porubsky et al. 2025), indicating highly conserved sequence inheritance and accurate assembly. Structural variation was limited, with germline insertions and deletions ranging from 1.2 to 7.5 kilobases, except for a 56 kilobase tandem duplication on chromosome 9 encompassing 69 higher order repeats. After this duplication, the CDR (corresponding to the site of CENP-A binding and kinetochore assembly) extended into the duplicated segment, although the relationship between these events remains unresolved. Overall, the CDR remained largely consistent in position across generations, with median shifts of 55 to 62 kilobases between transmissions. Individual subregions fluctuated within the broader centromere dip region span. When categorized as p-arm proximal, central, or q-arm proximal, approximately half of all CDRs were located in the p-arm proximal region (45.7%), while fewer than expected by chance were located near the q-arm (21.7%). Comparisons of homologous chromosomes within PAN027 showed that CDR positions were not consistently shared between maternal and paternal homologs. Only 9 of 23 homologous pairs localized their CDRs to the same relative region of the active array (Figure 3C). Concordant placement of CDRs in q-arm–proximal regions was particularly uncommon (1/23), and the most divergent configuration (where one homolog positioned its CDR near the p-arm and the other near the q-arm) was rarely observed (3/23). CDRs were generally consistent in size (105-135 kb, calculated as the combined span of subCDRs excluding interspersed spacing). Overall, these results reveal that centromeric arrays maintain high sequence fidelity and consistent functional domain organization across generations despite tolerating limited structural variation and CDR positional shifts.

Telomere length heritability has been documented using FISH (Graakjaer et al. 2004, 2006), but chromosome end-specific and haplotype resolved inheritance patterns have remained difficult to characterize. Complete assemblies across the pedigree provided high quality subtelomeric references that enabled telomere length measurement at individual chromosome ends through confident long read mapping. Telomere length measurements from blood derived HiFi data showed chromosome end-specific telomere length distributions consistent with prior reports (Supplementary Figure 21) (Karimian et al. 2024). Parent offspring comparisons revealed chromosome-end specific inheritance, with stronger correlations between mother and granddaughter. Parent offspring comparisons revealed haplotype specific inheritance, with correlations of r = 0.45 between grandmother or grandfather and mother, and r = 0.57 between mother and granddaughter. These correlations were substantially stronger than chromosome specific effects alone (r = 0.19 after haplotype permutation), demonstrating that telomere length transmission reflects inherited haplotype identity rather than simply chromosomal context. Notably, the shorter telomeres inherited from the grandfather remained among the shortest in both the mother and granddaughter, retaining their relative ranking across three generations despite modest length changes (Figure 4D). These findings establish that telomere length is inherited in a haplotype specific manner, with stable maintenance of relative chromosome end ranking across generational intervals despite age related telomere dynamics.

This work establishes a complete T2T pedigree spanning three generations, enabling genome-wide analysis of inheritance patterns in regions that have remained inaccessible to genomic studies for decades. While recent advances have achieved near-perfect individual genome assemblies (T2T-YAO v2.0 (Chu et al. 2025); 1002C; (Sarashetti et al. 2025)) and established benchmarks for T2T genome inference (HG002 genome benchmark; (Hansen et al. 2025)), the transmission dynamics of complete human genomes across generations have not been characterized. By directly observing Mendelian transmission of centromeres, telomeres, rDNA arrays, and acrocentric short arms at base pair resolution, we provide fundamental insights into how these essential but complex regions maintain stability across generations while tolerating specific forms of structural variation. The methods developed here, including panpatch and pedigree aware polishing strategies, offer a new framework for automating complete genomes at population scale. The inclusion of African American individuals with matched iPSC and LCL resources addresses the critical underrepresentation of diverse populations in complete reference genomes, paralleling recent population specific efforts for Indian (I002C; (Sarashetti et al. 2025)) and Han Chinese (T2T-YAO; (Chu et al. 2025)) genomes while uniquely enabling functional validation of genomic features across cellular contexts. Critically, this pedigree provides an inheritance based benchmark for the complete human genome. By establishing ground truth for variant transmission, meiotic recombination breakpoints, and epigenetic inheritance in repetitive regions, this resource enables comprehensive evaluation of phasing algorithms, structural variant callers, and methylation analysis tools across genomic contexts previously excluded from benchmarking efforts. More broadly, by demonstrating that centromeres, telomeres, rDNA arrays, and acrocentric short arms can be analyzed with quality comparable to the rest of the genome, this work provides a foundation for applying pedigree-based genomic medicine to regions implicated in chromosomal instability syndromes, repeat-associated disorders, and age-related cellular dysfunction. These complete assemblies improve variant interpretation in complex regions and demonstrate that inheritance in these regions can be reliably characterized using pedigree data.

## Resource Availability

### Lead contact

Requests for further information and resources should be directed to and will be fulfilled by the lead contacts, Karen H Miga or Monika Cechova.

### Materials Availability

All cell line data used in this study were generated from lymphoblastoid or induced pluripotent stem cell lines obtained from the NHGRI Sample Repository for Human Genetic Research at the Coriell Institute for Medical Research (HG06803, HG06804, HG06807, and HG06808). Open-access HiFi sequencing data were generated from blood samples collected from participants enrolled in a prior study approved by the Institutional Review Board of Washington University in St. Louis, MO (Protocol #201905060; see Supplemental Materials for consent documentation). During recruitment, participants were given the opportunity to self-identify their demographic background; all individuals in this cohort self-identified as African American.

### Data and Code Availability

The PAN027 v1.0 assembly is available from NCBI GenBank under the accession numbers GCA_046332035.1 (paternal) and GCA_046332005.1 (maternal), with associated BioSample SAMN33621959. The polished assemblies, and the associated code are available at https://github.com/biomonika/washu-pedigree. The sequencing data hosted in the AWS bucket are available from: https://github.com/biomonika/HPP/tree/main/T2T-Pedigree-project%20. The *panpatch* is available from https://github.com/glennhickey/panpatch. Any additional information required to reanalyze the data reported in this paper is available from the lead contact upon request.

### Competing Interests

MC is related to individuals owning Oxford Nanopore shares. SK has received travel funds to speak at events hosted by Oxford Nanopore Technologies.

### Declaration of generative AI and AI-assisted technologies in the writing process

During the preparation of this work the authors used LLMs in order to assist with specific programming tasks, and on rare occassions to improve the language of the manuscript. After using these services, the authors reviewed and edited the content as needed and take full responsibility for the content of the published article.

## Acknowledgements

We thank the participants who generously contributed samples to this work. This work was supported, in part, by NIH/NHGRI UM1HG010971 and R01HG011274 (to KHM). This work is also supported, in part, by MUNI Award in Science and Humanities StG/CoG (MUNI/SC/1916/2024). Computational resources were in part provided by the e-INFRA CZ project (ID:90254), supported by the Ministry of Education, Youth and Sports of the Czech Republic. Computational resources were in part provided by the ELIXIR-CZ project (ID:90255), part of the international ELIXIR infrastructure. Additionally, this work was supported by the Intramural Research Program of the US National Human Genome Research Institute, National Institutes of Health (SK, DA, SJS, and AMP). KHM was supported by the Searle Scholars Program. Certain commercial equipment, instruments, or materials are identified to specify adequately experimental conditions or reported results. Such identification does not imply recommendation or endorsement by the National Institute of Standards and Technology, nor does it imply that the equipment, instruments, or materials identified are necessarily the best available for the purpose. This work utilized the computational resources of the NIH HPC Biowulf cluster (http://hpc.nih.gov).

## Author Contribution

Assembly generation and polishing: MC, MM, DA, SK, AMP, Assembly characterization: RSM, SMY, RCM, MC, KHM Panpatch development and/or validation: GH, MC, Assembly QC, assembly characterization and/or variant analysis: MM, MC, TP, MA, SD, JKL, MK, CMi, EX, BP, JMZ, KHM, Cell line characterization: MWM, LS, Gene annotation and analysis: PH, HL, STS, rDNA experiments and/or analyses was performed by: TAP, MC, NP, SJS, MB, SAM, MLM, OH, JLG, Centromere analysis and/or validations: JM, FR, NP, MC, MK, JME, IAA, Telomere analysis was performed by: AR, AG, MC, Experimental or computational data generation: BM, JMVG, TH, IV, CM, SKr, SL, TWo, MC, Computational resources were sourced by: KHM, MC; KHM and MC wrote the manuscript draft. KHM and MC edited the manuscript, with the assistance of co-authors. Conceptualization was the responsibility of KHM, RSF, NOS, TW, TM, CG, MC

## Methods

### Data generation

Lymphoblastoid cell lines were obtained from the NHGRI Sample Repository for Human Genetic Research at the Coriell Institute for Medical Research (cat. ID: HG06803, HG06804, HG06807, HG0680) and cultured in RPMI-1640 media with 2 mM L-glutamine and 15% FBS at 37 °C, 5% CO2. Cells were counted by Automated Cell Counter (BioRad, TC20), washed two times with Phosphate Buffered Saline (Gibco, 10010023), placed into aliquots, flash frozen in liquid nitrogen, and stored at −80°C until extraction.

NOTES for UL: UL DNA extractions for 4 cell lines. Pellets were shipped by Coriell at a concentration of 50 million cells/pellet.

HG06803 (MGISTL_PAN010)
HG06804 (MGISTL_PAN011)
HG06807 (MGISTL_PAN027)
HG06808 (MGISTL_PAN028)

### Karyotype analysis

G-banded karyotype analysis for all LCLs was performed on 5×10^6^ cells harvested at passage 4. For iPSCs HG06803 and HG06804, karyotyping was performed at passage 16, and for HG06807 it was performed at passage 35. For all cell lines, twenty metaphase cells were counted, and a minimum of five metaphase cells were analyzed and karyotyped. Chromosome analysis was performed at a resolution of 400 bands or greater.

### Overview of the sequencing datasets

PacBio HiFi sequencing from blood was obtained for all four individuals, ranging from 44× to 70× coverage. In addition, a matched blood and cell line HiFi dataset was produced for the mother (70× and 64×, respectively), providing an opportunity to directly compare sequencing performance across sample types. Oxford Nanopore Technology (ONT) Duplex sequencing from cell lines was performed with coverage between 18× and 25×. Ultra-long nanopore datasets (ONT-UL) were generated at very high depth (169×–191× total coverage, including 70×–73× of reads ≥100 kb), providing continuity for assembly of highly repetitive regions. Chromatin conformation capture data were also produced, including Pore-C (24.8×–31.5×) and Hi-C (56×–100×) from cell lines, enabling haplotype phasing and scaffolding. Illumina short-read sequencing was performed on both blood and cell line DNA for all individuals, with coverage ranging from 7× to 11× for blood and 36× to 43× for cell lines.

### Duplex sequencing

High molecular weight DNA was extracted from approximately five million cells using the NEB Monarch HMW DNA extraction kit (T3060). 5 µg of Isolated DNA was then sheared using Diagenode Megaruptor 3, DNA fluid+ kit (E07020001) following at speed of 40. The size of sheared DNA fragments were analyzed on the Agilent Femto Pulse System using Genomic DNA 165 kb kit (FP-1002-0275). Fragment size distribution of post-sheared DNA had a peak at approximately 50kb. Small DNA fragments were removed from the sample using PacBio SRE kit (SKU 102-208-300). Library preparation was carried out using Oxford Nanopore Technologies (ONT) ligation sequencing kit V14 (SQK-LSK114). PromethION high duplex flow cells (FLO-PRO114HD) were used for sequencing on the PromethION 48 sequencer (PRO-SEQ048). Four libraries were prepared per flow cell. Flow cells were washed using ONT wash kit (EXP-WSH004) and reloaded with a fresh library every 24 hours for a total sequencing runtime of 96 hours.

### Ultra-long sequencing

High molecular weight DNA was extracted from approximately 18 million cells using the Circulomics Nanobind CBB Big DNA Kit (NB-900-001-01) and UHMW DNA Aux Kit (NB-900-101-01), following the manufacturer’s protocol (Document ID: EXT-CLU-001). Due to the high cell input, reagent volumes were doubled, and the samples were eluted in three times the recommended volume, resulting in a total elution volume of 2250 µL.

Library preparation was performed using the Circulomics UL Library Prep Kit (NB-900-601-01) and the Oxford Nanopore Ultra-Long DNA Sequencing Kit (SQK-ULK001), following the Circulomics Nanobind UL Library Prep Protocol (Document ID: LBP-ULN-001). Due to the three-fold increase in DNA input (2250 µL), the library preparation protocol was modified by scaling all reagent volumes by a factor of three. Samples were eluted in a total volume of 675 µL to accommodate three library loads across three flow cells, resulting in nine libraries per sample. All samples were sequenced on a PromethION 48 sequencer (PRO-SEQ048) using R9.4.1 flow cells (FLO-PRO002). Three libraries were loaded per flow cell, and the flow cells were washed using the ONT wash kit (EXP-WSH004) before being reloaded with fresh libraries every 24 hours. Sequencing continued for a total runtime of 72 hours.

### Adaptive sampling

High molecular weight DNA was extracted from approximately ten million cells using the NEB HMW DNA Extraction Kit for Tissues (NEB #T3060), following the ONT recommended gDNA Extraction Protocol (v114_revL_27Nov2022). Library preparation was performed using the ONT Ultra-Long DNA Sequencing Kit (SQK-ULK114), following the manufacturer’s recommended protocol, with libraries eluted in a slightly higher volume to accommodate four library loads per sample. All samples were run on the PromethION 48 sequencer (PRO-SEQ048) using R10.4.1 Flow Cells (FLO-PRO114M). We enriched for the repetitive regions by creating a depletion file with easy/unique genomic regions to be used for adaptive sampling. This file was generated using the following steps: 1) assembly gap annotations, 2) cenSat annotations, 3) annotations of subtelomeres, 4) merging of all annotations; for gaps, cenSat, and subtelomeres we added 1Mb padding to account for the potential preference of ONT reads to start in non-repetitive regions. Finally, we generated the complement of the merged annotations which created our depletion file. We utilized v1.0 assembly for PAN010, PAN027, and PAN028, and CHM13 for PAN011, to test whether sample-specific files were preferred. One flow cell was dedicated to sequencing each sample. In total, four libraries were loaded per flow cell, with the flow cells washed using the ONT wash kit (EXP-WSH004) and reloaded with fresh libraries every 24 hours, for a total runtime of 96 hours.

### Pore-C Prep, Library Prep, & Sequencing - R10.4.1 FCs

Pore-C DNA preparation was performed using five million cells for each sample following ONT’s Restriction Enzyme Pore-C v5 protocol (2nd March 2022). Libraries for the Pore-C DNA samples were prepared using the ONT Ligation Sequencing Kit (SQK-LSK114), eluted in 100 µL, and run on a single flow cell for 96 hours, with a wash and reload of fresh library every 24 hours. All samples were sequenced on a PromethION 48 sequencer (PRO-SEQ048) using R10.4.1 Flow Cells (FLO-PRO114M) and washed with ONT’s Flow Cell Wash Kit XL (EXP-WSH004-XL).

### Q27 ULTRA LONG SEQUENCING

High molecular weight DNA was extracted from approximately 6 million cells using the NEB HMW DNA Extraction Kit for Tissues (NEB #T3060), following the ONT recommended gDNA Extraction Protocol (v114_revL_27Nov2022). This protocol was performed three times to generate replicate samples for library preparation. Library preparation was carried out on all three replicate samples using the ONT Ultra-Long DNA Sequencing Kit (SQK-ULK114), which includes the new E8.2.1 adaptor for improved accuracy. The manufacturer’s recommended protocol was followed, with libraries eluted in a slightly higher volume to accommodate four library loads per sample. All samples were sequenced on the PromethION 48 sequencer (PRO-SEQ048) using R10.4.1 Flow Cells (FLO-PRO114M). Three flow cells were used in total, with four libraries loaded per flow cell. The flow cells were washed using the ONT wash kit (EXP-WSH004) and reloaded with fresh libraries every 24 hours, for a total runtime of 96 hours.

### APK - R10

The ONT protocol (T2T_9211_v114_revF_27Nov2024) was followed to generate data using the Assembly Polishing Kit (APK). DNA was extracted from approximately 5 million cells using the Puregene Cell Kit (QIAGEN, 158043) and sheared to the desired fragment length using g-TUBE™ (Covaris, 520079). Library preparation was performed using the ONT Assembly Polishing Kit (SQK-APK114) to generate a single library. Sequencing was carried out on the PromethION 48 sequencer (PRO-SEQ048) using R10.4.1 Flow Cells (FLO-PRO114M) for a continuous 72-hour run. Finally, the reads were basecalled using dorado 0.8.1 and sup@v5.0.0 model.

### HiC data

Omni-C libraries were prepared using the Dovetail™ Omni-C™ Kit (PN 21005G) following the manufacturer’s protocol with minor modifications. Chromatin was fixed in nuclei, digested in situ with DNase I, and captured on chromatin beads. Fragment ends were repaired, ligated to biotinylated bridge adapters, and proximity-ligated. After reversing crosslinks, DNA was purified, and excess external biotin was removed. Libraries were constructed using the NEB Ultra II DNA Library Prep Kit (NEB #E7103) with Illumina-compatible adapters, and biotinylated fragments were enriched on streptavidin beads. The final product was split into two indexed replicates prior to PCR to maintain complexity. Libraries were sequenced on an Illumina NovaSeq 6000 (Illumina, CA).

### PacBio Sequel II Data from the blood

DNA was extracted from blood samples using Qiagen MagAttract HMW DNA kit following company protocol. Extracted gDNA was sheared on the Diagenode Megaruptor1 targeting a fragment size with mode of 18kb-20kb. PacBio libraries were made using protocol “Procedure & Checklist-Preparing HiFi SMRTbell Libraries Using SMRTbell Express Template Prep Kit 2.0”. Libraries were size selected on Sage ELF using 0.75% agarose 1kb – 18kb cassette. PacBio polymerase binding complex was generated using Sequencing Primer v5 and Sequel II Polymerase v2.2 from the Sequel II Binding Kit 2.2. Sequencing was performed on PacBio Sequel II instrument using Sequencing plate 2.0 and SMRT Cell 8M tray. Instrument Control Software version was 10.1.0. and CCS algorithm version 10.1.0.115913 was used for CCS analysis.

### The Assembly Quality and Completeness

We validated the quality of our assembly in terms of correct k-mers, coverage, and the correspondence between the sequencing reads and the underlying assembly. For k-mer correctness, we used Merqury bundle to compare Illumina-derived k-mers of size 21 bps (that were not used for the assembly generation), and compared them with our diploid assembly.

### Assembly generation and patching

For acrocentric chromosomes, we found that the duplex assemblies provided the highest contiguity, and we therefore chose them as the starting point for the patching. We separately patched all rDNA distal and rDNA proximal parts of acrocentric chromosomes, and scaffolded them using poreC data (Supplementary Figure 10). Next, the p-arms q-arms were again scaffolded using poreC data, creating the complete acrocentric chromosomes (Supplementary Figure 10). We showed previously that HiC and poreC scaffolding yielded comparable results, while poreC had the longest reads, facilitating the mapping in the satellite-rich p-arms. Finally, we scaffolded assembled p-arms with the q-arms that contained the centromere itself, but were generated with verkko assemblies. This way, we were able to assemble and assign p-arms for all chromosomes in all individuals (Supplementary Figure 10 “Scaffolding of the rDNA distal and rDNA proximal sequences on acrocentric chromosomes”).

### Panpatch procedure

1. Construct an unclipped pangenome graph for each chromosome using ‘cactus-pangenom’ from the Minigraph-Cactus software package (Hickey et al. 2024) with the ‘--chrom-vg full’ option. This step represents the bulk of the overall runtime, taking approximately 30 minutes per sample on a cluster. The remaining steps take a few seconds on a single core, per chromosome.
2. In the case of diploid inputs, use non-overlapping 1kb windows of aligned regions in the graph to compute the alignment identity between the highest-priority assembly and all other assemblies. This procedure allows the problem to be decomposed into haplotypes which are then treated independently.
3. Anchor points are then detected by scanning the reference contig telomere-to-telomere, and finding all branching points from the highest-ranking assembly, and storing them in order along the reference.
4. A path is greedily searched along the non-reference assemblies that spans all the anchor points, greedily using the highest priority assembly path that is available. We recommend users to hardmask regions suspected of QC issues, which will favor the consideration of the next possible path accounting for the assembly priority.
5. The path will be reported as a patch if it spans the whole reference chromosome.

### Ancestry

We used flare (Browning et al. 2023) to infer the local ancestry composition of the PAN010, PAN011, PAN027, and PAN028 assemblies. We selected unrelated individuals from the African and European continental groups from the 1000 Genomes Project (the ESN, GWD, LWK, MSL, and YRI populations and the CEU, FIN, GBR, IBS, and TSI populations, respectively — excluding the ACB and ASW populations, which have a larger proportion of African and European admixture) (1000 Genomes Project Consortium et al. 2015), sequenced by the New York Genome Center (Byrska-Bishop et al. 2022), to define the reference panel (Byrska-Bishop et al. 2022). Phased SNV/indel genotypes for these individuals were generated by the T2T Consortium through alignment and variant calling against the T2T-CHM13 v2.0 reference genome (https://s3-us-west-2.amazonaws.com/human-pangenomics/index.html?prefix=T2T/CHM13/assemblies/variant <u>s/1000_Genomes_Project/chm13v2.0/</u>) (Rhie et al. 2023). We obtained a genetic map for the T2T-CHM13 v2.0 assembly from (Lalli et al. 2025) (https://zenodo.org/records/14891074). Because the Y chromosome is predominantly non-recombining and its ancestry is typically inferred via haplogroup rather than local ancestry (Rhie et al. 2023), we did not generate local ancestry calls on the Y chromosome for the one sample with an XY complement (PAN011).

For each sample assembly, we generated phased variant calls using dipcall (https://github.com/lh3/dipcall) (Li et al. 2018), which directly aligns the assembly haplotypes to the T2T-CHM13 reference. We removed variants lying outside of dipcall confident regions from local ancestry analysis. For all variable sites in the 1000 Genomes Project reference panel that were not called as variants in a sample assembly, we assumed that the sample was homozygous for the reference allele.

We ran flare with recommended parameters, first performing local ancestry inference on Chromosome 1 (chr1) and then using the chr1 demographic model for all subsequent chromosomes. For PAN010, whose chr1 demographic model failed to converge for the other chromosomes, possibly due to chr1 possessing only haplotypes that matched the 1KGP AFR reference panel, while most other chromosomes were a mix of AFR-and EUR-matching haplotypes, we inferred a demographic model separately for each chromosome. Finally, because XY individuals are hemizygous for variants on the X chromosome, we included only XX individuals in the reference panel for local ancestry inference within the X chromosome non-PAR region. We treated the X chromosome PAR 1 and PAR 2 regions as autosomal, using the chr1 demographic model and full 1000 Genomes Project reference panel.

Flare provides per-SNP estimates of the probability of African and European ancestry, formatted as a VCF. In order to represent the inferred local ancestry of each sample haplotype in the coordinates of its own assembly, rather than the coordinates of T2T-CHM13, we first lifted over these VCFs from T2T-CHM13 to each haplotype’s assembly coordinates. We then converted these per-SNP ancestry estimates into ancestry “blocks” by merging consecutive SNPs with identical ancestry proportions into larger regions.

### Meiotic recombination breakpoints

Meiotic recombination breakpoints were examined using multiple methods, as described below:

First, we built a pedigree assembly graph by pooling all HiFi and all ONT reads for the trio PAN010/PAN011/PAN027 and assembling them using Verkko (v.1.3.1). The HiFi coverage in the graph as output by Verkko was 23.57X on average, with minimum and maximum values of 1.0X and 494.93X, respectively. The mean coverage of ONT reads as output by Verkko was 74.72X, and values range from 0X to 3539.32X. The initial graph consisted of 20,563 nodes, 25,219 edges and contained 8.689 Gb of sequence data, referring to homopolymer-compressed sequence. The uncompressed assembly as output by Verkko contained 12.497 Gb of sequence. The N50 value of the unitigs was 2.02 Mb with a mean length of 0.42 Mb.

Next, we detected nodes in the graph that are transmitted across the three generations.

To do so, we first located all k-mers (k=72) within each graph node, and subsequently counted the unique k-mers - those which occurred exactly once - in the reads of all four family members (grandmother, grandfather, mother, granddaughter). This allowed for assigning the corresponding nodes of the graph to the subset of samples their sequence occurs in.

Nodes that are tagged with samples from three generations (grandmother/mother/granddaughter or grandfather/mother/granddaughter) represent sequences that have been transmitted twice to the subsequent generation. Accordingly, nodes tagged with samples from two generations (grandmother/mother or grandfather/mother) which do not occur in the granddaughter’s genome represent sequences with one transmission.

We identified 2,467 nodes with two transmissions, covering a total sequence length of 2.022 Gb. Of these, 1,546 nodes belong to the maternal haplotype (total length 1.148 Gb), indicated by their presence in the grandmother, mother, and granddaughter (PAN010, PAN027, and PAN028). Consequently, the remaining 921 nodes (total length 0.874 Gb) are part of the paternal haplotype.

Additionally, we identified 2,812 nodes with one transmission, spanning a total of 2.183 Gb. Among these, 1,485 are maternal nodes (0.949 Gb), found in the mother and daughter, while 1,327 are paternal nodes (1.234 Gb).

#### Shared-nodes approach

With the aid of the three-sample assembly graph, we detected meiotic recombination breakpoints in the assemblies. First, we aligned the previously detected shared contigs – those that were transmitted over two and three generations – to the diploid assemblies of PAN010 and PAN011 using minimap2 (v.2.26). We collected all mapped intervals from the resulting PAF files for each contig that had a unique match. We extracted the mapped regions for the two haplotypes of each parent separately and merged adjacent intervals if the gap between them was sufficiently short (< 1 Mb). Based on which parental haplotype the contigs mapped to, we colored the region accordingly (blue/green coloring in the plot) and detected meiotic recombination as switches between haplotype 1 and haplotype 2 in each parent.

#### Variant-based approach

As a complementary approach, we further detected candidates of recombination breakpoints through alignments of the parental HiFi reads to the corresponding parental haploid assembly of PAN027 via minimap2 and subsequent variant calling with DeepVariant (v1.5.0). The resulting callsets were phased with WhatsHap, using ONT reads. We filtered the variant set using bcftools, retaining only phased variant sites. The paternal callset contained 4,829,307 heterozygous variants (8,527,229 total), of which 3,354,553 were phased. The maternal callset included 4,721,957 heterozygous variants (7,936,864 total), with 3,316,209 were phased. Further filtering was applied using vcftools, where we removed less confident variant calls based on quality (--minQ 30) and depth (--max_meanDP 65). After filtering, 3,004,396 heterozygous SNVs remained in the paternal callset, and 2,960,084 in the maternal set.

From the phased variant callset, potential recombination breakpoints were identified by inspecting phase changes within phased blocks (phase sets), as output by WhatsHap. We flagged a variant site as a potential switch position if a phase change occurred within a phase set, and no additional switch appeared within a range of 100 kb, in order to prevent including 0|1|0 flip errors within a short distance, especially in regions with a low number of variants.

For the final curation of the most likely set of breakpoint positions, we included all breakpoints detected using the shared-nodes approach, all of which were additionally supported by the variant-calling approach. For the remaining variant-calling based candidates that lacked support by shared contigs, we examined each position individually. For this, the HiFi reads were haplotagged by WhatsHap haplotag, using the read alignments and the phased callset. Using IGV, we validated the candidate breakpoints by examining the variation in the two haplotagged read sets around the location. Finally, for the chr13 meiotic recombination in the grandmother, we identified all 200-bp k-mers unique to either the grandmother (PAN010) haplotype 1 or haplotype 2 array, and traced both haplotypes simultaneously in comparison to the mother (PAN027).

### Telomeres

The subtelomeric regions directly adjacent to the telomeres were mapped to each other and used to derive the telomere transmissions. We used 200kb sequences (we used short regions to reduce the likelihood of spanning the recombination breakpoints) and examined all biologically plausible transmission chains to find the reliable assignments. The HiFi reads derived from the blood were mapped (using minimap2 v2.26-r1175, settings“-y -x map-hifi --MD --eqx --cs -Y -L -a -K 10g”, and filtered with -F 2308) back to the sample-specific diploid references with trimmed telomeres, to ensure that the mapping was performed solely based on the subtelomeric sequences alone. Reads mapped within the last 20kb of chromosome ends with MAPQ>=2 were selected for telomere analysis, and their telomere lengths were estimated with TeloNP from the TeloBP package (https://github.com/GreiderLab/TeloBP). Finally, the reads with telomeres shorter than 400bp were removed from further analysis.

### Genome-wide mutation rate estimates

We defined transmitted blocks as a region in the grandparent that corresponds to a region in the mother and, in turn, to a region in the granddaughter. We then extract the associated sequences from the assemblies. Variant calling is performed with minimap2 (-cx asm5 --cs; version 2.30) and paftools.js, using the mother as the reference in all cases. This allows us to apply bcftools intersect -C to retain variants that arise between the grandparent and mother but are inherited by the granddaughter. Finally, we record the positions of these transmitted variants separately on the grandparent and maternal assemblies. Next, we filtered out potentially problematic regions. This included annotations by Flagger HiFi, Flagger ONT, Nucflag HiFi, and their 20kb flanks, as well as regions identified as potential switch errors (Supplementary Figure 24 “Switch error candidates”). Using the final dataset of variants, we calculated enrichment/depletion for specific annotations. To assess the significance of our observations (i.e. whether the variants were more or less likely to overlap specific annotations), we performed permutation tests. We calculated the mean distance between each variant (exact position for SNVs or midpoint for indels) and the nearest annotation, treating overlaps as zero. Variants were shuffled within transmitted blocks using BEDTools 2.31.1 shuffle to generate permuted datasets. Finally, we compared observed and permuted statistics to derive two-sided empirical p-values. Genomic regions excluding difficult regions from CHM13 were derived from the “notinalldifficultregions” bed unde https://ftp-trace.ncbi.nlm.nih.gov/ReferenceSamples/giab/release/genome-stratifications/v3.6/README.md, and lifted to the individual haplotypes of the pedigree family members. For liftover, we used version 482.

### Gene analysis

To evaluate the gene content in our newly generated assemblies, we first checked that we found and annotated all human genes expected to be present. Then we evaluated whether the assembled genes were assembled correctly by analyzing both the frameshift mutations and the presence of stop codons, using HG002 Q100 assembly as a control, or a “best case scenario”. To do this, we first annotated the assemblies and the HG002 Q100 assembly with the Comparative Annotation Toolkit (CAT2.0) using the transMap and liftoff modules based on the JHU RefSeqv110 + Liftoff v5.2 gene annotation set using CHM13 as the reference. Second, for these annotation sets, we identified the locations of frameshifts by iterating over the coding sequence of every transcript and looking for gaps in the alignment. If the gap length was not a multiple of 3 or if the length was longer than 30bp, the gap was determined to be a frameshift. To identify the number of nonsense mutations that would cause early stop codons in the annotation sets, we iterated through each codon in the coding sequence and looked for an early stop codon before the canonical stop codon at the end of the transcript. We used bedtools intersect to find the genes located on the acrocentric chromosome short arms.

### Annotation of the 3.3kb D4Z4 repeat

The sequence of the 3.3kb repeat was extracted from CHM13 chr4:193,538,193-193,541,504 and used to build a hidden markov model with HMMER version 3.4 with ‘hmmbuild’. The repeats were then annotated with ‘nhmmer --cpu 12 --notextw --noali --tblout file.out D4Z4rep.hmm <u>genome.fa</u>’. HMMER outfiles were converted to bed files using gawk script ‘gawk -v th=0.7 -f hmmertblout2bed.awk genome.out > genome.bed’ using script https://github.com/enigene/hmmertblout2bed/blob/master/hmmertblout2bed.awk. To determine whether the chromosome 4 and 10 macrosatellite arrays have alleles permissive to aberrant DUX4 expression we annotated the pLAM sequence, a feature necessary for the molecular pathogenesis of FSHD. The pLAM sequence was obtained from Yeetong et. al. (Yeetong et al. 2023) and annotated in the assemblies using blast BLAST 2.16.0+. Blast databases were constructed from the assemblies using ‘makeblastdb -in genome.fa-dbtype nucl -out genom’ and blastn search performed with ‘blastn -db genomeDB -outfmt 6 -query pLAM.fa-num_threads 8 -out genome.out’. Functional polyAdenylation sequences were identified by confirming the presence of the ATTAAA sequence (nonfunctional allele ATCAAA) at the 166th position of the pLAM sequence. Resulting annotations of the macrosatellite repeat and pLAM were filtered by position to exclude hits outside of the chromosome 4q and 10q subtelomeric regions.

### Polishing

To generate HiFi polishing edits on all samples, we took 40× PacBio HiFi DCv1.2 reads for each sample were aligned to the diploid assembly using minimap2 with parameters -a -x map-hifi --cs --eqx -L -Y -I8g. In order to correct read phasing in long stretches of homozygosity in these alignments, the PHARAOH pipeline (https://github.com/miramastoras/PHARAOH) (Mastoras et al. 2025) was run with default parameters, using as input 40× ONT UL >100kb HG002 reads from the HPRC aligned to each haplotype of the the Q100 v1.0.1 assembly with minimap2 and parameters ‘-a -x map-ont --cs --eqx -L -Y’. DeepVariant was run on the PHARAOH alignments with --model_type=PACBIO and docker version google/deepvariant:1.6.1. Following the procedure described in Mastoras et. al (Mastoras et al. 2025), for optimizing GQ filters based on Merqury error k-mers, we applied a GQ cutoff of 5 to the DeepVariant variant calls to obtain the final HiFi callset for all samples.

For all pedigree samples we obtained 90x Element AVITI Cloudbreak data generated using Element’s Elevate kit. The libraries were QC’d for size and concentration, diluted for sequencing,then loaded into the AVITI instrument and amplified into polonies on the flow cell surface using the Cloudbreak sequencing chemistry. For PAN027, we used a trio-based strategy modelled after the one used by the Q100 consortium (Hansen et al. 2025) for generating AVITI polishing edits on PAN027. We aligned all Element AVITI reads from PAN027, PAN011, and PAN010 to each haplotype of PAN027 with bwa mem and default parameters. Next, we ran DeepVariant on each alignment with parameters --model_type=WGS and docker version google/deepvariant:1.6.1. We used glnexus v1.4.1 with --config DeepVariant_unfiltered to merge the DeepVariant gvcfs for each haplotype. Next, we ran rtg mendelian with parameters --Xphase --all-records to phase the variant calls and exclude any variants violating mendelian consistency. We filtered the PAN027 edits by those with a GQ greater than 20, read depth (DP) greater than 20 and less than 90 (double the mean coverage). We only kept edits phased to the correct haplotype and 1-2 bp indels for the final PAN027 polishing call set. For PAN010, PAN011, and PAN028, we aligned AVITI reads for a given sample to each haplotype of its assembly, then ran DeepVariant with the same parameters as described above. We kept only homozygous DeepVariant calls for these samples due to the inability to phase the short reads without parental data. We applied the same filters as for PAN027 (GQ > 20, DP > 20 and < 90, keeping only 1-2 bp indels). To avoid mismapping of reads between PAN011 X and Y chromosomes, we merged both X and Y contigs into the haplotype 1 assembly fasta file before alignment of all reads to that assembly haplotype, then excluded any edits in the PAR regions, which were annotated by aligning PAR regions from chm13v2.0 using using wfmash 0.7.0 and parameters “--keep-low-map-id --segment-length=100 --map-pct-id=99 --no-split” to the assemblies. The final 0-based coordinates for the v 1.0 assemblies were PAN011.chrX.haplotype1:0-2386588, PAN011.chrX.haplotype1:153719008-154053196, PAN011.chrY.haplotype2:0-2427643, PAN011.chrY.haplotype2:46873975-47210280.

After producing our final HiFi and AVITI call-sets for each sample as described above, we used bcftools isec -c all to identify overlaps between the two call-sets (Supplementary Table 9 “Polishing edits”). For PAN027, only 4 calls overlapped by position between HiFi and AVITI, but contained different alleles. We inspected these manually in IGV and concluded that the AVITI edits had more read support, and therefore selected only the AVITI edits in these cases for all samples. We merged the remaining HiFi and AVITI edits into a single vcf with bcftools concat, and applied the edits to the unpolished assemblies with bcftools consensus -H2.

### Horhap

#### Annotation of AS HORs

The complete annotation pipeline included 3 steps performed sequentially: HOR monomer, HOR StV (structural variants), Horhap (HOR-Haplotypes).

First, ASat HOR monomer annotations were generated using the HumAS-HMMER tool (https://github.com/enigene/HumAS-HMMER) with the AS-HORs HMM file (https://github.com/fedorrik/HumAS-HMMER_for_AnVIL/blob/main/AS-HORs-hmmer3.4-071024.hmm), as described in Altemose et al., 2022. Each record in this annotation includes the HOR name (e.g. S2C9H1L) and the monomer number within the HOR (e.g. S2C9H1L.2).

Next, StV (structural variants of a given HOR) annotations of active AS arrays were generated from the HOR monomer annotations using stv_tool (https://github.com/fedorrik/stv), as described in Altemose et al., 2022. This tool merges consecutive monomers into continuous HOR units, such that each annotated item corresponds to an entire full-length HOR (e.g. S2C9H1L.1-7) or to a deleted variant of a HOR (e.g. S2C9H1L.1-3_6-7 would stand for deletion of monomers 4-5). Other, more complex variants (e.g. with insertions) are depicted in a similar manner.

Finally, the StV annotations were used as an input to horhap_tool (https://github.com/fedorrik/horhap_tool) to generate the horhap annotation. In this step, for one chromosome all sequences of frequent StVs of HORs (present at ≥5% frequency) from all 8 centromeres (4 assemblies * 2 homologs) were extracted using bedtools getfasta (https://doi.org/10.1093/bioinformatics/btq033) and aligned with MUSCLE v3 (https://doi.org/10.1093/nar/gkh340). If a certain chromosome had 3 frequent StV (chromosomes 2, 7, 8, 9, 15), alignment was performed by stages using merge_alignments.py script from the horhap_tool github repo. Pairwise Hamming distances between all sequences in the alignment were computed, and hierarchical clustering using Ward’s method (https://doi.org/10.1080/01621459.1963.10500845) was performed. Ward dendrogram was cut to partition HORs into 6 HORhaps. The BED file where each analyzed HOR has one of 6 HORhap assignments was produced.

A shallow tree cut would produce a simple picture featuring a small number of distinct regions dominated by one of a few HORhaps. Deeper cuts (like 6 as used in this paper) would produce a complex mosaic of HORhaps, and the HORhap maps which could be used as a proxy of nucleotide sequence to identify and align regions, etc.

HORhap annotations for chromosomes 1, 5, 19 were not generated due to the complex StV structure of the HOR array (S1C1/5/19H1L) which includes stretches of dimers (monomers 6/4_5) of various lengths (Altemose 2022). However this complex StV annotation was used instead of HORhap annotation in subsequent analysis.

#### HORhap annotation alignment

To compare HORhap annotations of transmitted centromeres (or StV annotations in case of chromosomes 1, 5, 19), a custom implementation of the Needleman–Wunsch global alignment algorithm was used (https://github.com/fedorrik/annotaligner). The alignment “alphabet” consisted of strings encoding the HOR name, its StV structure, and HORhap class ID (e.g. S2C9H1L.1-7::C3). From the resulting alignments, indels were collected into a summary table (SuplTables22), including the event length (in bp and the number of HORs), and a classification of each event as an insertion or deletion and as somatic or germline.

### Centromeres

For each chromosome haplotype of the sample, all reads aligning to the active alpha-satellite array were isolated. These reads were subsequently re-mapped to the full PAN027 assembly using minimap2 (2.30-r1287). Following realignment, reads were filtered to retain only those aligning to a single haplotype of a corresponding chromosome in PAN027. Each of the two PAN027 haplotypes individually had aggregate methylation quantified with modkit (v0.5.0) on the realigned reads. Centromere dip regions (CDRs) were identified using centrodip (v0.1.0-pre1) on those aggregated methylation profiles. For the correlation analysis, we first removed CDRs below 3 kb and then filtered out the outliers using an approach based on the DBSCAN algorithm. Two and three-generational transmission plots were constructed with matplotlib, integrating realignment-based CDRs, methylation profiles, local sequence identity, PAN027 HORhaps, and identified variants to illustrate inheritance patterns across the pedigree.

#### Local sequence identity

Centromeric sequences were used as an input to ModDotPlot (Sweeten et al. 2024) using a 5 kb window size and delta = 0. The resultant bed files, with all-vs-all identities, were processed using a custom Python script that averages the identity values for each region’s closest neighbors (default of two regions on each side). Averaged values were assigned to a color scale determined either by a fixed minimum identity threshold (90%) or by the 10th percentile of observed values, and the results were written to BED files.

### rDNA

#### Cell culture, chromosome spreads, and Fluorescent In-Situ Hybridization (FISH)

Human lymphoblastoid cell lines (LCL) HG06803, HG06804, HG06807, HG0680 were obtained from Coriell and cultured in RPMI 1640 (Gibco) with L-glutamine supplemented with 15% fetal bovine serum (FBS) in a 37°C incubator with 5% CO_2_. For the preparation of chromosome spreads, cells were blocked in mitosis by the addition of Karyomax colcemid solution (0.1 µg/ml, Life Technologies) for 6-7h. Collected cells were incubated in hypotonic 0.4% KCl solution for 12 min and prefixed by addition of methanol:acetic acid (3:1) fixative solution (1% total volume). Pre-fixed cells were spun down and then fixed in Methanol:Acetic acid (3:1). Chromosome spreads were dropped on a glass slide and incubated at 65°C overnight. Before hybridization, slides were treated with 0.1mg/ml RNAse A (Qiagen) in 2xSSC for 45 minutes at 37°C and dehydrated in a 70%, 80%, and 100% ethanol series for 2 minutes each. Slides were denatured in 70% deionized formamide/2X SSC solution pre-heated to 72°C for 1.5 min. Denaturation was stopped by immersing slides in 70%, 80%, and 100% ethanol series chilled to −20°C. Labeled rDNA BAC probe RP11-450E20 was from Empire Genomics. Chromosome identification markers CenSat D14Z1/D22Z1 (LPE 014) and PML (LPH501-A) were from Cytocell. Chromosome 15 paint was from Applied Spectral Imaging. The oligonucleotide biotin-labeled probe for WaluSat was from IDT: 5’- /5Biosg/AGA AAG GGA TAG GAG TGA AGA ACA CAG GTC GCT GCA TTT AGA AAG GAG GCG GGG TCA GAG GAA T /3Bio/ −3’. Probes used in combinations were denatured in a hybridization buffer by heating to 80°C for 10 minutes before applying to denatured slides. Specimens were hybridized to the probes under a glass coverslip or HybriSlip hybridization cover (GRACE Biolabs) sealed with the rubber cement or Cytobond (SciGene) in a humidified chamber at 37°C for 48-72hours. After hybridization, slides were washed in 50% formamide/2X SSC 3 times for 5 minutes per wash at 45°C, then in 1x SSC solution at 45°C for 5 minutes twice and at room temperature once. For biotin detection, slides were incubated with streptavidin conjugated to Cy5 (Thermo) for 2-3 hours in PBS containing 0.1% Triton X-100 and 5% bovine serum albumin (BSA), and then washed 3 times for 5 minutes with PBS/0.1% Triton X-100. Slides were rinsed in water, air-dried in the dark, and mounted in Vectashield containing DAPI (Vector Laboratories). Wide-field images were acquired on the Nikon TiE microscope equipped with 100x objective NA 1.4 and Prime 95B sCMOS camera (Photometrics). Z-stack images were acquired on the Nikon TiE microscope equipped with 100x objective NA 1.45, Yokogawa CSU-W1 spinning disk, and Flash 4.0 sCMOS camera (Hamamatsu).

### Estimating rDNA copy number from FISH images

Image processing was performed in FIJI and Python. For chromosome identification in spreads from human cell lines, labeling centromeric satellite 14/22 and PML locus on the q-arm of chromosome 15 was sufficient to identify all rDNA-containing chromosomes, but for some experiments whole chromosome 15 paint was used instead of the PML.

For semi-automated quantification performed for chromosome spreads wide-field single Z-plane images were used. rDNA-containing acrocentric chromosomes were segmented using a Cellpose model trained on 2-channel images including the DAPI and rDNA signals. rDNA regions were also segmented using a trained Cellpose model. The chromosome segmentations and identification were examined and, if necessary, curated manually in Napari. rDNA intensities for each array were measured after subtracting the fluorescence background for the respective chromosomes, and the fraction of the total rDNA fluorescence intensity in the cell was calculated for each array. The sum of all intensities of all rDNA loci represented the total amount of rDNA per cell. The total rDNA copy number was estimated from Illumina sequencing data (see “Estimating rDNA copy number from k-mer coverage”). The fraction of the total rDNA fluorescence intensity was used as a proportion of the total rDNA copy number to determine the number of rDNA copies on specific chromosomes in each chromosome spread. At least 10 spreads were quantified for each individual.

### Estimating rDNA copy number from k-mer coverage

Ribosomal DNA copy numbers were estimated from k-mer frequencies in the Illumina PCR-free short read whole genome sequencing data from Illumina cell-line. NCBI entry KY962518.1 https://www.ncbi.nlm.nih.gov/nuccore/KY962518.1 was used as a reference sequence for human 45S rDNA. The 18S copy number served as a proxy for the greater 45S unit, as each unit contains a single 18S segment. A custom pipeline counted k-mers of size 31 from the 18S consensus in short read Illumina sequencing data and normalized it to counts of 31mers from G/C matched windows elsewhere in the rDNA containing chromosomes. The matched windows were of similar size to the 18S, and ten of these were randomly selected per rDNA-containing chromosome. Any k-mers which also occurred outside the matched windows were removed to ensure that counts were exclusively from the matched windows. k-mer sets were filtered to remove those with whole genome sequencing counts greater than three standard deviations from the mean of the set, or those which were missing entirely. Counts were divided by their genomic multiplicity. Finally, the median count from the 18S k-mers was divided by the median count of the matched windows to yield a copy number approximation. A pipeline referred to as CONKORD (version 7) was used for this process.

### Estimating rDNA methylation

To quantify rDNA methylation per chromosome, a custom pipeline was used. For each individual, ONT reads were aligned back to the assembled diploid genome (minimap v2.26), then filtered to retain the primary alignments overlapping rDNA arrays, not overlapping coordinates deemed unreliable by flagger, with MAPQ60. These reads were binned by chromosome, and then aligned to a reference rDNA unit (https://www.ncbi.nlm.nih.gov/nuccore/KY962518, minimap v2.26). Using a BED annotation of the rDNA unit and modkit (v0.3.0), methylation was calculated across the 45S gene or within just the promoter, and copy number was estimated as unit coverage / genome coverage. In order to analyze the methylation on the fully assembled chr14 paternal array, we utilized high accuracy hyperbasecalled sequencing reads and supplemented them with the methylation calls from the super accuracy model, requiring the relative length difference between two sequences to be less than 10%.

### rDNA assembly

We performed a targeted rDNA assembly using pre-filtered rDNA reads. To do this, we first extracted rDNA reads by using an array consisting of 10 rDNA units as our probe/reference, and hyperbasecalled reads as query. We then kept reads with alignment scores greater, or equal than 3000. Finally, we filtered reads to only keep those with a minimal length of 52kb, set haplotype number to 8 (--n-hap), and assembled paternal and maternal chromosome 14 using hifiasm version 0.25.0-r726 and r835, respectively. In order to extract the individual rDNA units from the assembled rDNA array, we built an HMM model using HMMER 3.4 and the first 3824 basepairs of the KY962518-ROT reference file. All nucleotides between two consecutive starts were considered to be a part of the one unit.

### rDNA assembly validation

We partitioned the chr14 paternal rDNA array into 100kb windows, and required each window to be spanned by high-confidence reads. A read was considered confident if it had exact match to the 80% of rare k-mers in that window, while rare k-mers were defined as k-mers of size 31, with a multiplicity less than 10.

## Acknowledgments

We would like to thank Nancy Hansen and Arang Rhie for their helpful suggestions regarding assembly quality, satellite analysis, and acrocentric chromosomes. We also thank Angelika Lahnsteiner for advice regarding mutational spectra, methylation, and nonB DNA.

## Notes

https://github.com/biomonika/washu-pedigree

