## Supplementary figures for "Complete genomes of a multi-generational pedigree to expand studies of genetic and epigenetic inheritance"


Supplementary Figure 1: Parent of origin analysis 2
Supplementary Figure 2: Assembly patching 3
Supplementary Figure 3: Meiotic recombination breakpoints in PAN027 (mother) 7
Supplementary Figure 4: Assembly polishing 8
Supplementary Figure 5: Flagger quality control 9
Supplementary Figure 6: D4Z4 11
Supplementary Figure 7: Variant calling pipeline across the three generations 13
Supplementary Figure 8: Genome-wide mutation profiles for SNVs 14
Supplementary Figure 9: Genome-wide mutation profiles for indels 17
Supplementary Figure 10: Scaffolding of the rDNA distal and rDNA proximal sequences on acrocentric chromosomes 20
Supplementary Figure 11: The most variable genes in the transmitted regions 22
Supplementary Figure 12: rDNA FISH 24
Supplementary Figure 13: rDNA sequence quality improvements with hyperbasecalling 25
Supplementary Figure 14: rDNA methylation analysis comparing the 45S gene and the promoter 30
Supplementary Figure 15: G4s in rDNA 32
Supplementary Figure 16: Centromere characterization 33
Supplementary Figure 17: Validations of de novo mutation candidates in centromeres 35
Supplementary Figure 18: Structural changes in chromosome 9 centromere 35
Supplementary Figure 19: Segmental duplications 39
Supplementary Figure 20: Pedigree telomere lengths 41
Supplementary Figure 21: Telomere length chromosome ordering 43
Supplementary Figure 22: Haplotype- and chromosome-specific telomere lengths 45
Supplementary Figure 23: HORhap variants 47
Supplementary Figure 24: Switch error candidates 48
Supplemental Note 1: Assembly generation 49
Supplemental Note 2: Panpatch 50
Supplemental Note 3: Transposable element analysis 52
Supplemental Note 4: 13p recombination event 53
Supplemental Note 5: Telomere Length Differences in LCLs and Blood 54
Bibliography 57

### Supplementary Figure 1: Parent of origin analysis

**The assemblies can be phased using imprinted regions**

We analyzed 56 canonical imprinting control regions (ICRs) in the v1.0 assemblies. PAN027 was used as a positive control, since it has trio-based parent-of-origin assignment. We also verified PAN028 using SNP-based genetic similarity, which was consistent with the ICR methylation–based assignment.

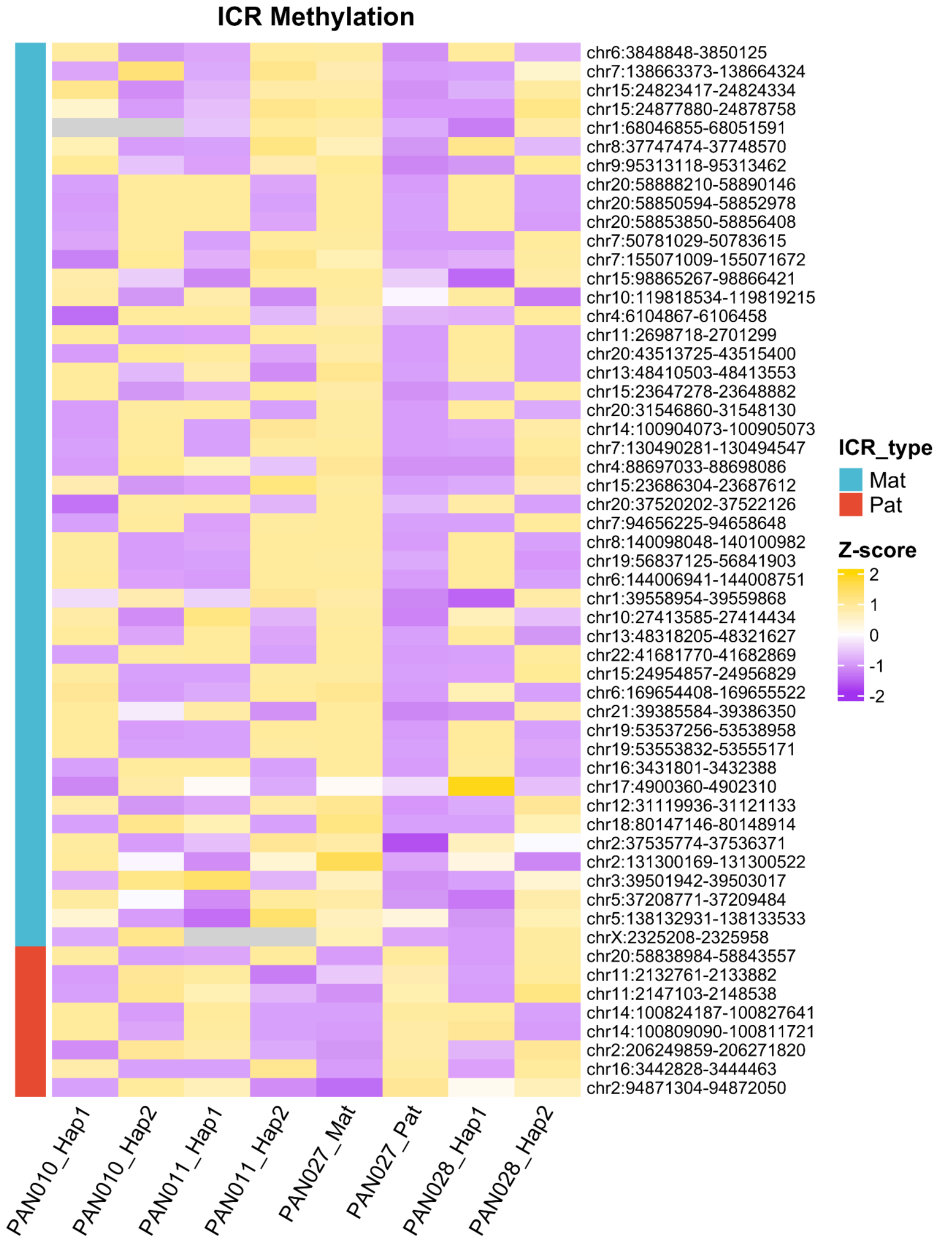


Allele-specific CpG methylation at 56 canonical imprinting control regions (ICRs) across haplotypes of four family members. Mat/Pat in the ICR_type indicates whether the region is a reported maternal imprinting region or a paternal imprinting region. Methylation values were row-standardized to z-scores using R's scale() function applied across rows. In the trio-phased PAN027 assembly, maternally and paternally imprinted regions show the expected allele-specific hyper- or hypomethylation patterns on the maternal and paternal haplotypes, validating the phasing accuracy obtained using parental k-mers. Haplotypes of PAN010, PAN01, and PAN028 show inconsistent patterns, as expected after the HiC phasing without trio-phasing.

| Similarity | PAN027 vs PAN010 | | | | | PAN027 vs PAN011 | | | | | PAN028 vs PAN027 | | | | |
|---|---|---|---|---|---|---|---|---|---|---|---|---|---|---|---|
| Chr | nSNP | PAN027Mat_PAN010Hap1 | PAN027Mat_PAN010Hap2 | PAN027Pat_PAN010Hap1 | PAN027Pat_PAN010Hap2 | nSNP | PAN027Mat_PAN011Hap1 | PAN027Mat_PAN011Hap2 | PAN027Pat_PAN011Hap1 | PAN027Pat_PAN011Hap2 | nSNP | PAN028Hap1_PAN027Mat | PAN028Hap1_PAN027Pat | PAN028Hap2_PAN027Mat | PAN028Hap2_PAN027Pat |
| chr1 | 265529 | 0.723 | 0.708 | 0.566 | 0.581 | 255821 | 0.515 | 0.639 | 0.841 | 0.594 | 263708 | 0.655 | 0.517 | 0.600 | 0.819 |
| chr2 | 280874 | 0.839 | 0.585 | 0.514 | 0.648 | 283364 | 0.549 | 0.618 | 0.777 | 0.629 | 286470 | 0.789 | 0.640 | 0.513 | 0.609 |
| chr3 | 251516 | 0.838 | 0.623 | 0.514 | 0.635 | 245476 | 0.433 | 0.757 | 0.997 | 0.430 | 242800 | 0.579 | 0.574 | 0.683 | 0.726 |
| chr4 | 252662 | 0.611 | 0.819 | 0.624 | 0.522 | 239894 | 0.687 | 0.487 | 0.481 | 0.930 | 247772 | 0.482 | 0.680 | 0.894 | 0.537 |
| chr5 | 212406 | 0.947 | 0.461 | 0.424 | 0.699 | 202980 | 0.684 | 0.477 | 0.535 | 0.872 | 203205 | 0.528 | 0.624 | 0.797 | 0.615 |
| chr6 | 217263 | 0.890 | 0.569 | 0.489 | 0.661 | 211579 | 0.709 | 0.445 | 0.418 | 0.990 | 218379 | 0.900 | 0.524 | 0.459 | 0.676 |
| chr7 | 193942 | 0.901 | 0.549 | 0.468 | 0.666 | 191161 | 0.711 | 0.447 | 0.445 | 0.959 | 193041 | 0.548 | 0.624 | 0.764 | 0.648 |
| chr8 | 187203 | 0.619 | 0.779 | 0.622 | 0.522 | 190736 | 0.685 | 0.463 | 0.529 | 0.896 | 189943 | 0.580 | 0.822 | 0.647 | 0.518 |
| chr9 | 148435 | 0.676 | 0.742 | 0.598 | 0.556 | 142668 | 0.454 | 0.709 | 0.948 | 0.489 | 142237 | 0.596 | 0.577 | 0.703 | 0.723 |
| chr10 | 173629 | 0.788 | 0.624 | 0.543 | 0.628 | 161709 | 0.457 | 0.722 | 0.957 | 0.456 | 168185 | 0.920 | 0.498 | 0.479 | 0.663 |
| chr11 | 168950 | 0.721 | 0.701 | 0.591 | 0.585 | 171128 | 0.670 | 0.481 | 0.475 | 0.916 | 171276 | 0.815 | 0.614 | 0.532 | 0.633 |
| chr12 | 153783 | 0.745 | 0.713 | 0.564 | 0.573 | 150596 | 0.466 | 0.692 | 0.919 | 0.526 | 155945 | 0.612 | 0.563 | 0.700 | 0.715 |
| chr13 | 120807 | 0.575 | 0.868 | 0.669 | 0.507 | 124872 | 0.684 | 0.485 | 0.540 | 0.891 | 118367 | 0.659 | 0.783 | 0.611 | 0.555 |
| chr14 | 111611 | 0.999 | 0.428 | 0.435 | 0.747 | 111443 | 0.591 | 0.585 | 0.696 | 0.703 | 110906 | 0.594 | 0.574 | 0.699 | 0.741 |
| chr15 | 100799 | 0.870 | 0.583 | 0.504 | 0.650 | 99897 | 0.744 | 0.432 | 0.432 | 0.981 | 99985 | 0.654 | 0.510 | 0.607 | 0.814 |
| chr16 | 106383 | 0.933 | 0.524 | 0.480 | 0.673 | 104759 | 0.725 | 0.450 | 0.380 | 0.998 | 106913 | 0.635 | 0.804 | 0.647 | 0.519 |
| chr17 | 95809 | 0.621 | 0.817 | 0.610 | 0.523 | 87920 | 0.593 | 0.549 | 0.623 | 0.754 | 93564 | 0.761 | 0.645 | 0.544 | 0.620 |
| chr18 | 92688 | 0.750 | 0.678 | 0.540 | 0.610 | 92502 | 0.680 | 0.473 | 0.523 | 0.931 | 94023 | 0.570 | 0.606 | 0.761 | 0.647 |
| chr19 | 81879 | 0.895 | 0.559 | 0.412 | 0.647 | 73723 | 0.707 | 0.455 | 0.500 | 0.877 | 77865 | 0.751 | 0.629 | 0.593 | 0.592 |
| chr20 | 81730 | 0.700 | 0.685 | 0.522 | 0.576 | 73839 | 0.421 | 0.746 | 0.993 | 0.399 | 81452 | 0.890 | 0.487 | 0.459 | 0.686 |
| chr21 | 57538 | 0.666 | 0.802 | 0.639 | 0.542 | 54599 | 0.488 | 0.742 | 0.999 | 0.452 | 57657 | 0.573 | 0.607 | 0.783 | 0.680 |
| chr22 | 49598 | 0.423 | 0.998 | 0.733 | 0.419 | 49720 | 0.759 | 0.417 | 0.439 | 0.997 | 47487 | 0.491 | 0.676 | 0.883 | 0.549 |
| chrX | 140066 | 0.500 | 0.931 | 0.695 | 0.489 | | | | | | 142836 | 0.433 | 0.689 | 0.917 | 0.529 |

SNP similarity analysis across chromosomes. In PAN027, similarity to parental haplotypes (PAN010 and PAN011) reproduced trio-based labels with complete concordance. In PAN028, haplotype similarity to PAN027 yielded parent-of-origin assignments that were fully consistent with ICR methylation, demonstrating that accurate parental origin can be resolved using either ICR methylation or genetic similarity to a single parent.

#### Supplementary Figure 2: Assembly patching

We generated three main assemblies:

“Verkko assemblies”
Using the assembler verkko 2.0, and the **HiFi**+ONT-UL+HiC data.

“Hifiasm assemblies”
Using the assembler hifiasm-ONT v0.19.8, and the **HiFi**+ONT-UL+HiC data.

“Duplex assemblies”
Using the assembler verkko 2.0, and the **HiFi/Duplex**+ONT-UL+HiC data.

The reads provided in the accurate bin are marked in bold.

**(A) The semi-manual assembly patching schematics.** First, all contigs from the primary “verkko assembly” are scaffolded using secondary assemblies (“hifiasm”, and “verkko duplex”) using a semi-manual approach. Second, the assemblies are polished using Element and HiFi sequencing reads. Third, the gaps that could not be patched using secondary assemblies were considered for patching using matching haplotype blocks from the family members.
**(B) Panpatch.** Using this automatic, graph-based method, primary assemblies are patched using secondary assemblies in the order of priority.
**(C) Assembly versioning.** The description of the individual released assembly versions.

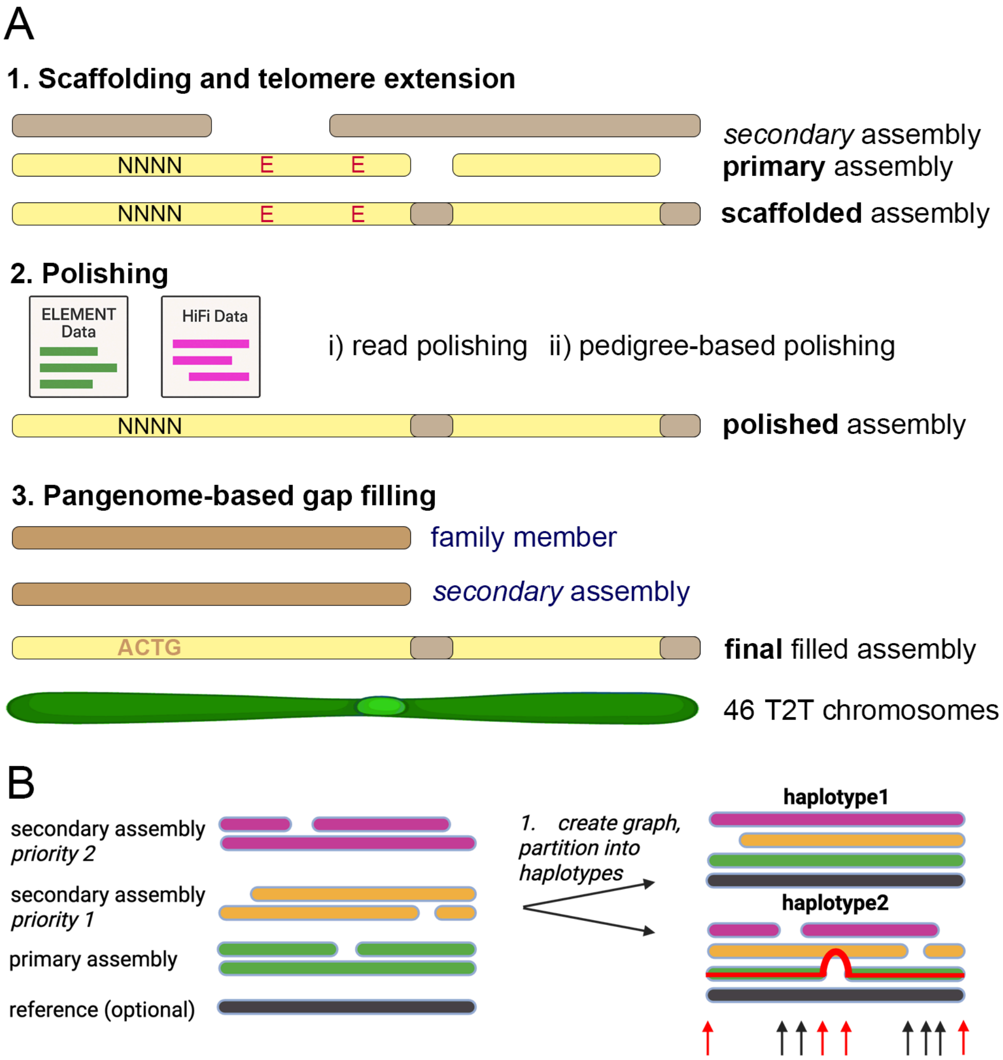

A
1. Scaffolding and telomere extension
secondary assembly
primary assembly
NNNN
E
E
NNNN
E
E
scaffolded assembly
2. Polishing
ELEMENT Data
HiFi Data
i) read polishing
ii) pedigree-based polishing
NNNN
polished assembly
3. Pangenome-based gap filling
family member
secondary assembly
ACTG
final filled assembly
46 T2T chromosomes
B
secondary assembly
priority 2
secondary assembly
priority 1
primary assembly
reference (optional)
1. create graph, partition into haplotypes
haplotype1
haplotype2

#### ASSEMBLY VERSIONING

C

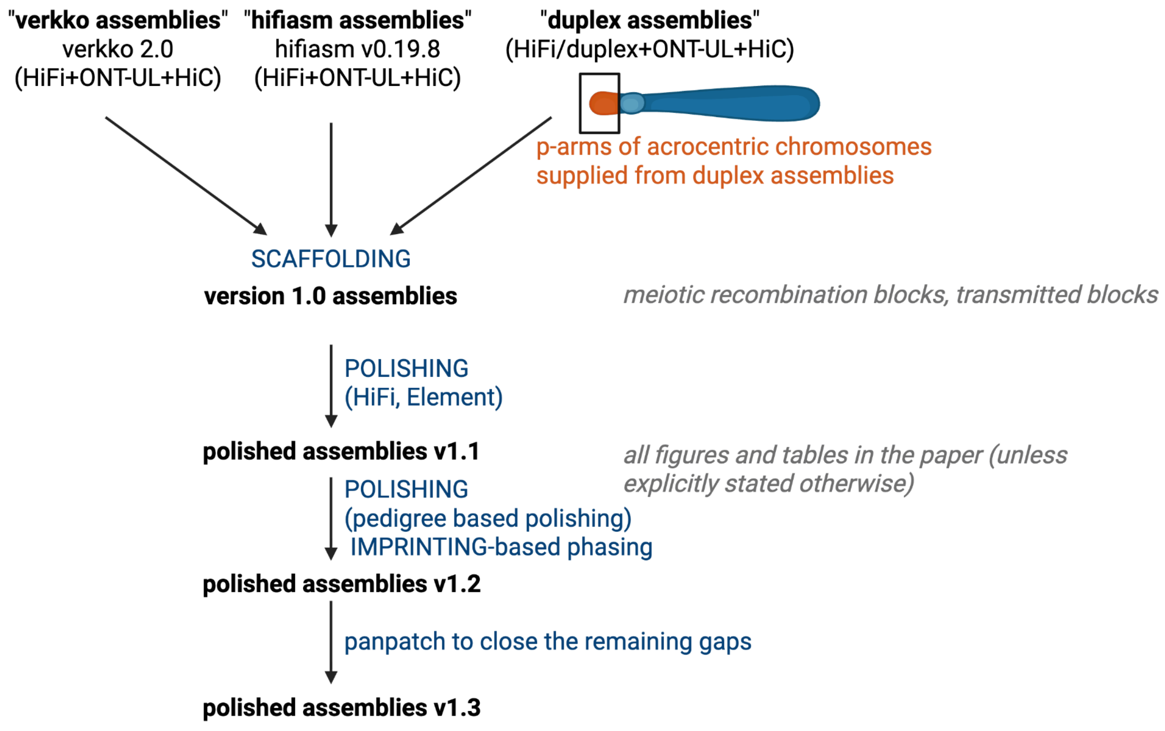

#### Supplementary Figure 3: Meiotic recombination breakpoints in PAN027 (mother)

The meiotic recombination breakpoints in the mother, PAN027 (v1.0), as reported using the Shared-nodes approach (alternating green and blue colors for the two haplotypes; Supplementary Table 5). Additionally, triangles represent the variant-based approach (Supplementary Table 6). Both maternal (left) and paternal (right) haplotypes are plotted.

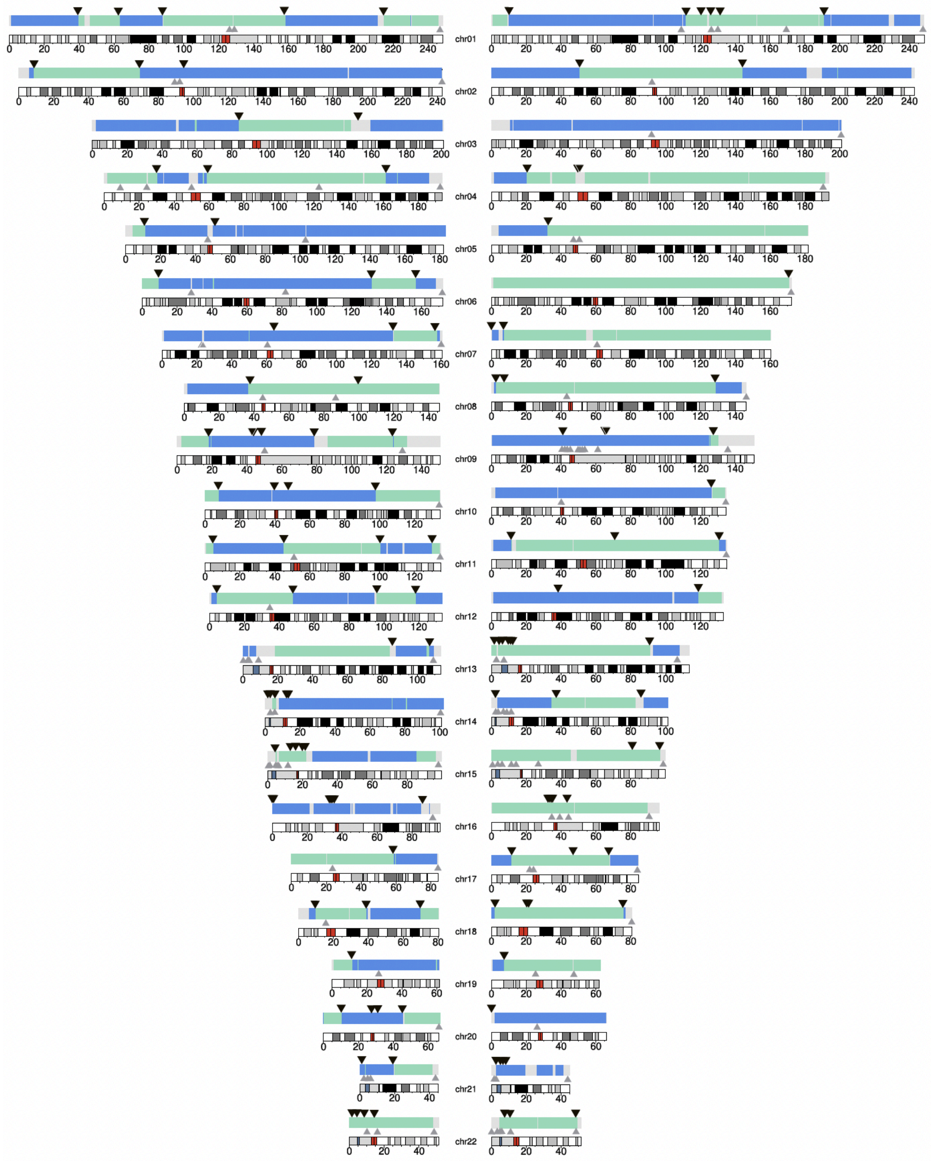

### Supplementary Figure 4: Assembly polishing

The schematics of the polishing process for the family members.

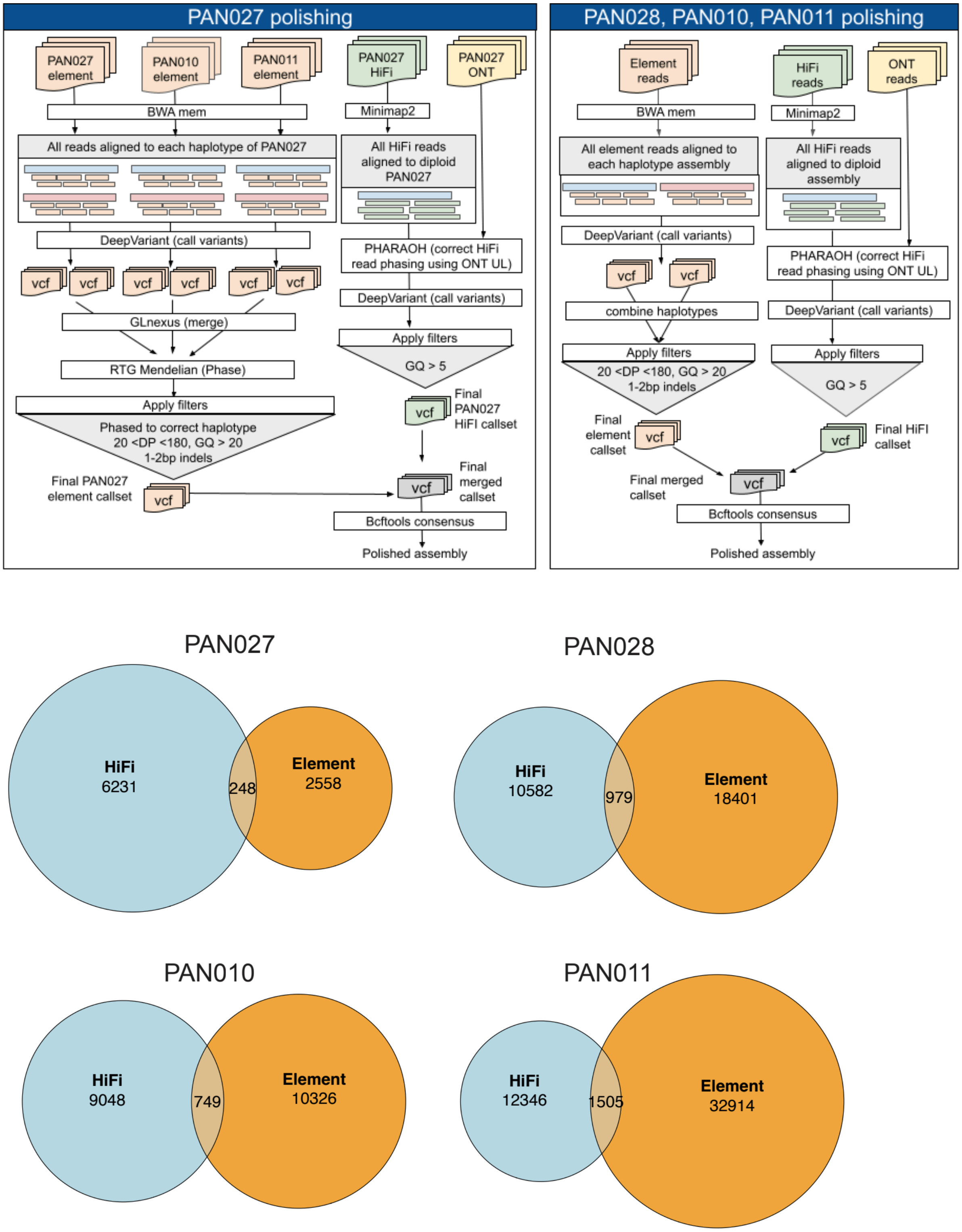

#### Supplementary Figure 5: Flagger quality control

The red color represents the flagged, potentially problematic regions in the mother PAN027 (flagger v1.2.0, conservative setting), noting that the first or the last flagged window is an artifact of the expected coverage dropout towards the assembly edges. The coordinates of the flagged regions in the rest of the family members are available as Supplementary files.

##### PAN027 Flagger HiFi:

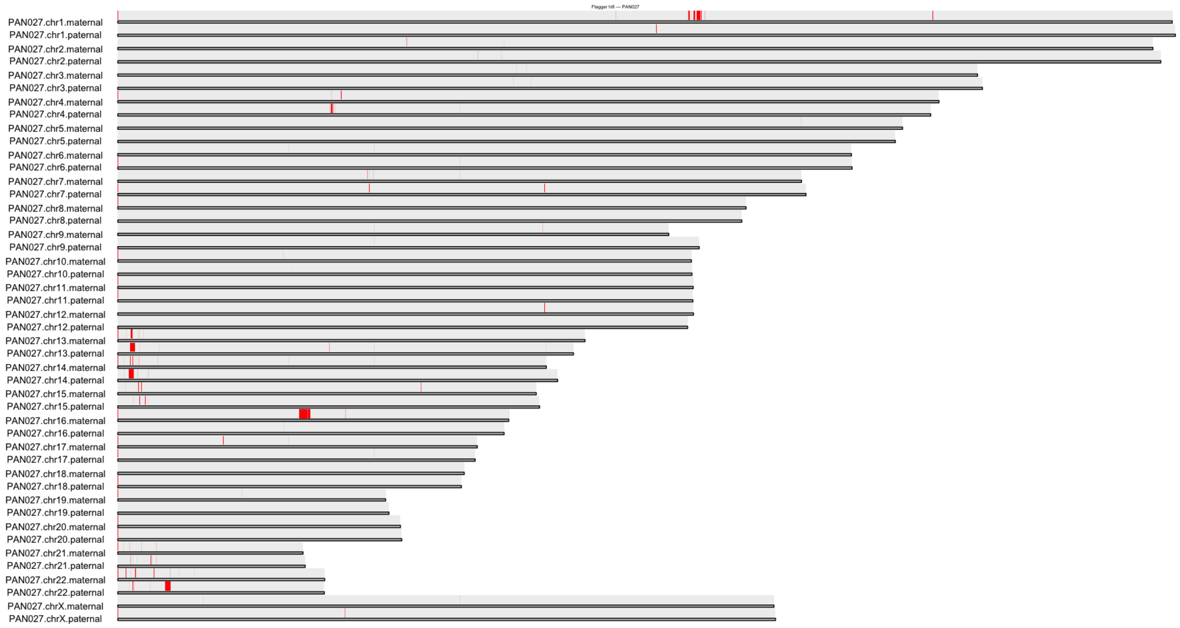


##### PAN027 Flagger ONT:

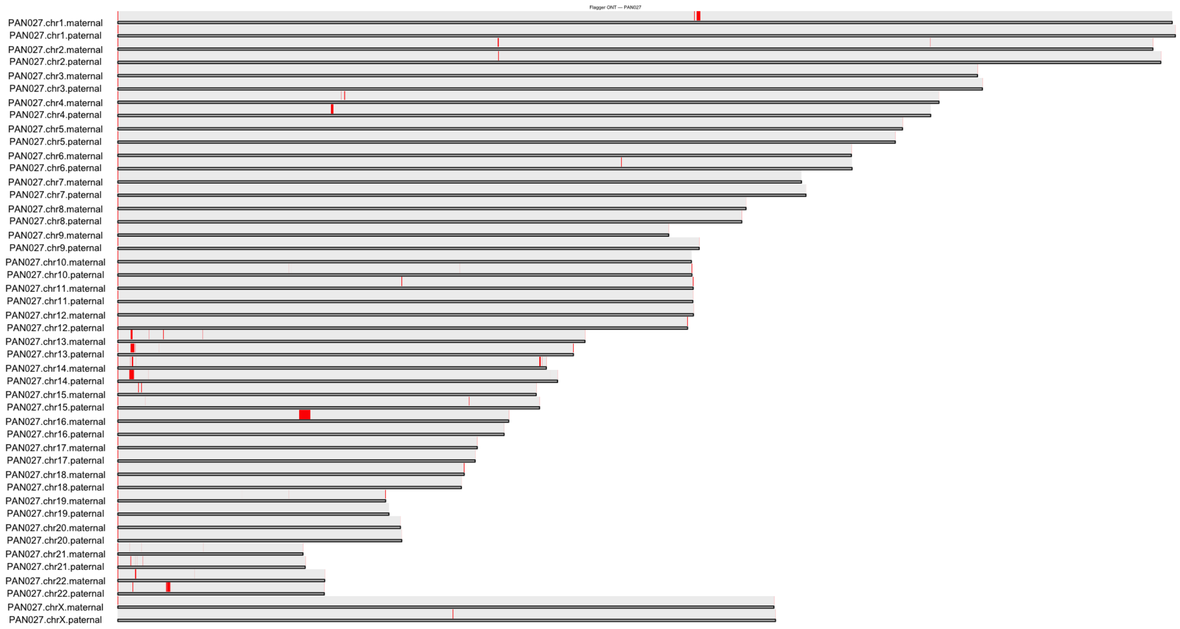

#### PAN027 Nucflag HiFi:

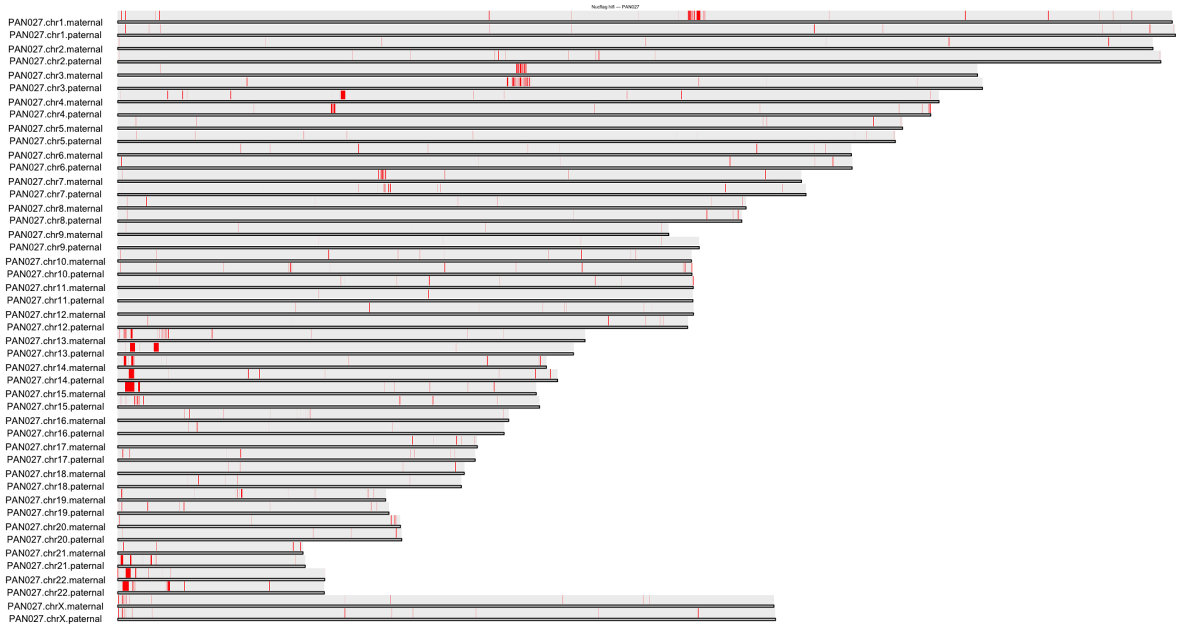


#### PAN010 Flagger HiFi:

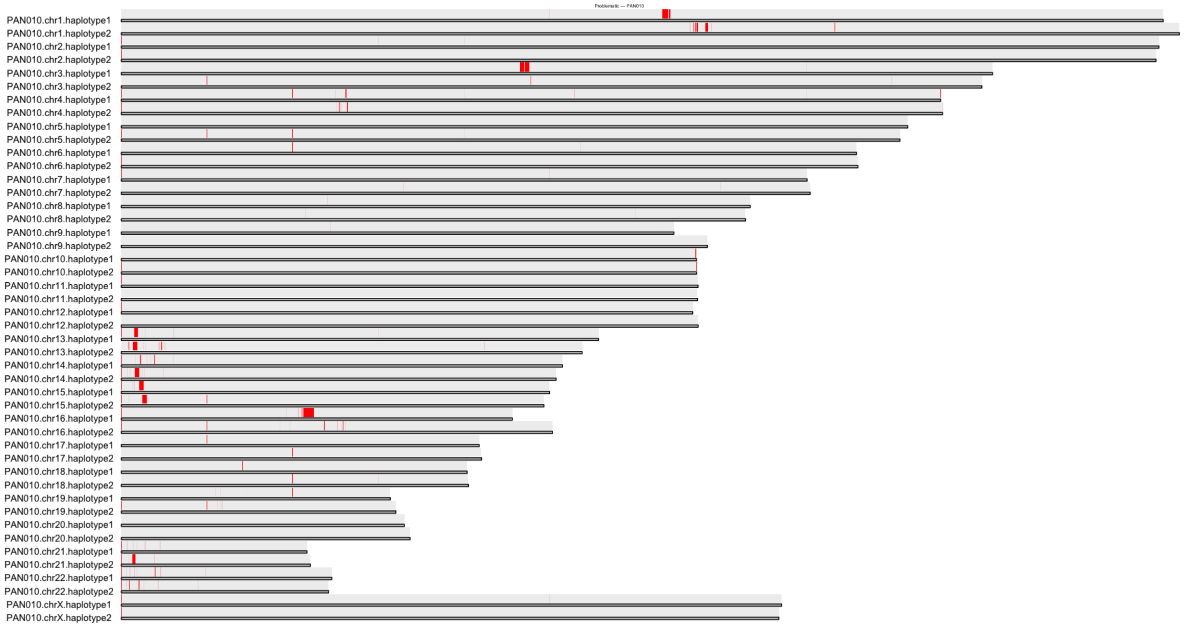

### PAN011 Flagger HiFi:

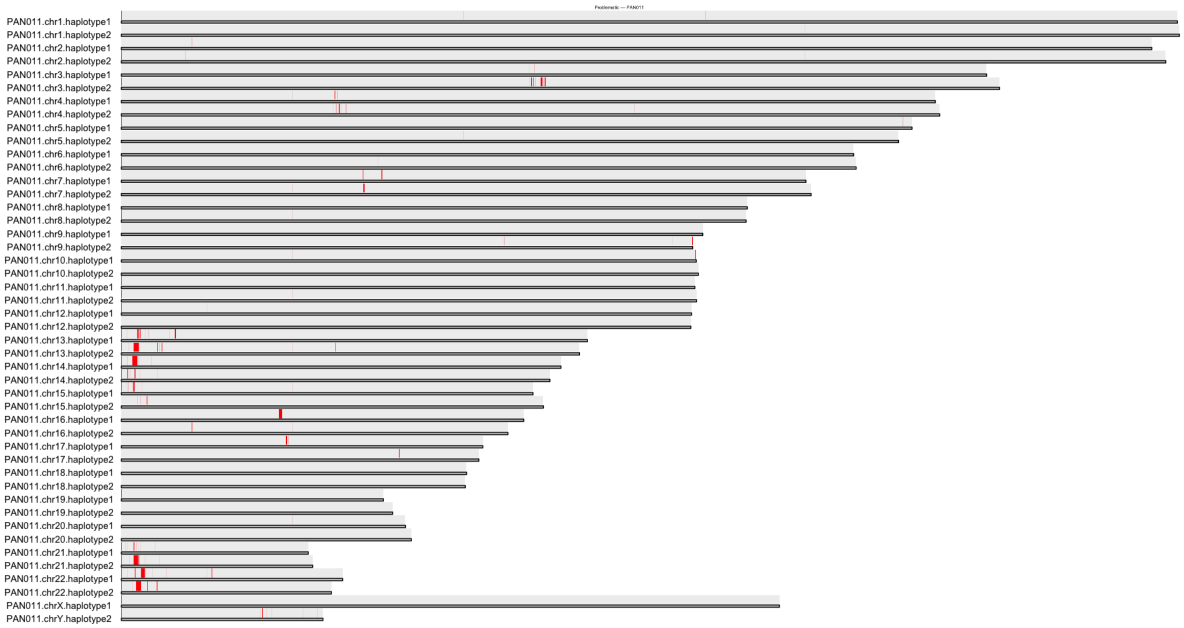


### PAN028 Flagger HiFi:

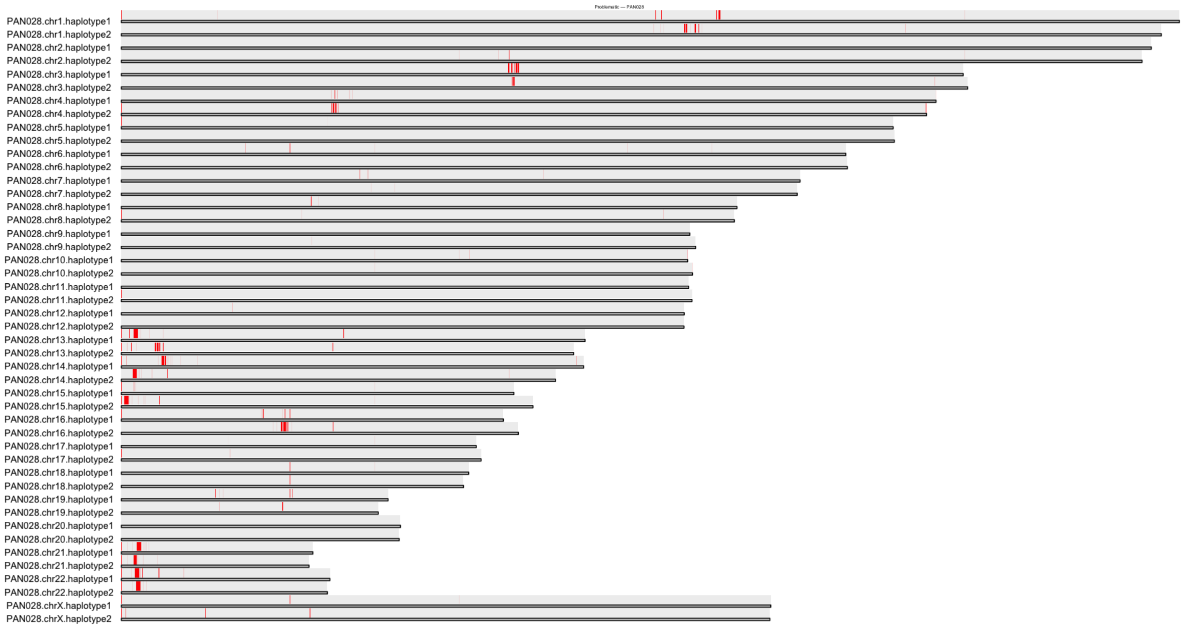

### Supplementary Figure 6: D4Z4

For the chr4 locus, the pedigree shows the stable transmission of the D4Z4 macrosatellite array between PAN011, PAN010, and PAN027. Maternally inherited haplotypes colored red and paternally inherited colored blue.

This region for chr4 is not in our three-generational transmitted blocks, so we do not have sufficient evidence to determine which version the granddaughter (PAN028) inherited from the mother (PAN027).

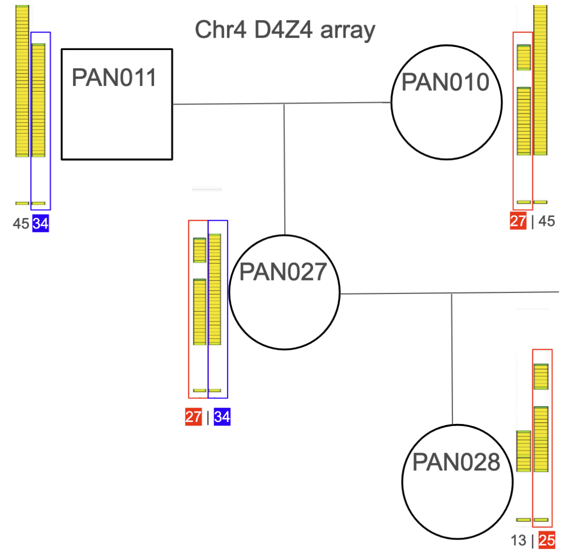


PAN010 - 27 repeats
PAN027 - 27 repeats
PAN028 - 25 repeats

PAN011 - 34 repeats
PAN027 - 34 repeats

| Sample | Haplotype | chr4 repeats | Functional polyA? |
|---|---|---|---|
| chm13 | | 35 | yes |
| PAN010 | hap1 | 27 | |
| PAN010 | hap2 | 45 | yes |
| PAN011 | hap1 | 45 | yes |
| PAN011 | hap2 | 34 | yes |
| PAN027 | mat | 27 | |
| PAN027 | pat | 34 | yes |
| PAN028 | hap1 | 13 | yes |
| PAN028 | hap2 | 25 | |

For the chr10 locus, the array inherited by PAN027 from PAN011 included a copy number change. Using sequence similarity, the most likely inheritance appears to be a deletion from 42 to 21 copies in the inherited array in PAN027.

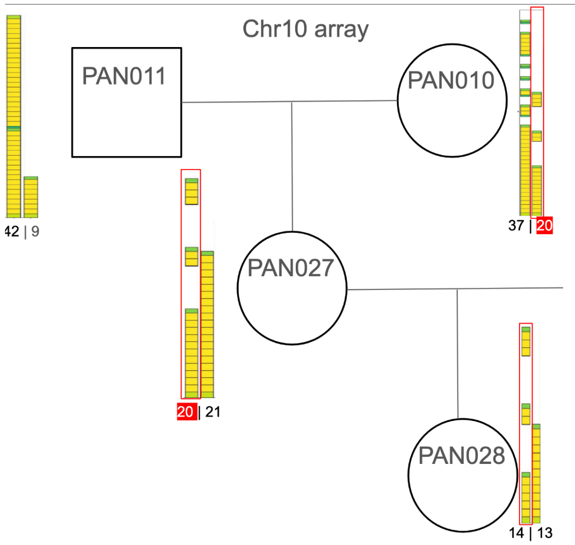


PAN010 - 20 repeats
PAN027 - 20 repeats

| Sample | Haplotype | chr10 subterminal repeats |
|---|---|---|
| chm13 | | 35 |
| PAN010 | hap1 | 37 |
| PAN010 | hap2 | 20 |
| PAN011 | hap1 | 42 |
| PAN011 | hap2 | 9 |
| PAN027 | mat | 20 |
| PAN027 | pat | 21 |
| PAN028 | hap1 | 14 |
| PAN028 | hap2 | 13 |

#### Supplementary Figure 7: Variant calling pipeline across the three generations

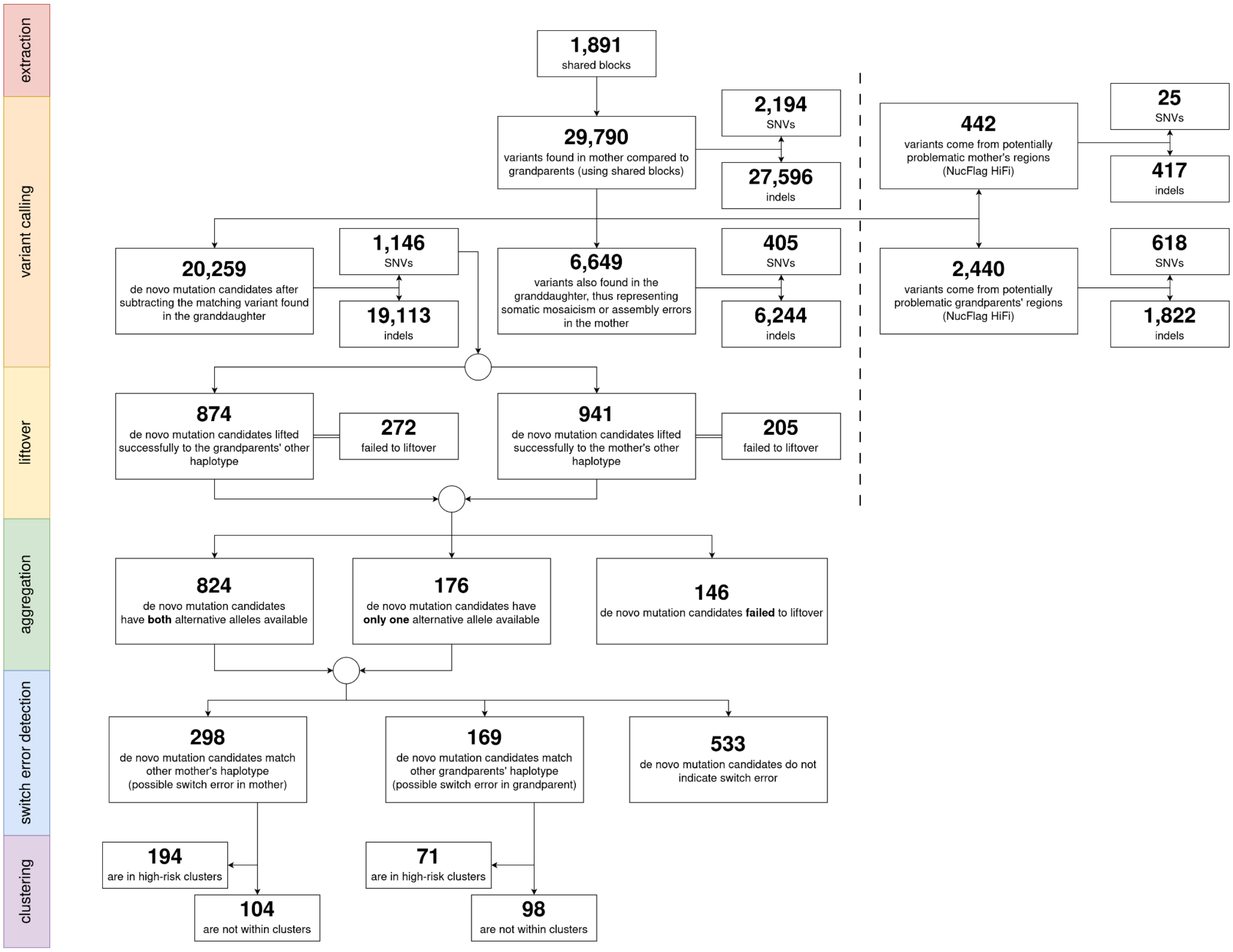

### Supplementary Figure 8: Genome-wide mutation profiles for SNVs

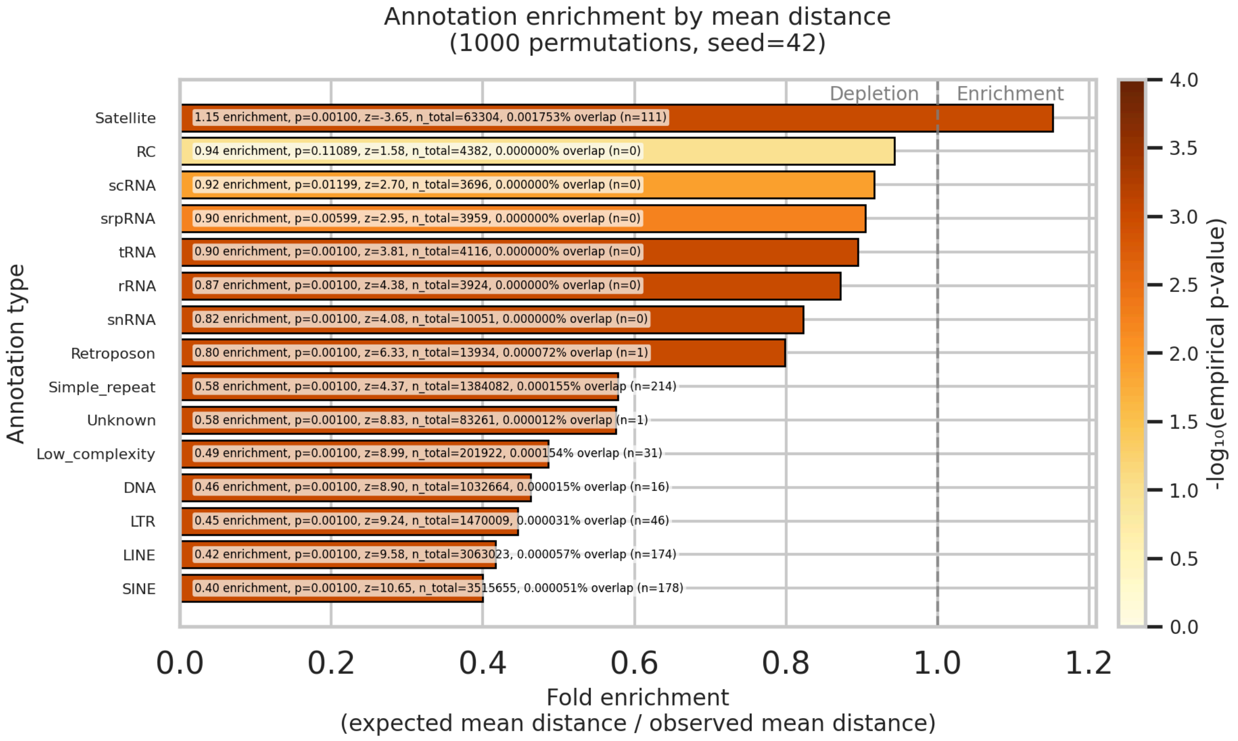


*RepeatMasker | SNVs*

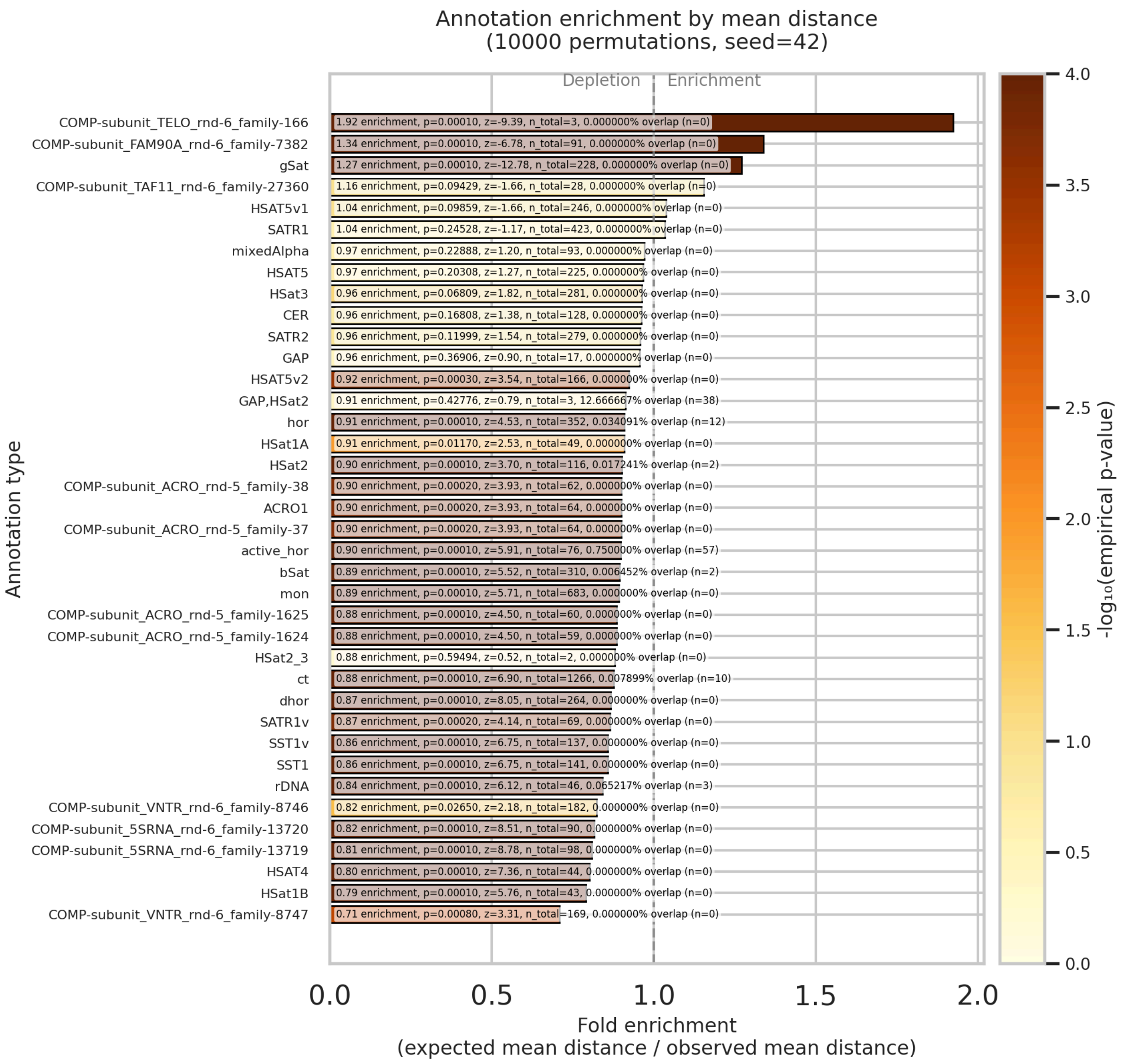
Annotation enrichment by mean distance
(10000 permutations, seed=42)
Depletion
Enrichment
COMP-subunit_TELO_rnd-6_family-166
1.92 enrichment, p=0.00010, z=-9.39, n_total=3, 0.000000% overlap (n=0)
COMP-subunit_FAM90A_rnd-6_family-7382
1.34 enrichment, p=0.00010, z=-6.78, n_total=91, 0.000000% overlap (n=0)
gSat
1.27 enrichment, p=0.00010, z=-12.78, n_total=228, 0.000000% overlap (n=0)
COMP-subunit_TAF11_rnd-6_family-27360
1.16 enrichment, p=0.09429, z=-1.66, n_total=28, 0.000000% overlap (n=0)
HSAT5v1
1.04 enrichment, p=0.09859, z=-1.66, n_total=246, 0.000000% overlap (n=0)
SATR1
1.04 enrichment, p=0.24528, z=-1.17, n_total=423, 0.000000% overlap (n=0)
mixedAlpha
0.97 enrichment, p=0.22888, z=1.20, n_total=93, 0.000000% overlap (n=0)
HSAT5
0.97 enrichment, p=0.20308, z=1.27, n_total=225, 0.000000% overlap (n=0)
HSat3
0.96 enrichment, p=0.06809, z=1.82, n_total=281, 0.000000% overlap (n=0)
CER
0.96 enrichment, p=0.16808, z=1.38, n_total=128, 0.000000% overlap (n=0)
SATR2
0.96 enrichment, p=0.11999, z=1.54, n_total=279, 0.000000% overlap (n=0)
GAP
0.96 enrichment, p=0.36906, z=0.90, n_total=17, 0.000000% overlap (n=0)
HSAT5v2
0.92 enrichment, p=0.00030, z=3.54, n_total=166, 0.000000% overlap (n=0)
GAP,HSat2
0.91 enrichment, p=0.42776, z=0.79, n_total=3, 12.666667% overlap (n=38)
hor
0.91 enrichment, p=0.00010, z=4.53, n_total=352, 0.034091% overlap (n=12)
HSat1A
0.91 enrichment, p=0.01170, z=2.53, n_total=49, 0.000000% overlap (n=0)
HSat2
0.90 enrichment, p=0.00010, z=3.70, n_total=116, 0.017241% overlap (n=2)
COMP-subunit_ACRO_rnd-5_family-38
0.90 enrichment, p=0.00020, z=3.93, n_total=62, 0.000000% overlap (n=0)
ACRO1
0.90 enrichment, p=0.00020, z=3.93, n_total=64, 0.000000% overlap (n=0)
COMP-subunit_ACRO_rnd-5_family-37
0.90 enrichment, p=0.00020, z=3.93, n_total=64, 0.000000% overlap (n=0)
active_hor
0.90 enrichment, p=0.00010, z=5.91, n_total=76, 0.750000% overlap (n=57)
bSat
0.89 enrichment, p=0.00010, z=5.52, n_total=310, 0.006452% overlap (n=2)
mon
0.89 enrichment, p=0.00010, z=5.71, n_total=683, 0.000000% overlap (n=0)
COMP-subunit_ACRO_rnd-5_family-1625
0.88 enrichment, p=0.00010, z=4.50, n_total=60, 0.000000% overlap (n=0)
COMP-subunit_ACRO_rnd-5_family-1624
0.88 enrichment, p=0.00010, z=4.50, n_total=59, 0.000000% overlap (n=0)
HSat2_3
0.88 enrichment, p=0.59494, z=0.52, n_total=2, 0.000000% overlap (n=0)
ct
0.88 enrichment, p=0.00010, z=6.90, n_total=1266, 0.007899% overlap (n=10)
dhor
0.87 enrichment, p=0.00010, z=8.05, n_total=264, 0.000000% overlap (n=0)
SATR1v
0.87 enrichment, p=0.00020, z=4.14, n_total=69, 0.000000% overlap (n=0)
SST1v
0.86 enrichment, p=0.00010, z=6.75, n_total=137, 0.000000% overlap (n=0)
SST1
0.86 enrichment, p=0.00010, z=6.75, n_total=141, 0.000000% overlap (n=0)
rDNA
0.84 enrichment, p=0.00010, z=6.12, n_total=46, 0.065217% overlap (n=3)
COMP-subunit_VNTR_rnd-6_family-8746
0.82 enrichment, p=0.02650, z=2.18, n_total=182, 0.000000% overlap (n=0)
COMP-subunit_5SRNA_rnd-6_family-13720
0.82 enrichment, p=0.00010, z=8.51, n_total=90, 0.000000% overlap (n=0)
COMP-subunit_5SRNA_rnd-6_family-13719
0.81 enrichment, p=0.00010, z=8.78, n_total=98, 0.000000% overlap (n=0)
HSAT4
0.80 enrichment, p=0.00010, z=7.36, n_total=44, 0.000000% overlap (n=0)
HSat1B
0.79 enrichment, p=0.00010, z=5.76, n_total=43, 0.000000% overlap (n=0)
COMP-subunit_VNTR_rnd-6_family-8747
0.71 enrichment, p=0.00080, z=3.31, n_total=169, 0.000000% overlap (n=0)
Annotation type
0.0
0.5
1.0
1.5
2.0
Fold enrichment
(expected mean distance / observed mean distance)
-log10(empirical p-value)
4.0
3.5
3.0
2.5
2.0
1.5
1.0
0.5
0.0

*CenSat | SNVs*

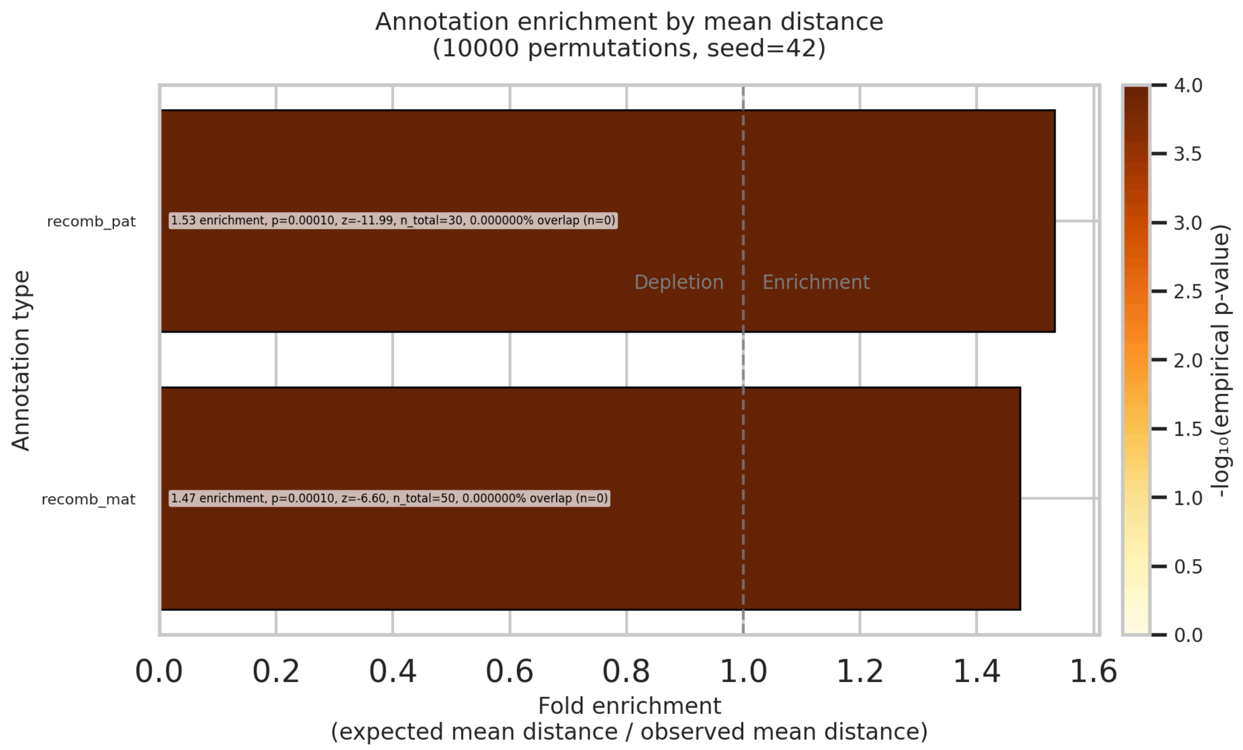


*Meiotic Recombination | SNVs*

### Supplementary Figure 9: Genome-wide mutation profiles for indels

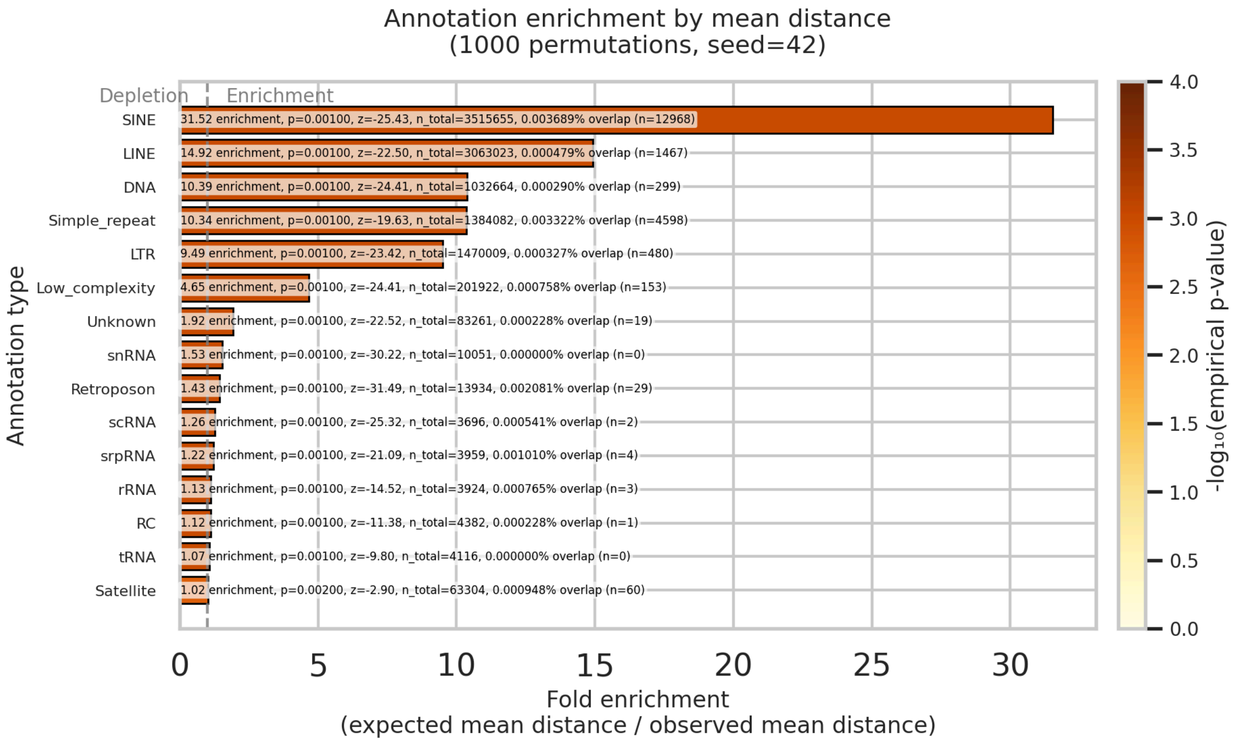


*RepeatMasker | indels*

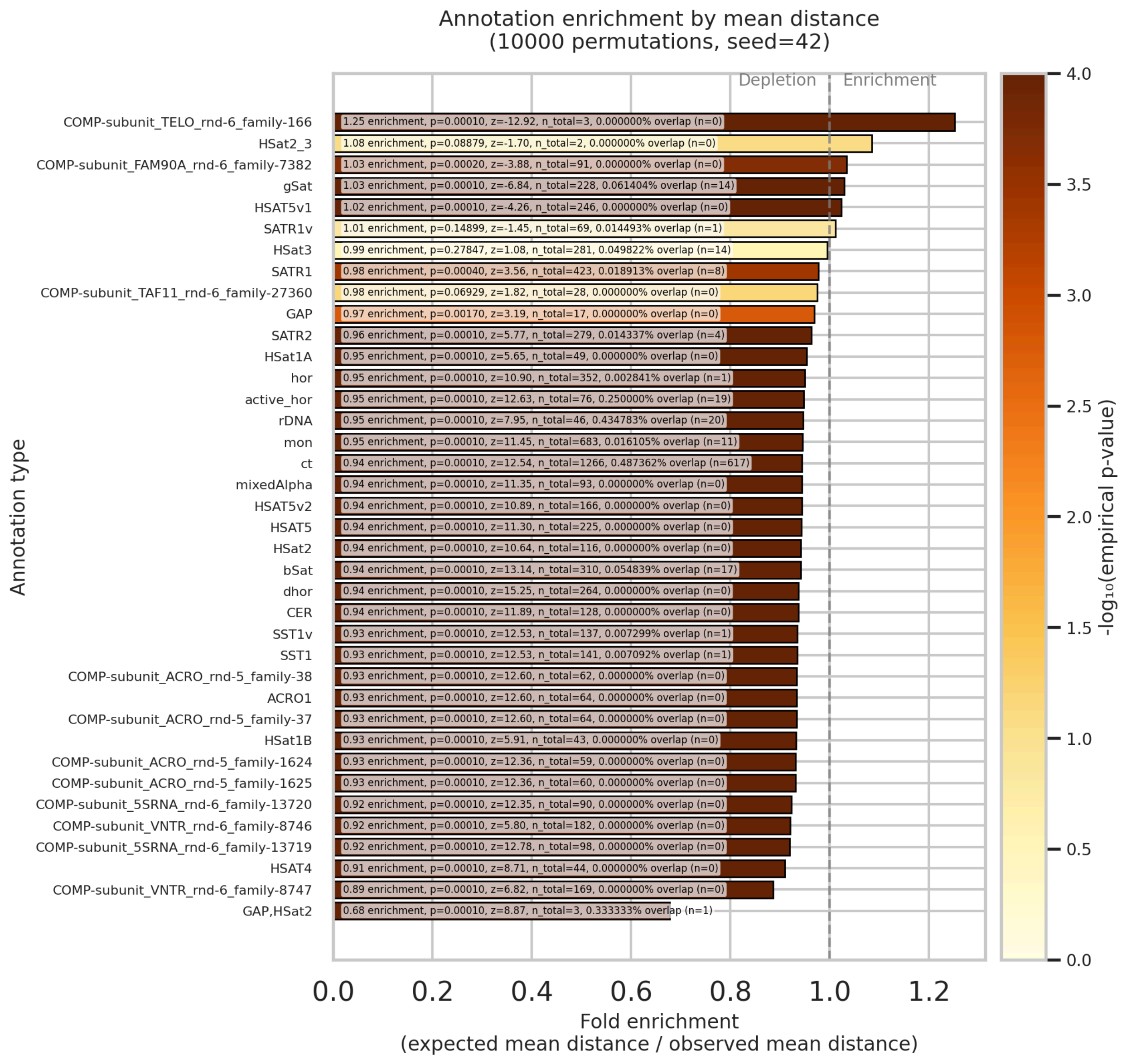


*CenSat | indels*

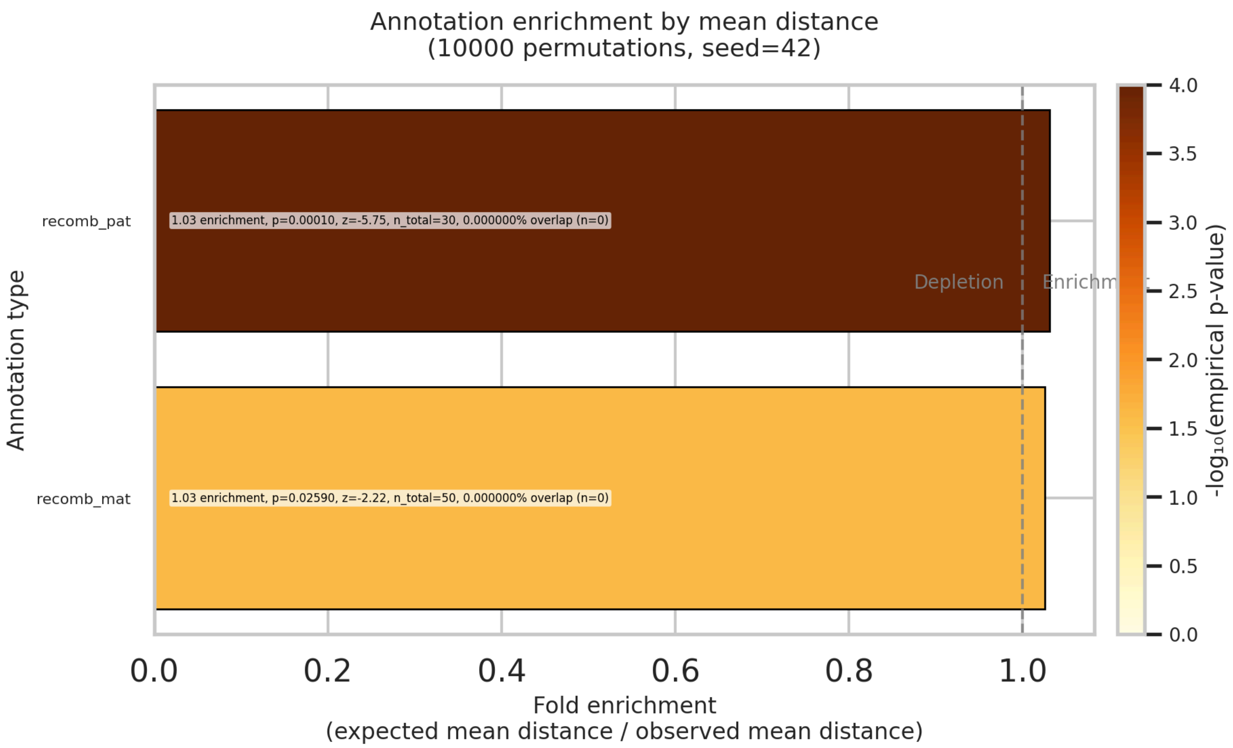


*Meiotic Recombination | indels*

### Supplementary Figure 10: Scaffolding of the rDNA distal and rDNA proximal sequences on acrocentric chromosomes

(A) A schematic representation of the scaffolding of rDNA-distal bits to the rest of the chromosome (rDNA-proximal bit, centromere, and q-arm of the chromosome). This way, despite a gap in the rDNA array, the whole chromosome can be represented by a single sequence. (B) In order to validate the general assignment of rDNA distal bits to the ten acrocentric chromosomes, we identified the presence and length of the WaluSat, satellite located on the human acrocentric chromosomes, and variable in length (Hoyt et al. 2022). The location in the assembled haplotypes matched the experimentally derived locations via FISH (see Supplementary Figure 11). Importantly, the length of the individual arrays matches perfectly in length across generations (each color represents one satellite being transmitted), with a single exception on chromosome 22, where the arrays differed only by 128 bps.

**A**

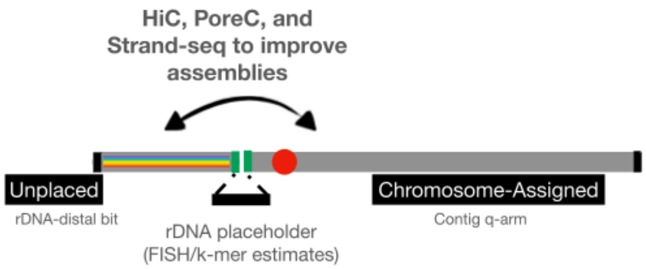


**B**

WaluSat: AGAAAGGGATAGGAGTGAAGAACACAGGTCGCTGCATTTAGAAAGGAGGCGGGGTCAGAGGAAT

| transmissions WALU_SAT | PAN010 | | PAN011 | | PAN027 | | PAN028 | |
|---|---|---|---|---|---|---|---|---|
| | hap1 | hap2 | hap1 | hap2 | maternal | paternal | hap1 | hap2 |
| chr13 | 8,940 | 1,006 | 7,982 | 9,838 | 8,940 | 9,838 | 8,940 | 10,700 |
| chr14 | 9,708 | 1,583 | 46,255 | 1,292 | 1,583 | 46,255 | 1,356 | 46,255 |
| chr15 | 12,208 | 44,549 | 6,895 | 19,824 | 44,549 | 19,824 | 12,333 | 19,824 |
| chr21 | 9,198 | 4,655 | 69,656 | 37,604 | 9,198 | 69,656 | 98,475 | 69,656 |
| chr22 | 4,073 | 17,258 | 17,263 | 2,031 | 17,130 | 2,031 | 5,038 | 2,031 |

#### SCAFFOLDING PROCEDURE

To perform the scaffolding, we first map poreC reads using minimap2 and filter to only retain alignments with MAPQ 10 or higher, representing a compromise between the quality of mapping and the number of mapped reads. In poreC, a single read will carry sequences from potentially distant sequences in the linear space that are in close proximity in 3D chromatin structure. Unlike Hi-C reads -- that always have a single pair of sequences -- poreC will contain multiple concatemers. In the current implementation, we are converting these multi-contacts into pairwise contacts, although incorporating multi-contact information could improve the performance. For example, a single poreC read carrying sequences A, B, C will be treated as three interactions: A-B, B-C, and C-A. In terms of normalization (since various contigs will have various lengths and thus the number of poreC reads/contacts), we look up all contacts for a specific p-arm containing contig, and use mean of this numerical vector. For each candidate q-arm containing contig, we calculate qcontacts/mean_across_all. We visualize this in the heatmap to select the most likely p-arm and q-arm assignments for each haplotype of acrocentric chromosomes, considering all the combinations and guided by the highest numbers.

#### Supplementary Figure 11: The most variable genes in the transmitted regions

| | ACRO1 COPIES | | | | | USP17L COPIES |
|---|---|---|---|---|---|---|
| | **chr13** | **chr14** | **chr15** | **chr21** | **chr22** | **chr4** |
| **PAN010_hap1** | 21 | 2 | 2 | 6 | 1 | 66 |
| **PAN010_hap2** | 9 | 2 | 4 | 0 | 8 | 63 |
| **PAN011_hap1** | 9 | 2 | 0 | 5 | 4 | 41 |
| **PAN011_hap2** | 18 | 5 | 11 | 1 | 4 | 15 |
| **PAN027_mat** | 13 | 2 | 4 | 4 | 5 | 62 |
| **PAN027_pat** | 16 | 2 | 11 | 5 | 4 | 40 |
| **PAN028_hap1** | 14 | 1 | 5 | 2 | 1 | 61 |
| **PAN028_hap2** | 3 | 2 | 7 | 4 | 4 | 64 |

The first part of the Table shows the copy number variability of of ACRO1 gene across the 8 haplotypes and the five acrocentric chromosomes. Highlighted are the p-arms that were transmitted across the three generations. The second part of the Table shows the copy number for USP17L. Together, ACRO1 and USP17L are the most variable genes in the transmitted regions of all four family members. In contrast, other genes distributed genome-wide but transmitted across the three generations were stable.

We further restricted our analysis to the p-arms of acrocentric chromosomes, as we hypothesized that they would be the regions with the highest variation. We studied pseudogenes restricted to acrocentric short p-arms and the transmitted regions across three generations, and found them to be stable.

USP17L gene is located in clusters on both chromosomes four and eight in the human genome. The chromosome eight cluster shows stable transmission across the three generations, whereas the cluster on chromosome four is variable across generations. Below, we are zooming in on the cluster on chromosome four. In pink, we are showing the genes; in blue, we are showing cenSat(SATR1,SATR2) , cenSat(HSAT5v1), cenSat(SATR1) & similar.

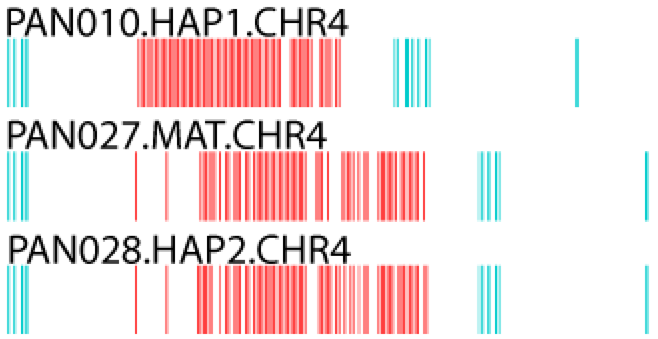


**USP17L gene on chromosome 4**

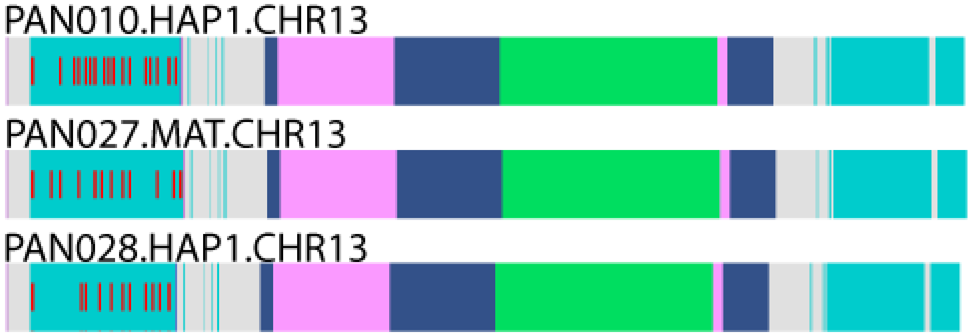


**The ACRO1 transmission changes across chromosome 13.** The short red bars represent the putative ACRO1 genes, the colored rectangles represent the cenSat annotations, as described previously.

### Supplementary Figure 12: rDNA FISH

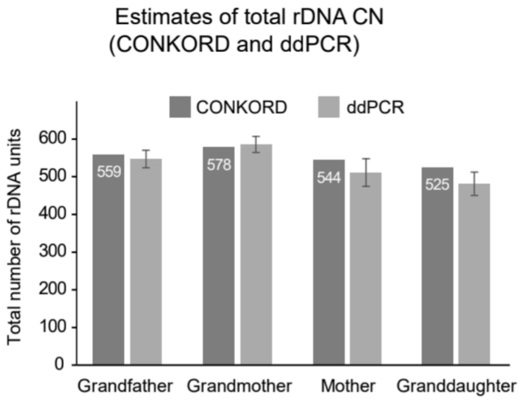


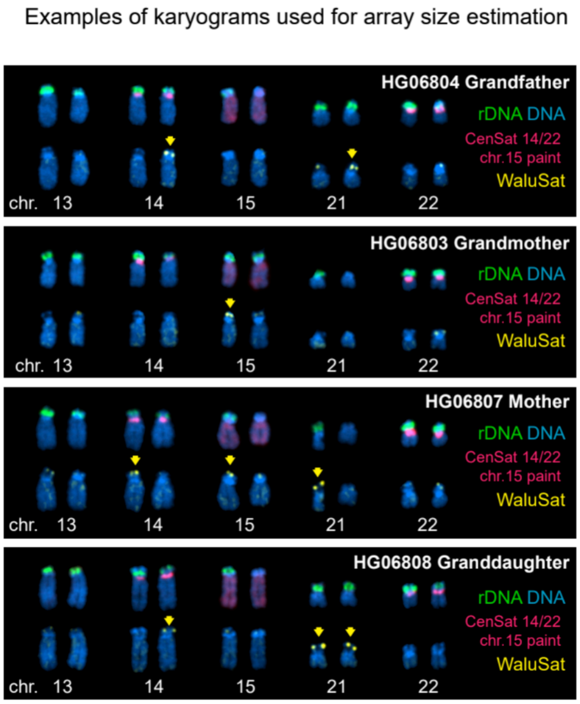


**ddPCR panel (left)**

Total rDNA copy number estimates obtained by DNAseq using the CONKORD pipeline (dark gray bars with numbers) and ddPCR (light gray bars). ddPCR estimates represent an average copy number calculated from four rDNA/single copy gene ratios:18S/TBP, 28S/TBP, 18S/RPP30, and 28S/RPP30. Bars denote the standard deviation of copy numbers obtained from these ratios.

**FISH image (right)**

Representative acrocentric karyograms from cell lines used in this study. The top rows show chromosomes labeled by FISH with rDNA probe (green) and chromosome identification markers - CenSat 14/22 and whole chromosome 15 paint (red) serving as identification markers for chromosomes 14, 22 (distinguished by size), and chromosome 15. Chromosome 13 was identified as a large unlabeled acrocentric, and chromosome 21 as a small unlabeled acrocentric. Bottom rows show the corresponding chromosomes labeled with WaluSat probe (yellow signal present on some acrocentric p-arms), with large signals highlighted by arrows. DNA was counter-stained with DAPI.

#### Supplementary Figure 13: rDNA sequence quality improvements with hyperbasecalling

We compared the same subset of the rDNA reads basecalled twice, once with the super-accuracy model, and once with the hyperbasecalling model, combining the whole-genome sequencing reads with adaptive sampling reads.

The quality and the length distribution of super-accuracy reads:

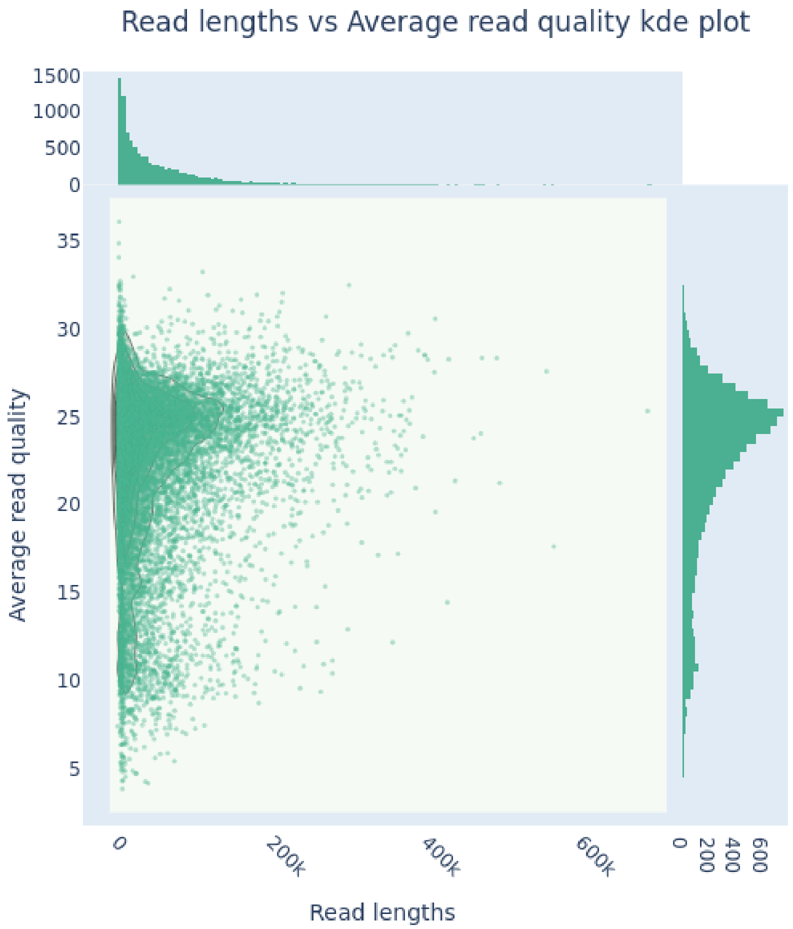

The quality and the length distribution of hyperbasecalled reads:

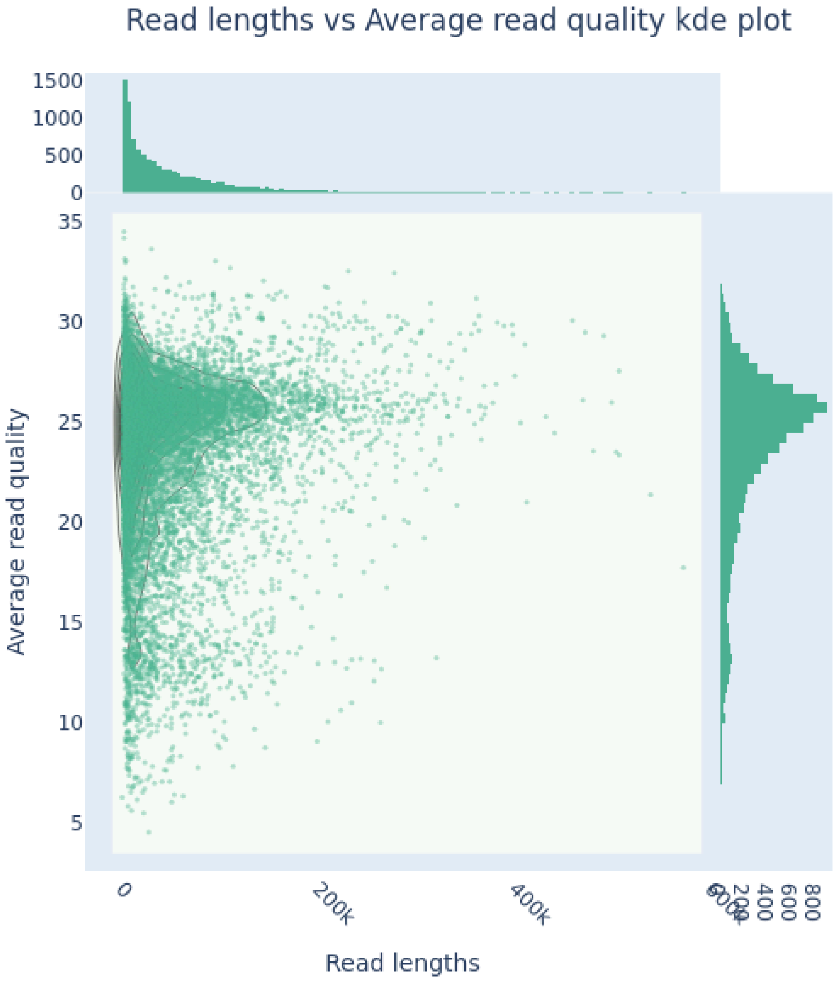

Adaptive sampling reads that the median of 99.16% identity between the sup and hyper reads. This was calculated using the formula of 1 - (mismatches + gaps) / length when aligning the identical reads basecalled with the two basecalling algorithms using stretcher.

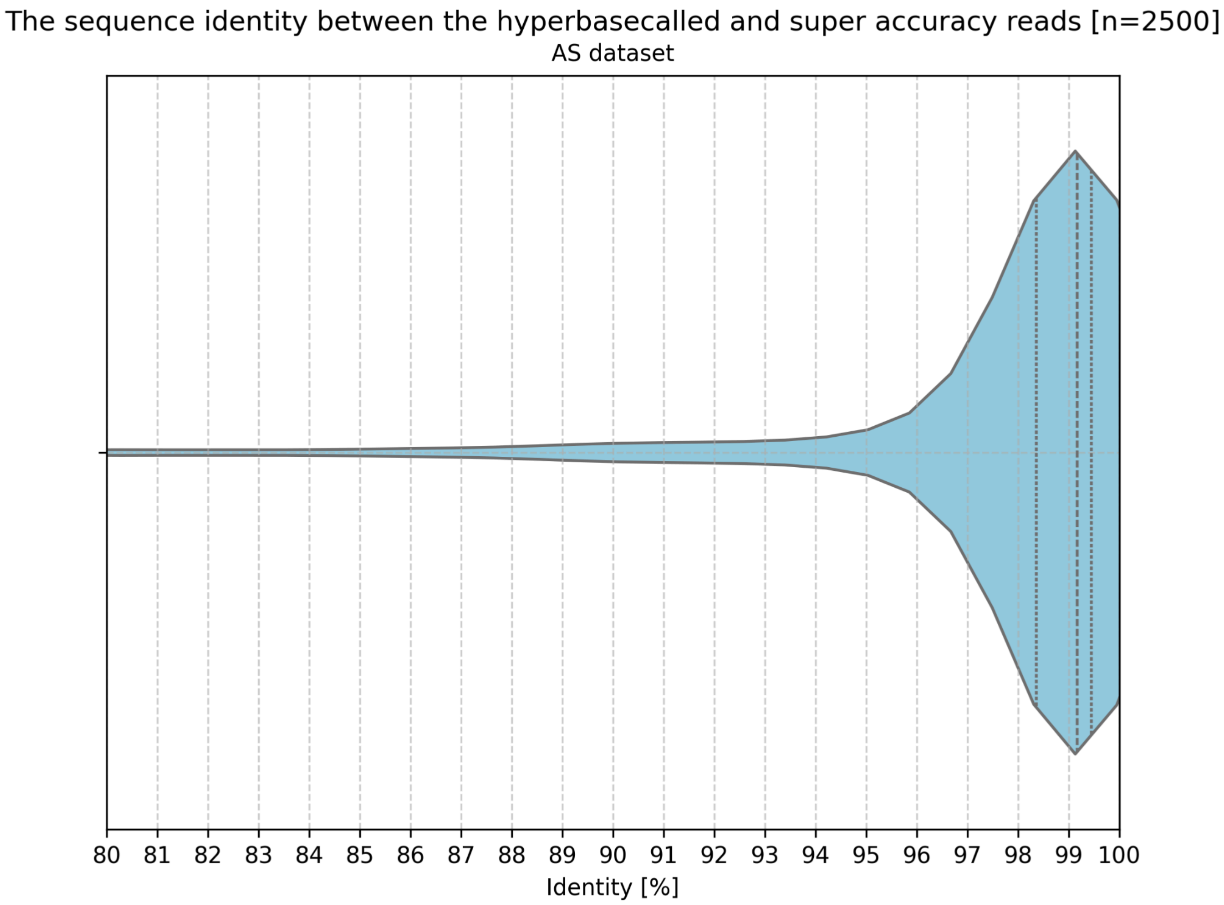

Whole-genome sequencing reads have a median of 99.12% identity between the sup and hyper reads. This was calculated using the formula of 1 - (mismatches + gaps) / length when aligning the identical reads basecalled with the two basecalling algorithms using stretcher.

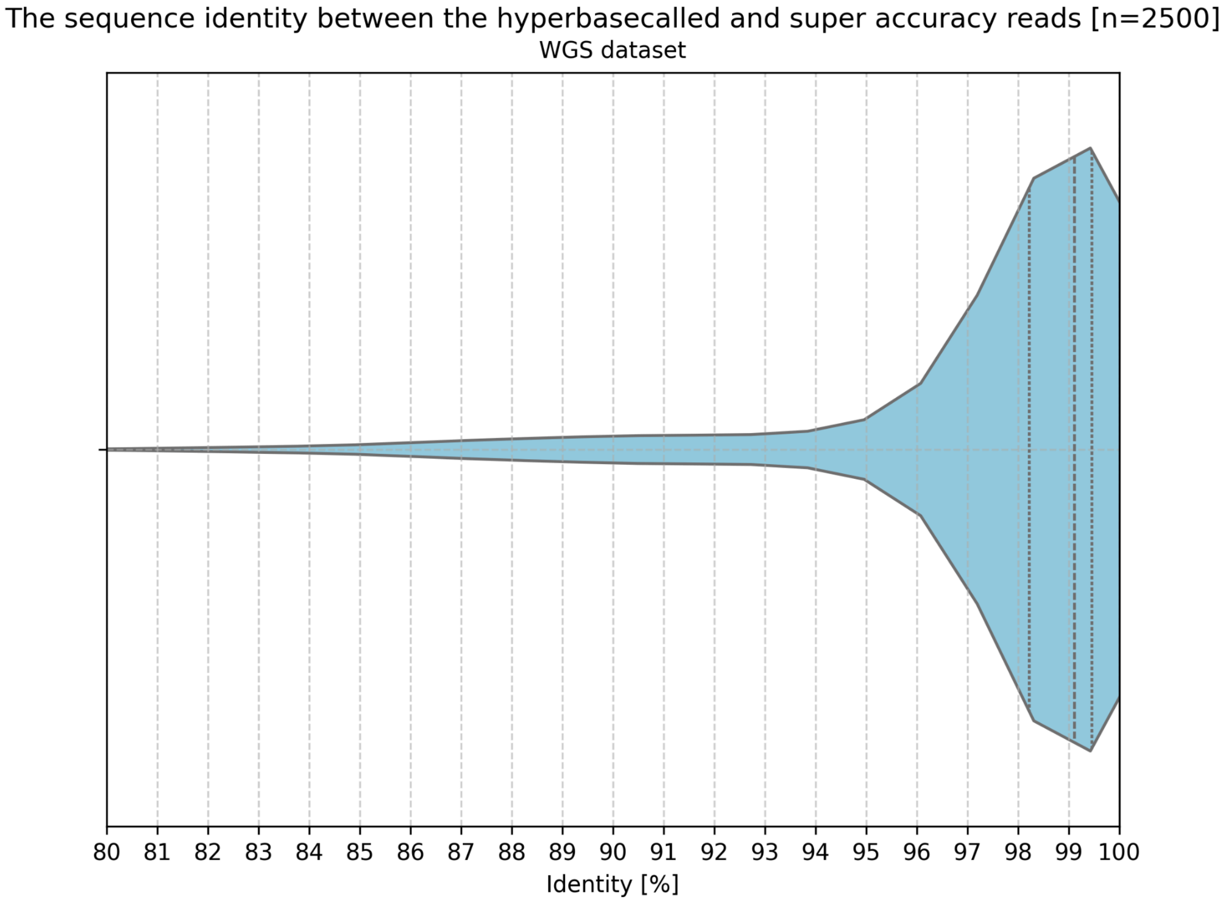

The read length and the % identity between the sup- and hyper-basecalled reads didn’t have a specific relationship, with the exception of longer reads all being high identity.

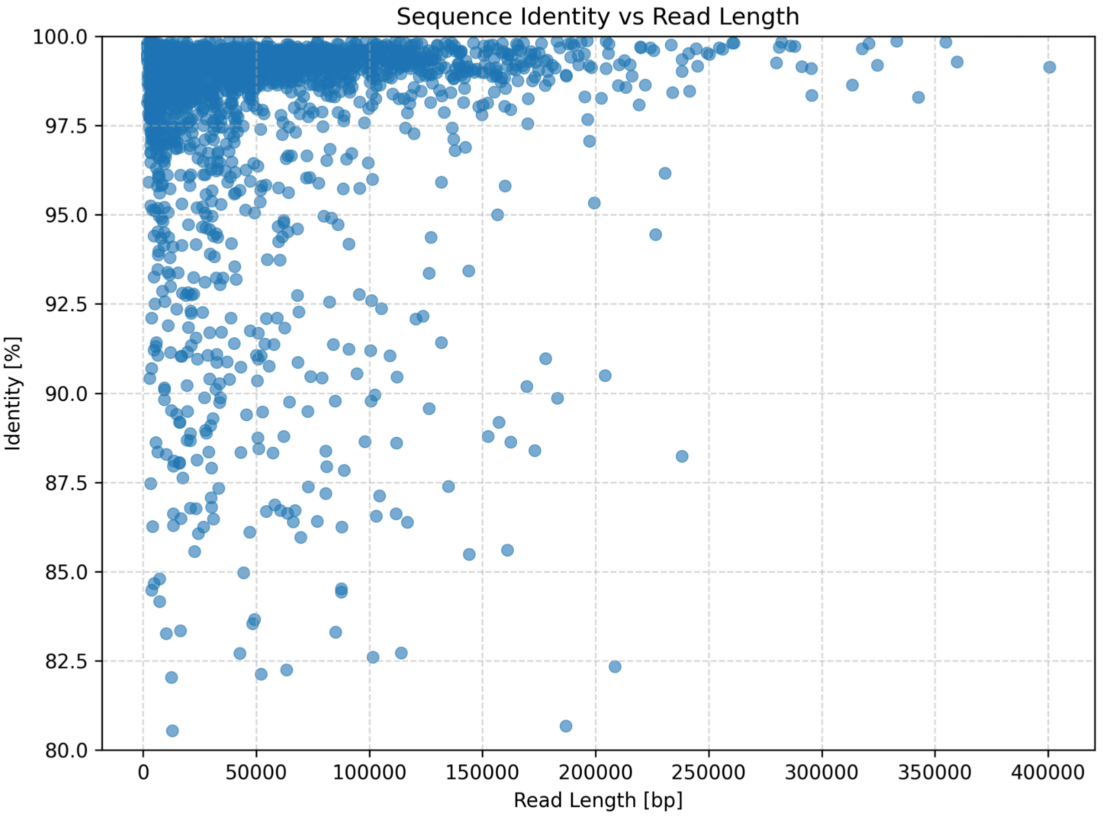

The individual introduced changes between AS and WGS were dominated by single-basepair indels, especially those involving A and G nucleotides.

The Table below calculates the changes using 1,000 random reads (the first 1,000 read names in alphabetical order).

Adaptive sampling

| Count | Hyper | Sup |
|---|---|---|
| 90403 | G | - |
| 81956 | A | - |
| 74504 | - | A |
| 73847 | - | G |
| 68964 | C | - |
| 67655 | G | A |
| 61565 | A | G |
| 59822 | T | - |
| 58952 | - | C |
| 54834 | - | T |
| 37398 | G | C |
| 34755 | C | G |
| 30481 | A | C |
| 27658 | T | C |
| 26851 | C | A |
| 26522 | C | T |
| 20484 | A | T |
| 19176 | G | T |
| 16158 | T | G |
| 15130 | T | A |

WGS

| Count | Hyper | Sup |
|---|---|---|
| 124638 | - | G |
| 113277 | - | A |
| 73907 | A | G |
| 68082 | G | A |
| 62242 | - | T |
| 57296 | - | C |
| 55736 | G | - |
| 50287 | A | - |
| 47792 | C | - |
| 35315 | C | G |
| 32148 | T | - |
| 27767 | G | C |
| 25023 | C | A |
| 20069 | C | T |
| 19862 | A | C |
| 19323 | T | C |
| 15524 | T | A |
| 12491 | A | T |
| 12482 | G | T |
| 10879 | T | G |

#### Supplementary Figure 14: rDNA methylation analysis comparing the 45S gene and the promoter

Based on our analysis, the 45S gene and the promoter provided the same information about the activity of rDNA units, as their values were highly correlated with each other.

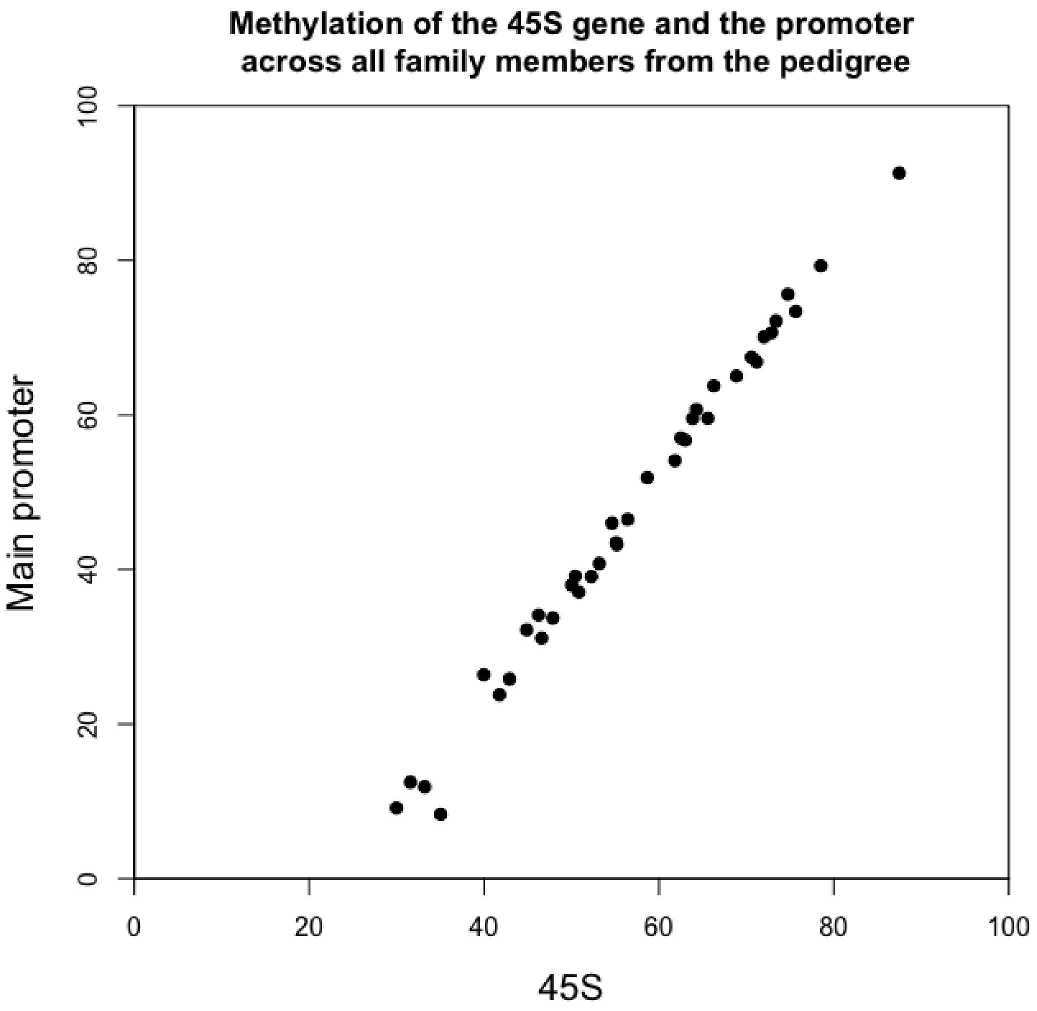

#### Supplementary Figure 15: G4s in rDNA

G4s on the positive strand, as detected by G4Hunter (score>1.5), were split into quartiles from Q1 (lowest score) to Q4 (highest score). G4s with the highest score were among the least conserved across rDNA arrays, suggesting their potential functional significance facilitated by variable G4 formation or disassembly.

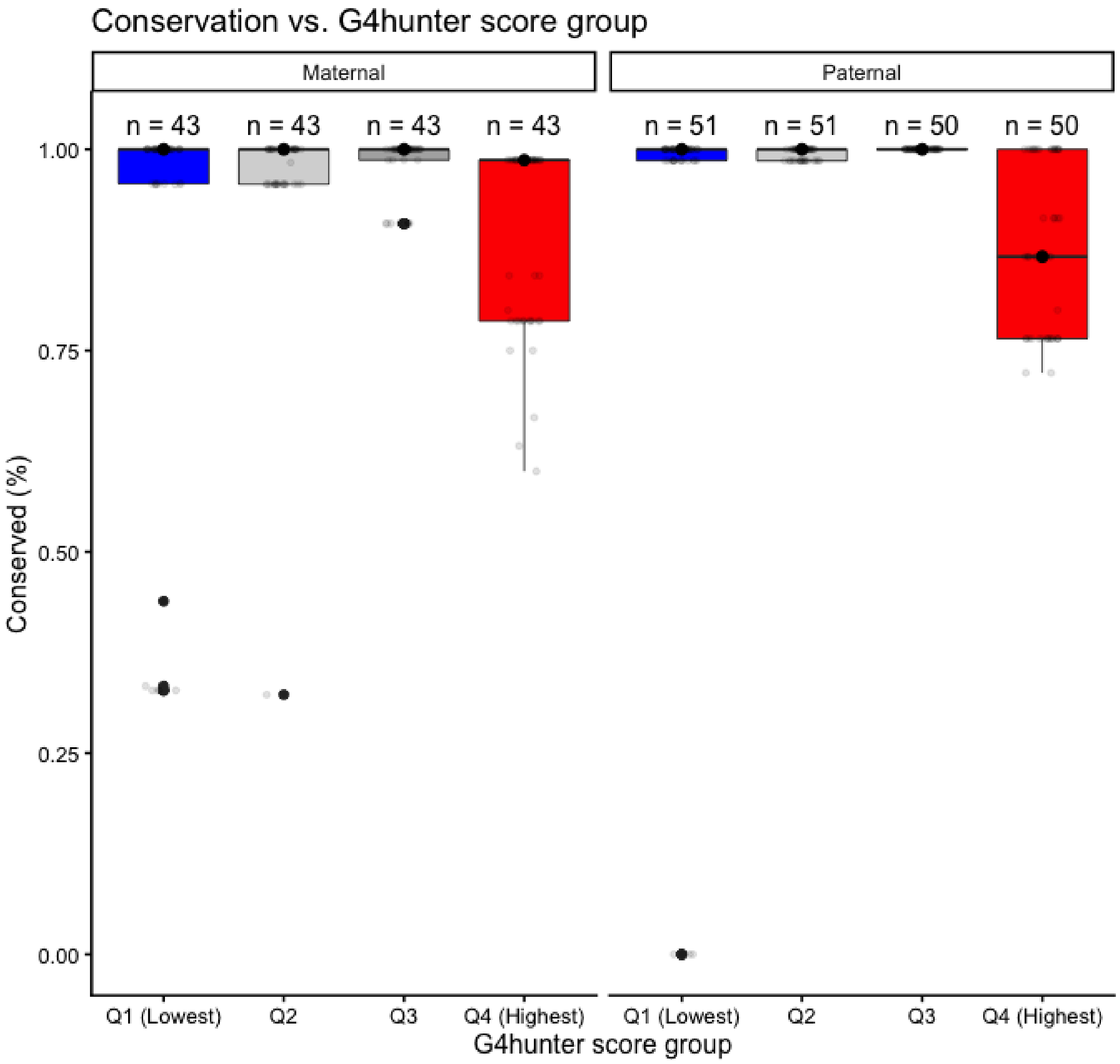

#### Supplementary Figure 16: Centromere characterization

Dinucleotide repeat content shows that all centromeres are rich for AA and TT dinucleotides, while underrepresented in GC nucleotides, despite exhibiting high de novo mutation rates in this context.

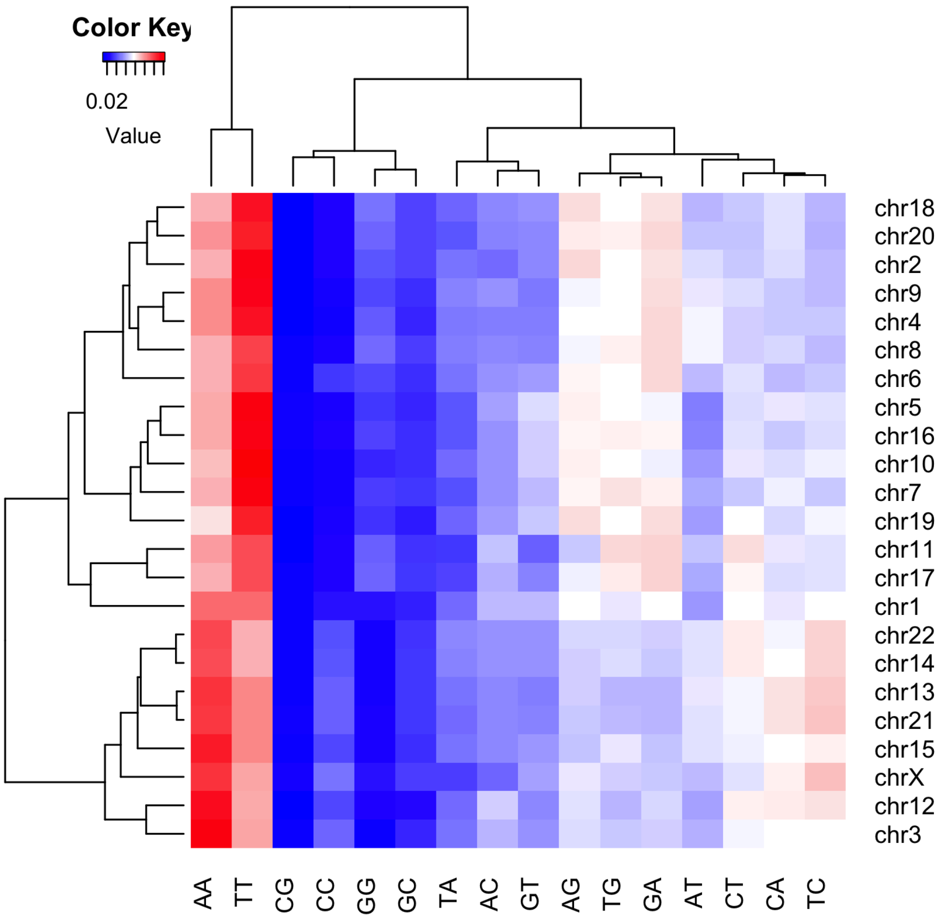

Non-B DNA content of the same centromeres.

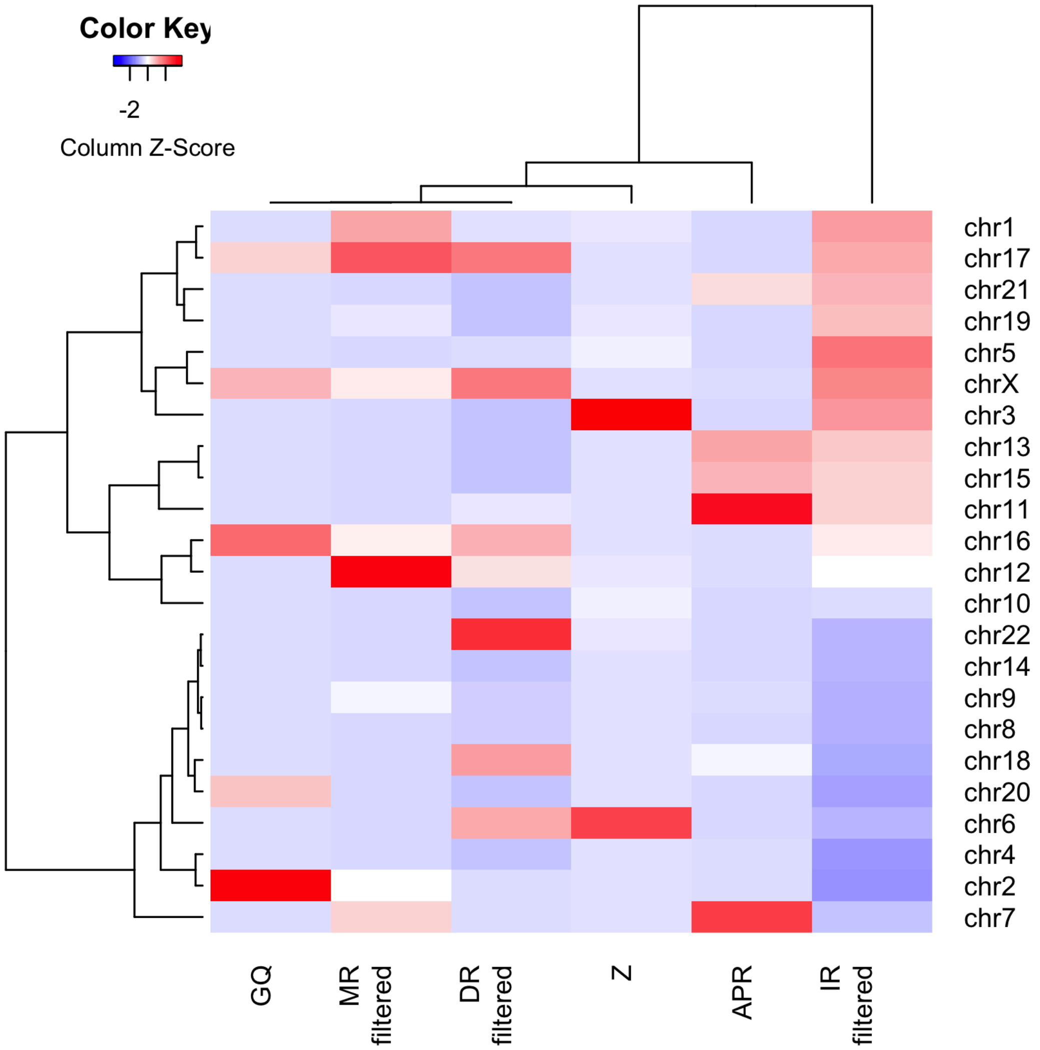

#### Supplementary Figure 17: Validations of de novo mutation candidates in centromeres

While the profile of the background mutation spectra matched our expectations for de novo germline variants, candidate centromeric variants exhibited unique mutation spectra dominated by G->C transversions (listed direction in dark color, the light color represents the reverse complement). However, only four SNVs were validated using blood-derived HiFi reads, specifically G->C, T->C, A->G, T->A (Supplementary Table “SNV_validation”).

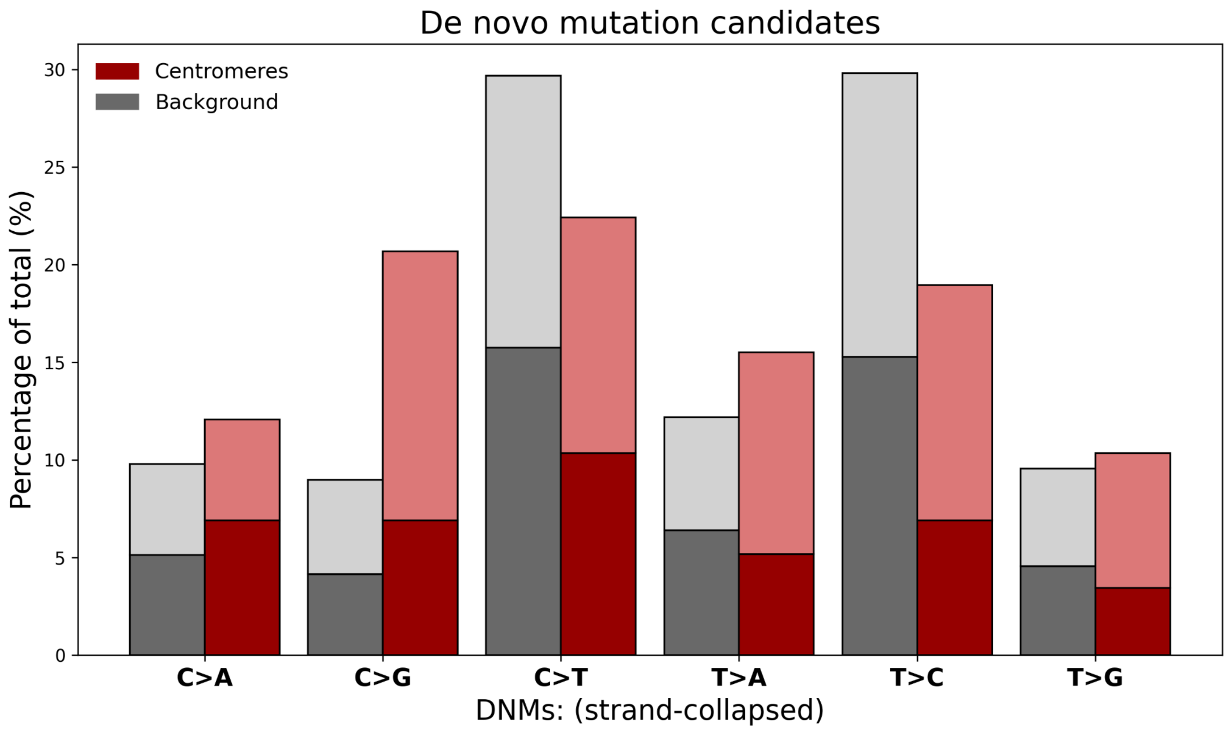

#### Supplementary Figure 18: Structural changes in chromosome 9 centromere

**Dotplot**

A dotplot of active ASat array sequences from *PAN011.chr9_haplotype1* and *PAN027.chr9_paternal* was generated using the Gepard program [https://doi.org/10.1093/bioinformatics/btm039]. Array coordinates were defined as the region spanning from the first to the last HOR based on horhap annotations (*PAN011.chr9_haplotype1*: 45,210,616–47,901,728; *PAN027.chr9_paternal*: 45,213,245–47,952,837). The word length parameter was set to 2000.

The diagonal, representing perfect sequence alignment, is interrupted at two positions. A red dotted line marks an insertion, and a red arrow indicates a deletion that occurred during the transmission from *PAN011* to *PAN027*. The tracks represent horhap annotations and CDR positions.

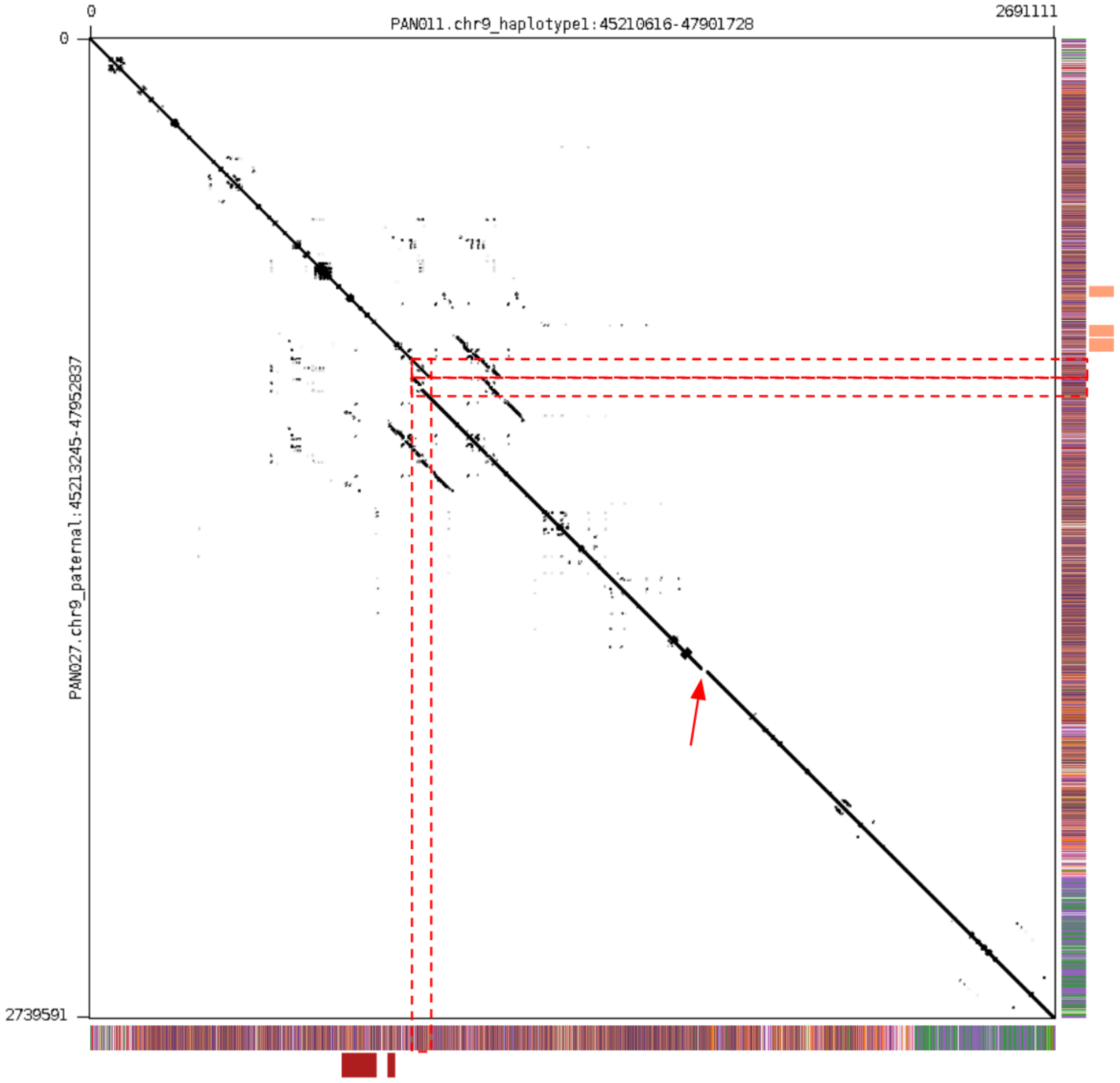

##### Horhap annotation alignment

K6 horhap annotations for *PAN011.chr9_haplotype1*, *PAN027.chr9_paternal, PAN028.chr9_haplotype2* were aligned using custom implementation of the Needleman-Wunsch algorithm (https://github.com/fedorrik/annotaligner). The alignment was completely identical except for two indel events, which were consistent with the dotplot results.

The first event is a presumably germline inherited insertion of a 56.1 kbp segment containing 69 HORs (HORs #1047–#1115; *PAN027.chr9_paternal*:46,160,135–46,216,271).

The second is a 7.7 kbp presumably somatic deletion encompassing 9 HORs (HORs #1972–#1980 alignment coords or #1903–#1911 PAN011 coords; *PAN011.chr9_haplotype1*:46,906,049–46,913,705).

##### Duplication nature of insertion event

Prior to the Inserted 69 HORs (#1047–#1115) there is a sequence of 57 HORs (#989–#1046; PAN027.chr9_paternal:46112503-46160135) which is the same as in inserted peace, except 11 extra HORs in the inserted peace (#1076–#1086) PAN027.chr9_paternal:46183951-46192455.

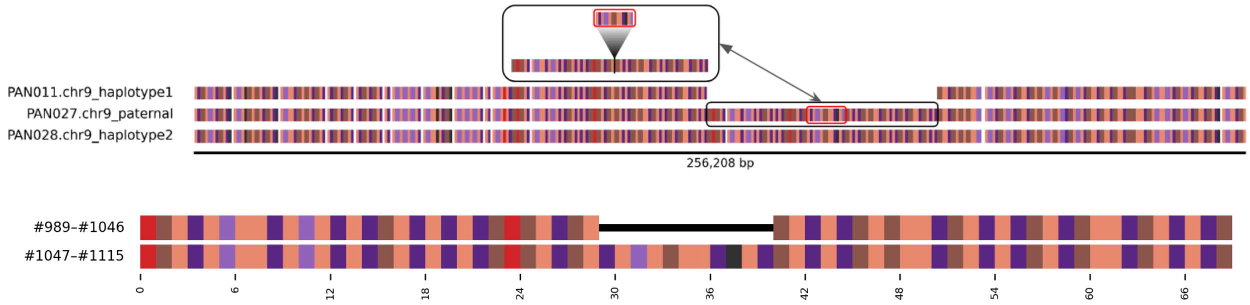


So, presumably #989–#1046 were duplicated and 11 extra HORs were inserted, likely after the duplication. The same 11 HOR sequence is in #1307–#1317 (PAN027.chr9_paternal:46382318-46390822), which is ~150kb away from the end of the duplication.

##### Mapping the secondary insertion (11-HOR piece)

The nucleotide sequence comparison of the 11-HOR pieces #1076–#1086 and #1307–#1317 showed the presence of 18 SNPs per 8.5 kbp. Most of these mutations are localized in the first 1 kbp, suggesting that the first 1 kbp has a different origin, whereas the remaining 7.5 kbp were transferred to #1076–#1086 from #1307–#1317.

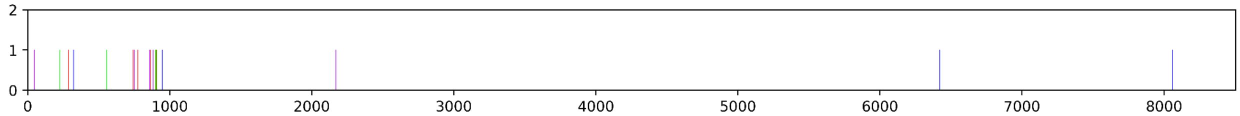


Minimap2 mapping of the 11 extra HORs from the duplication (#989–#1046) to the complete chr9 of PAN027 resulted in two hits:

1. With itself (#989–#1046), but with 34–315 bp trimmed at the edges

2. With the #1307–#1317 piece, but with 983 bp trimmed from the left (removing most of these 18 SNPs) and with 315 bp trimmed at the right (same as in 1. Technical issue of minimap2 (?))

Minimap2 of the first 950 bp of the 11 extra HORs gave 6 hits, including one hit within 10 kbp upstream of the #1307–#1318 minimap hit.

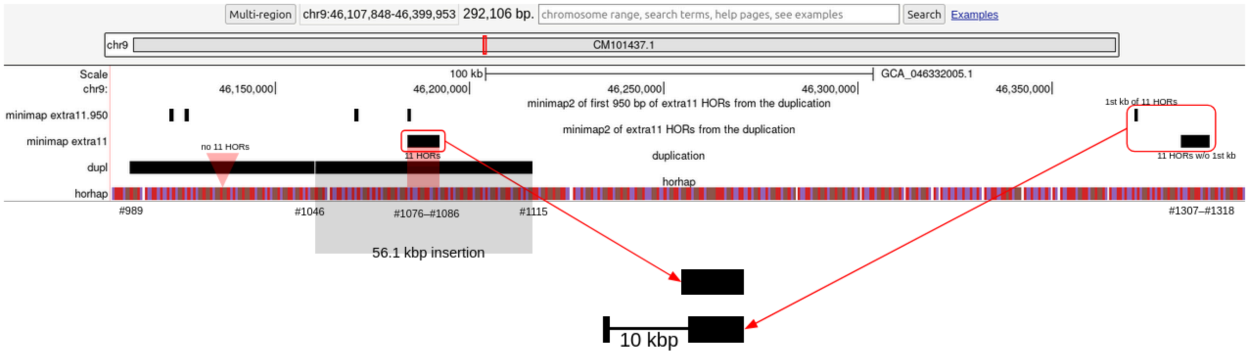


The upper track indicates Minimap2 hits of the first 950 bp of the 11 extra HORs, the track below that indicates Minimap2 hits of the 11 extra HORs, the next track shows the entire duplication, and the track on the bottom is a horhap annotation (with slightly different coloring compared to all other horhap tracks in this note).

**Alternative locations of indels**

The annotation aligner selects one of a few potential positions for a gap. As is obvious from the figure below (somatic deletion of PAN011 HORs #1903–#1911), the position of the gap can be shifted to various positions while the alignment score remains the same. So, in fact, we only show one of the possible positions.

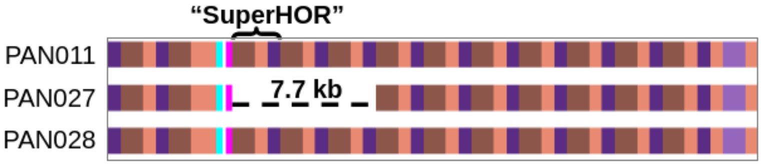


With the same idea, the germline duplication described above can occur not only by scenario A in the figure below (as described above) but also by scenario B. Colored panels between A and B indicate CDR positions in PAN011, PAN027, and PAN028, respectively.

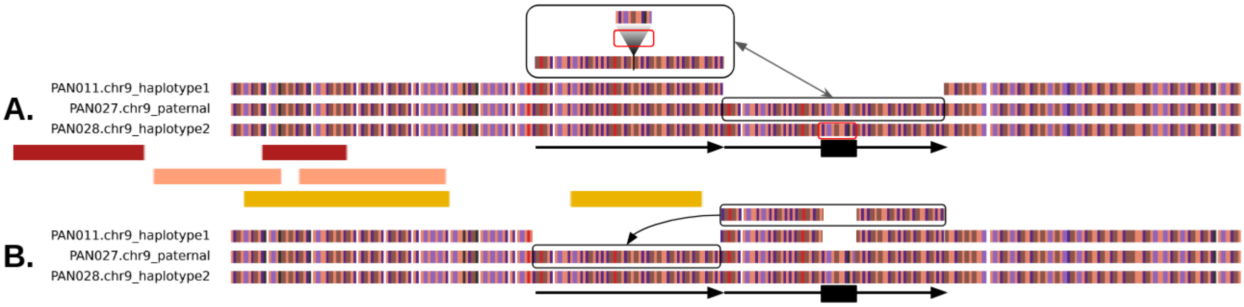

### Supplementary Figure 19: Segmental duplications

Two examples of the transmission of subtelomeric regions, visualized with the segmental duplications as detected by biser. Note that segmental duplications were reported independently for each haplotype, and thus their presence/absence might be influenced by the assortment of individual chromosomes into haplotypes, and not necessarily the presence/absence of specific sequences. The stars mark the transmitted haplotypes, as assigned for the analysis of telomeres.

*p*-arm of chromosome 4:

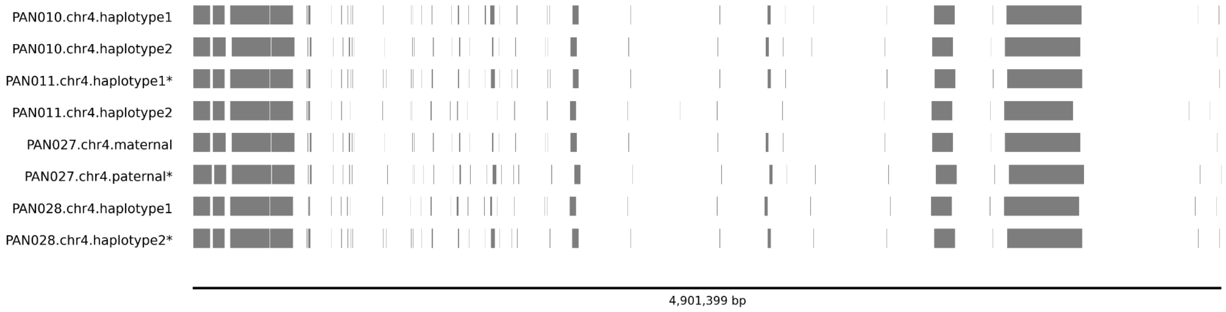


*q*-arm of chromosome 4:

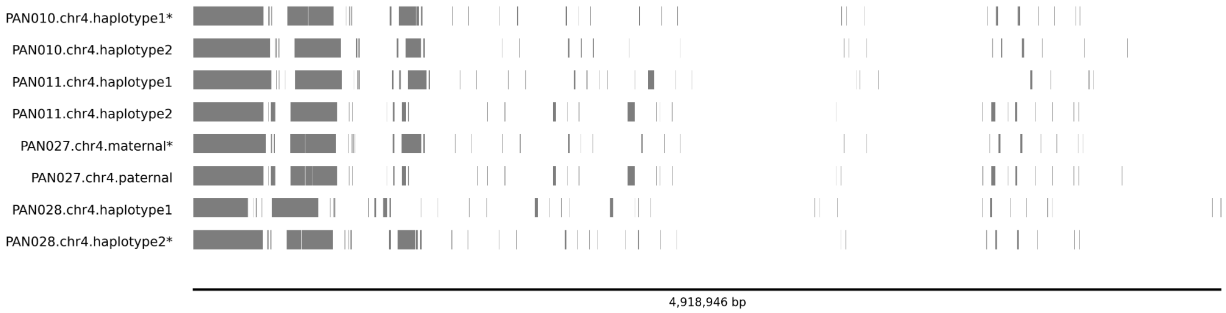


*p*-arm of chromosome 16:

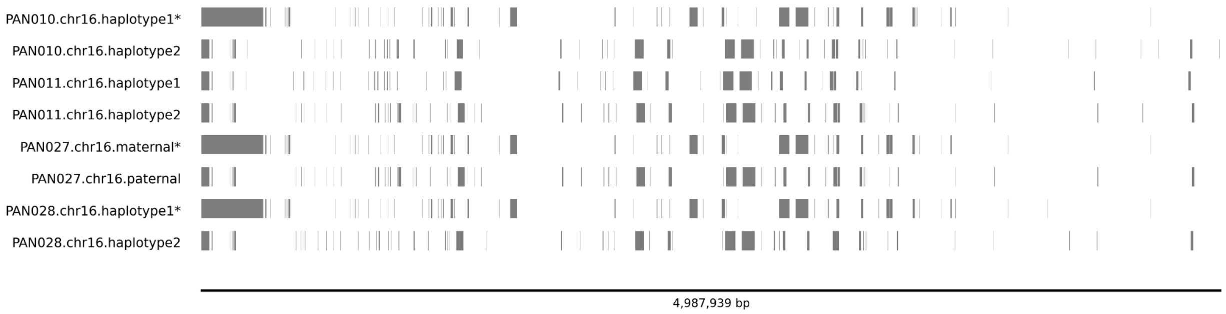

*q*-arm of chromosome 16:

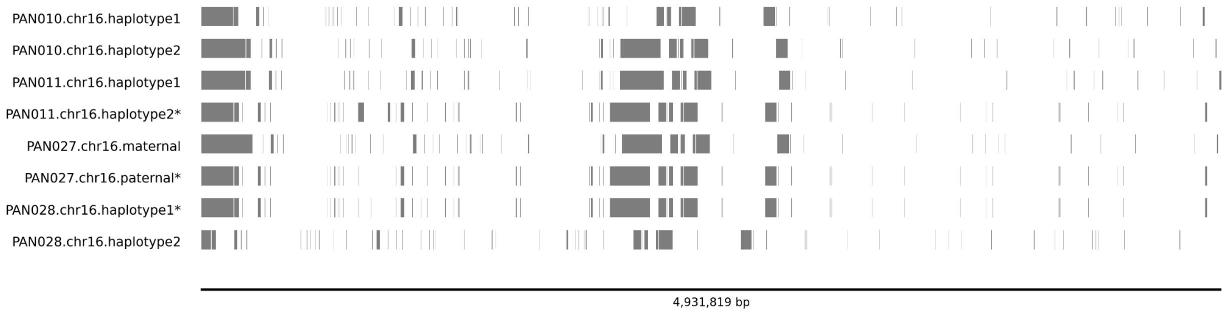

#### Supplementary Figure 20: Pedigree telomere lengths

The telomere length and read length distributions of the datasets used for establishing telomere length. The similarity of the read distributions in all four individuals suggests that the derived telomere lengths are not driven by read lengths. This is especially relevant for the grandfather (PAN011) that had significantly shorter telomeres than the rest of the family. Note that mother (PAN027) was sequenced deeper than the rest of the individuals.

**TELOMERE LENGTHS:**

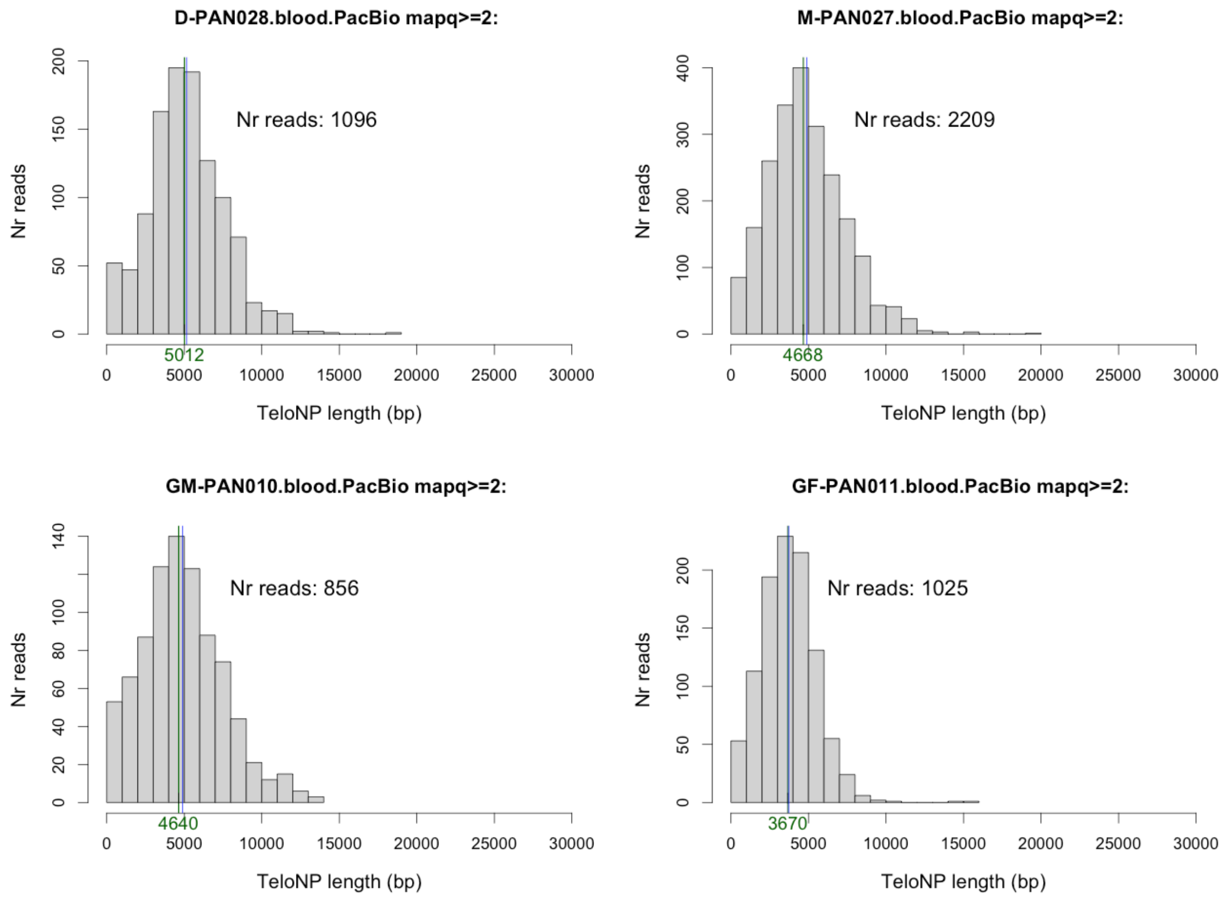

### READ LENGTHS:

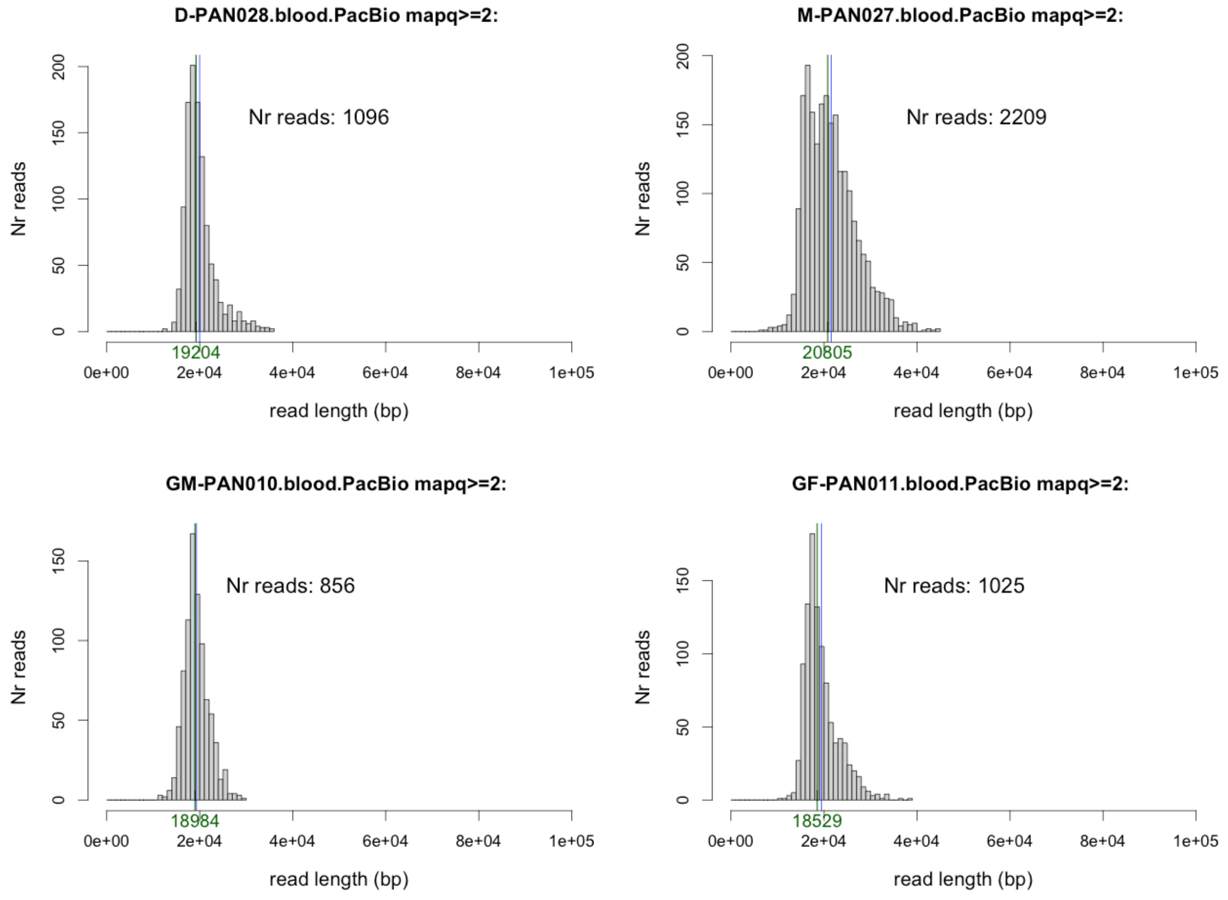

### Supplementary Figure 21: Telomere length chromosome ordering

The telomere lengths of the pooled blood samples (4 individuals) and pooled ONT cell lines (3 individuals, excluding the grandmother since her cell-line data more closely resembled blood) were correlated with the previously published population-based estimates (Karimian et al. 2024).

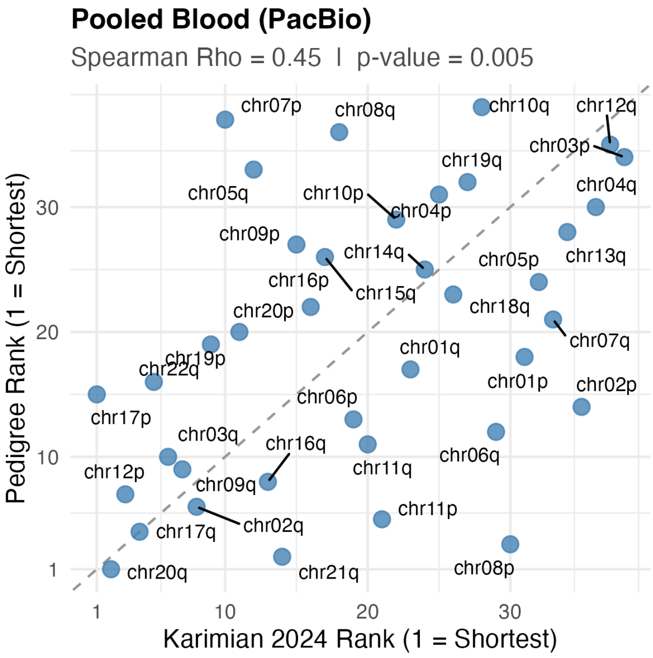


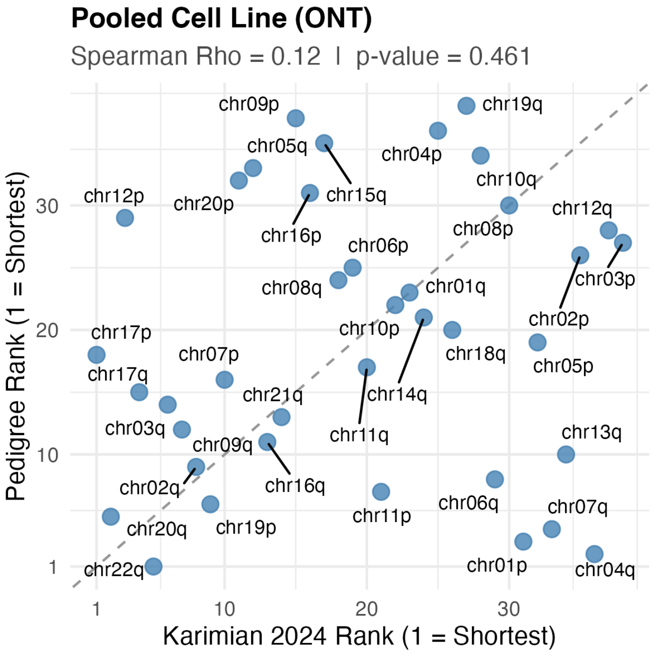

**1. Blood samples**

| SampleID | Rho | P_Value | Arms_Analyzed |
|---|---|---|---|
| M-PAN027.blood.PacBio | 0.41 | 0.011 | 38 |
| D-PAN028.blood.PacBio | 0.37 | 0.022 | 38 |
| GF-PAN011.blood.PacBio | 0.34 | 0.040 | 38 |
| GM-PAN010.blood.PacBio | 0.32 | 0.052 | 37 |

**2. Cell Line**

| SampleID | Rho | P_Value | Arms_Analyzed |
|---|---|---|---|
| D-PAN028.cell.ONT | 0.07 | 0.684 | 38 |
| M-PAN027.cell.ONT | 0.07 | 0.686 | 38 |
| M-PAN027.cell.ONT.dup | 0.06 | 0.713 | 38 |
| M-PAN027.cell.PacBio | 0.09 | 0.581 | 38 |
| GF-PAN011.cell.ONT | -0.07 | 0.657 | 38 |
| GM-PAN010.cell.ONT | 0.40 | 0.014 | 38 |

**3. Statistics for the 4 blood PacBio samples and the 3 ONT cell line samples combined:**

| Group | Rho | P_Value | Arms_Analyzed |
|---|---|---|---|
| Blood PacBio (4) | 0.45 | 0.0049 | 38 |
| Cell Line ONT (3) | 0.12 | 0.4609 | 38 |

Supplementary Figure 22: Haplotype- and chromosome-specific telomere lengths

**HAPLOTYPE- and CHROMOSOME-SPECIFIC TELOMERE LENGTHS**

**(A)** Correlation between end-specific telomere lengths between Gen2 and Gen3. **(B)** To verify that our correlation was not driven by chromosome-specific length (rather than inherited haplotypes traced using subtelomeric regions), we swapped the assigned haplotypes from correct to incorrect ones. There was a weak correlation of r=0.19, which is based on matched chromosome only (meaning that the telomeric length of a given chromosome tends to weakly correlate with the telomeric length of the same chromosome, even if they are unrelated by descent).

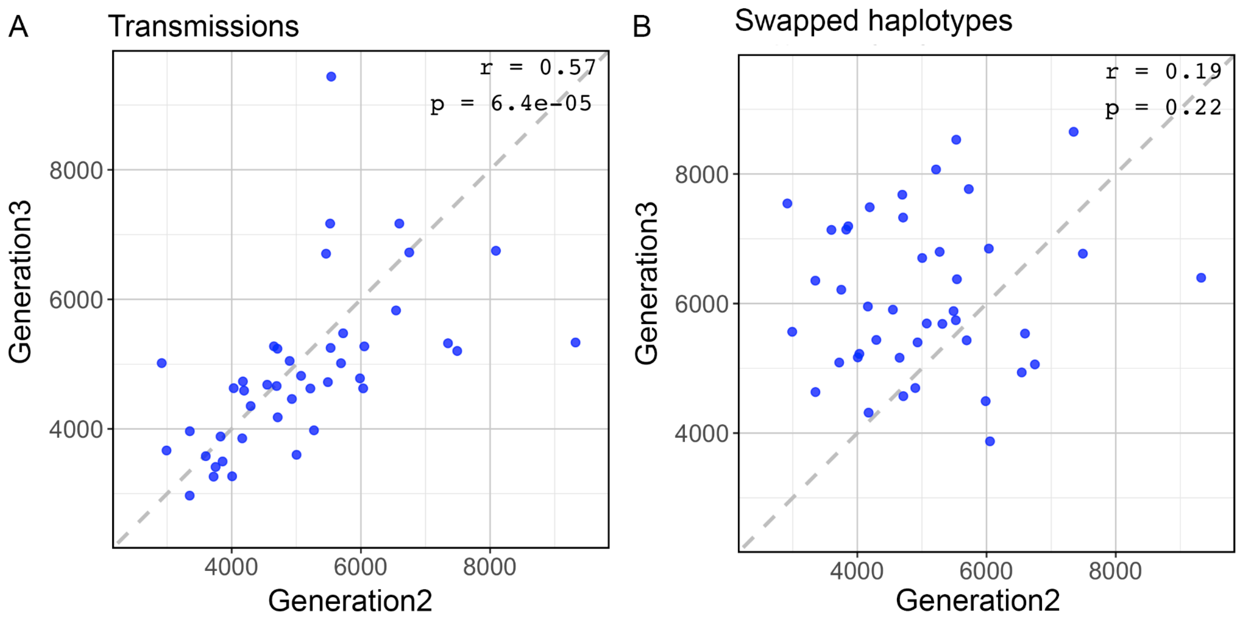

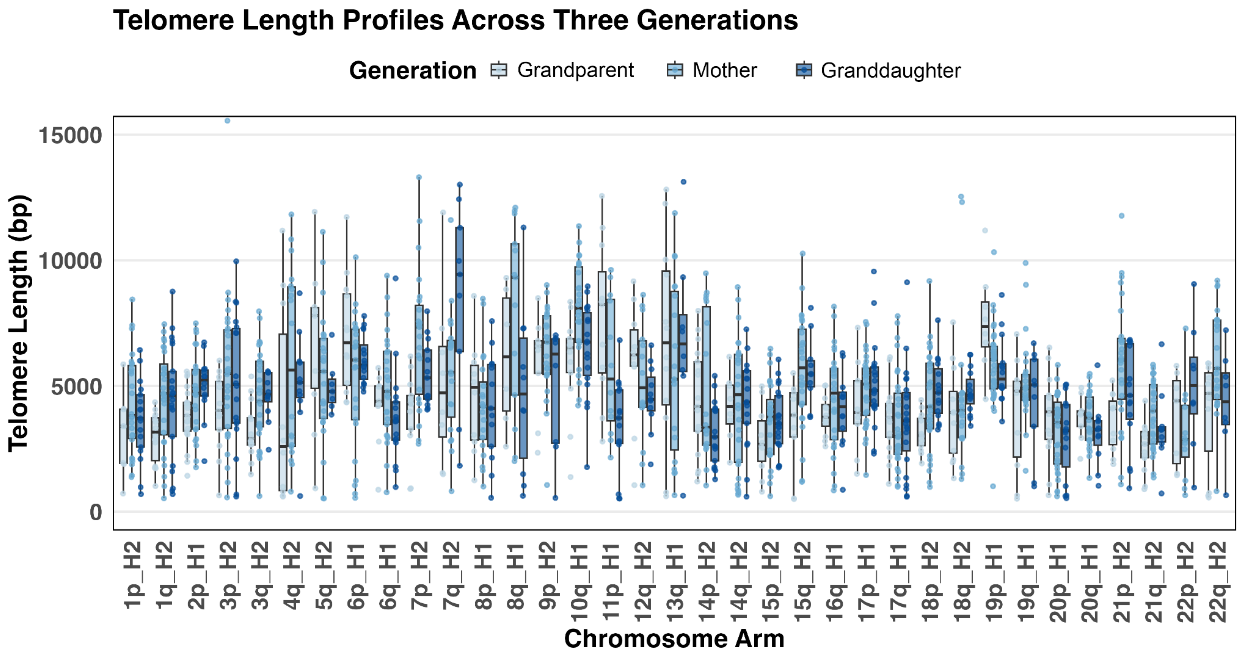


**(C)** Telomeric lengths for all chromosomes with at least 5 PacBio reads in all three generations.

#### Supplementary Figure 23: HORhap variants

HORhaps, variants of higher-order repeats sharing characteristic sequence or structural variants that define local haplotype structure within centromeric arrays, shown both in a linear representation, as well as a pie chart summarizing the proportions of individual variants within an array. The centromeres transmitted across the three generations are marked with an asterisk '*'.

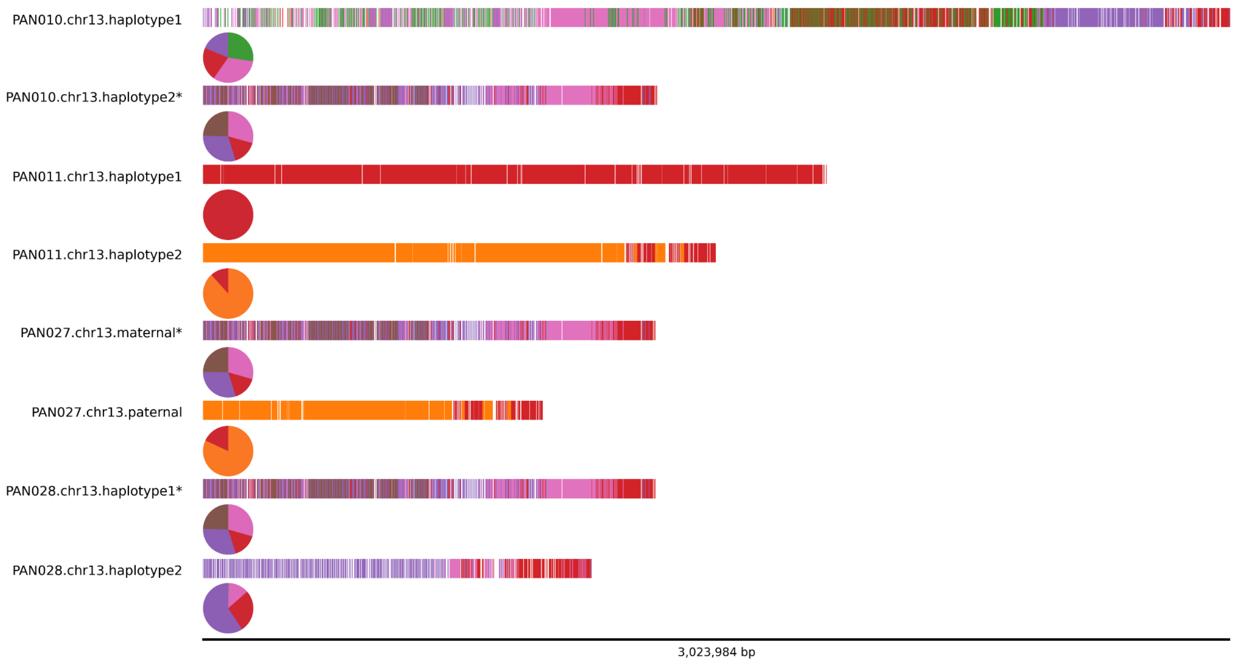

#### Supplementary Figure 24: Switch error candidates

To identify potential switch error candidates, we compared the PAN027 variants within 100kb windows to all haplotypes that could potentially be sources of a switch error. This means that for every variant in the mother, we checked the second haplotype (by lifting over the variant from the haplotype carrying the variant in the mother to the second haplotype in the mother; and by lifting over the variant from the haplotype carrying the variant in the grandparents to the second haplotype in the grandparents). We identified possible switch error regions using a clustering approach, defining such regions as any cluster with a maximum inter-variant distance of 20kb and at least eight switch-error–prone variants. We selected these parameters empirically: increasing the distance threshold beyond 20kb had minimal impact on cluster structure; while a minimum cluster size of eight variants was selected following manual review of switch-error–prone clusters in PAN027, by identifying clusters that plausibly represented switch-error regions and choosing parameters that retained them. The Figure below shows the variants in PAN027, with the clusters flagged as potential maternal switch errors in red, and paternal in blue. Maternal variants outside of these clusters were plotted in orange, and paternal variants outside of these clusters were plotted in green.

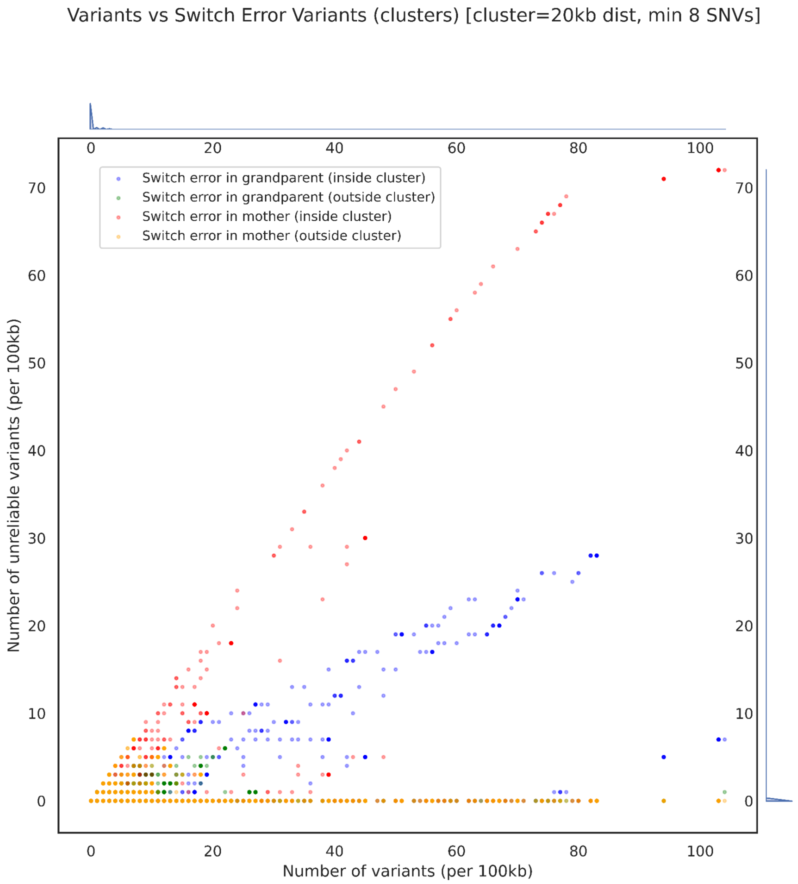

#### Supplemental Note 1: Assembly generation

Alongside verkko 2.0 assemblies, we additionally generated “hifiasm assemblies” using the same data, motivated by the observation that the breakpoints and gaps between the two assemblers frequently differ, and one can be used for patching the other. The accurate reads are a critical component for the initial graph building, and the subsequent scaffolding with ultra-long reads. Because HiFi reads have known coverage dropout regions, such as gaps in GA-rich regions, we additionally spiked duplex reads (generated by the consensus of forward and reverse reads hence increasing the accuracy) by Oxford Nanopore into the accurate portion of the reads, alongside HiFi. This allowed us to generate another set of assemblies, henceforth referred to as “duplex assemblies”, that have a superior resolution of satellite-rich p-arms of acrocentric chromosomes. While the vast majority of chromosomes assembled T2T (~32 for verkko assemblies and ~39 for duplex assemblies with verkko 2.0), the remaining chromosomes typically broke into two or three pieces, especially in centromeres, human satellites, or in close proximity to telomeres (missing telomeric repeat). Intriguingly, we didn’t find an enrichment of segmental duplication in the latest verkko 2.0 assemblies, compared to earlier versions.

We followed the traditional T2T recipe consisting of the long accurate reads, ultra-long reads, and HiC reads for the scaffolding and/or phasing. For the accurate reads, we used HiFi reads basecalled with DeepConsensus v1.2 (with coverages between 40x-70x depending on the sample), and for ultra-long reads, we generated over ~70x of ONT-UL reads, prepared with HMW protocol. All assemblies were phased with Hi-C mode of verkko, and we additionally labeled all scaffolds in PAN027 (mother) as maternal or paternal, using parent-specific unique Illumina k-mers.

We used verkko2.0 assemblies as our initial set and fixed the existing breakpoints with hifiasm whenever available. In the second iteration, we fixed the remaining breakpoints with duplex assemblies. Moreover, we replaced p-arms of all acrocentric chromosomes with the duplex version of herein. This initial manual curation enabled us to transform the learned lessons into an automated process.

#### Supplemental Note 2: Panpatch

We implemented a tool called Panpatch (https://github.com/glennhickey/panpatch) that attempts to automate the assembly patching process. It takes as input a pangenome graph generated with minigraph-cactus [2] of chromosomal assemblies for the *same sample*, along with a ranking of the assemblies. A reference assembly that spans the entire chromosome in a single contig or scaffold must also be specified. If none of the input assemblies for the target sample meet this criteria, then a reference genome such as T2T-CHM13 can be used (and needs to be included in the pangenome graph as well). Note that some complex regions, such as centromeres, may not be patchable if a non-sample reference is used since they will not align well enough within the graph. Panpatch then attempts to thread the highest priority (rank=0) assembly across the reference. If a gap (represented by one or more Ns) or contig break is encountered, it will search for the highest-priority assembly that spans the gap or break and incorporate it into the patch. If the entire chromosome can be traversed in this way, the patched contig is returned.

1. PAN027 Verkko assembly was corrected using a HERRO-corrected assembly of the same sample, as well as the Verkko assembly of PAN011.
2. Ten HPRC samples (from the top of this list https://s3-us-west-2.amazonaws.com/human-pangenomics/index.html?prefix=submissions/DC27718F-5F38-43B0-9A78-270F395F13E8--INT_ASM_PRODUCTION) were selected. Corresponding verkko assemblies (from https://s3-us-west-2.amazonaws.com/human-pangenomics/index.html?prefix=submissions/6807247E-4F71-45D8-AECE-9E5813BA1D9F--verkko-v2.2.1-release2_asms) were patched by the release/hifiasm assembly using *panpatch* and T2T-CHM13 as a reference.
3. PAN010, PAN011 PAN027 and PAN028 Verkko assemblies were patched using hifiasm (rank 1) and duplex (rank 2) assemblies, with sample-specific initial references.

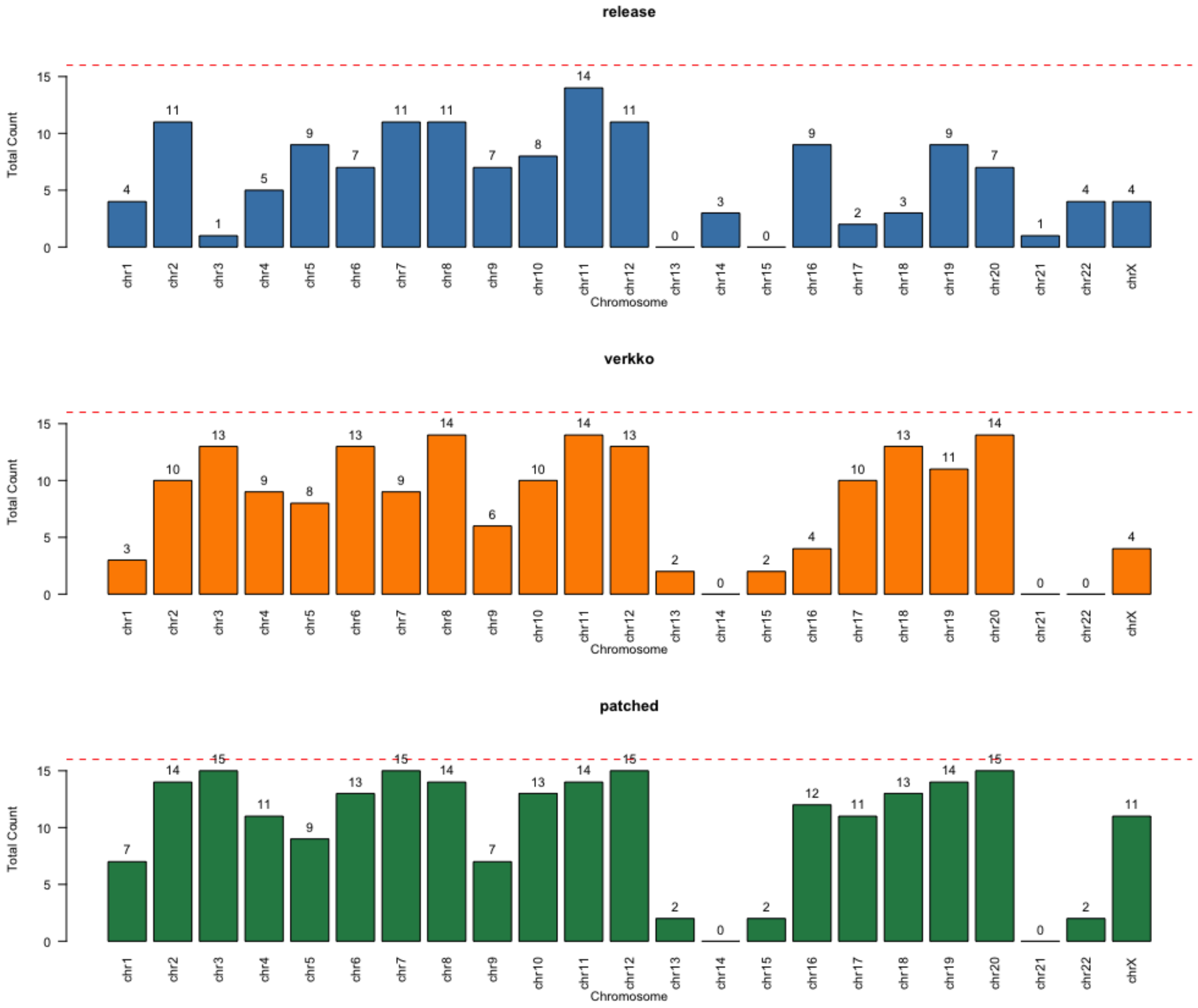


The barplots are showing T2T contigs. The verkko assemblies (orange) were patched with the release/hifiasm assemblies (blue) to produce the more contiguous assemblies (green)

#### Supplemental Note 3: Transposable element analysis

Transposable element analysis focused on de novo insertion identification inside syntenic blocks shared across a three-generation pedigree (only regions inherited from grandparents to granddaughter). By comparing shared blocks, we identified insertions longer than 50bp and determined whether these insertions represent novel TE copies by intersecting with repeat annotation created by RepeatMasker. This resulted in 6 insertion candidates in generation 1 (grandparents to mother) and 5 candidates in generation 2 (mother to granddaughter). After manually checking each candidate in Integrative Genomics Viewer (IGV), all candidates were found to be a result of incorrect haplotype phasing (element was present in the second haplotype of the ancestor, which can be a result of a switch error or gene conversion event) or identified as an incorrect variant call.

#### Supplemental Note 4: 13p recombination event

To investigate meiotic exchange within the HSat1A array, we identified all 200-bp k-mers unique to either the PAN010 haplotype 1 or haplotype 2 array. This analysis revealed that the HSat1A array inherited in PAN027 corresponded to the PAN010 haplotype 1 lineage. We localized the putative meiotic recombination breakpoint to a ~800 bp segment at the 3′ end of the array, directly adjacent to the centromere, where the two PAN010 haplotypes are nearly identical (2 differences per 1,000 sites). Examination of adjacent HORhap patterns provided further support for a transition to the PAN010 haplotype 2. Within this terminal ~800 bp segment of the HSat1A array, we observed a sharp increase in de novo variation, including 21 novel variants (two of which were indels).

Additionally, we validated that our acrocentric chromosome of chr13 was correct, by comparing our assignment with the automatic scaffolding by verkko 2.0, and comparing the haplotype matches with moddotplot. This revealed that our scaffolded haplotypes, assigned with poreC, matched the scaffolded haplotypes as reported by verkko.

#### Supplemental Note 5: Telomere Length Differences in LCLs and Blood

Telomere length was assessed in a three-generation pedigree (PAN010, grandmother; PAN011, grandfather; PAN027, mother; PAN028, granddaughter). DNA was obtained from blood and sequenced with PacBio individuals, and LCL, sequences by Oxford Nanopore Technologies (ONT) ultra-long protocols. In addition, PacBio HiFi Data for the mother's LCL was available. These data provided the opportunity to compare telomere length estimation across sequencing platforms as well as between blood and LCL sources.

| Method used for sequencing | HiFi PacBio | HiFi PacBio | Ultra Long.ONT |
|---|---|---|---|
| Source of gDNA | Blood | LCLs | LCLs |
| grandmother (PAN010) | 47X | - | 169X (71X 100kb+) |
| grandfather (PAN011) | 44X | - | 180X (70X 100kb+) |
| mother (PAN027) | 70X | 64X | 189X (70X 100kb+) |
| granddaughter (PAN028) | 44X | - | 191X (73X 100kb+) |

**Figure S5.1. The available datasets and their coverage.**

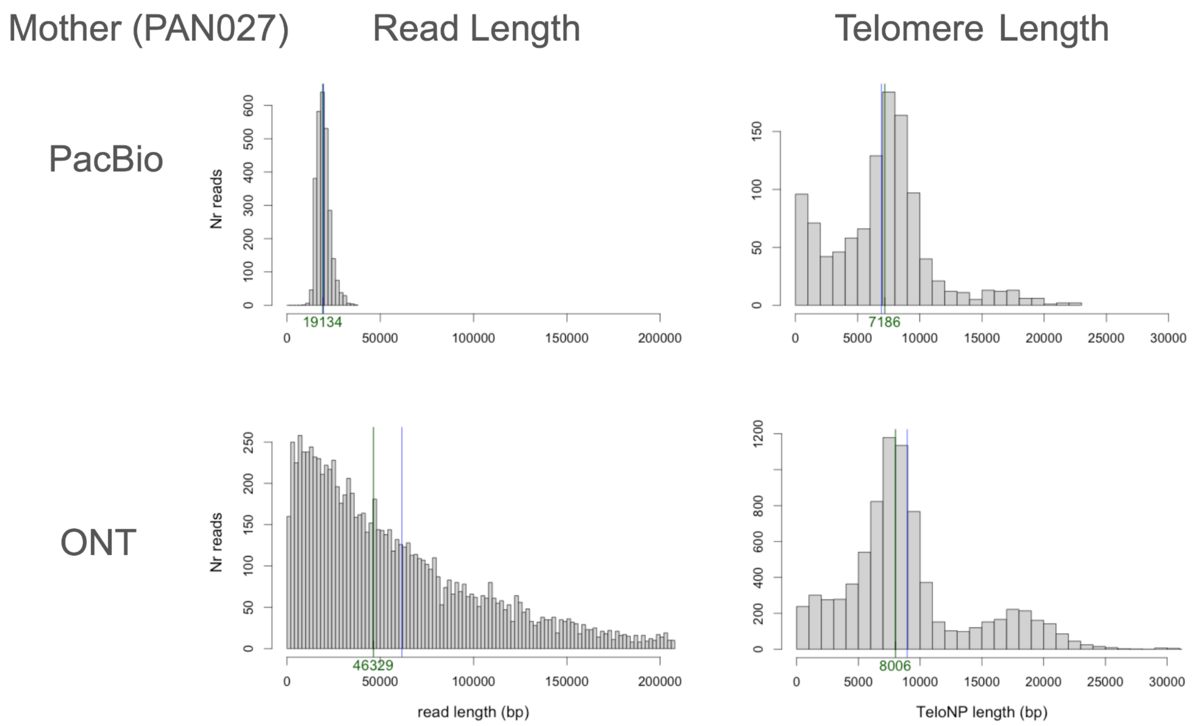


**Figure S5.2. A comparison of read length and telomere lengths for LCLs in the mother**
Our telomere profiling pipeline was applied to both HiFi and ONT datasets from the pedigree. For the mother's LCLs we had available both ONT and HiFi sequencing data. A comparison of sequencing platforms revealed a median read length of 19 kb for HiFi and 46 kb for ONT, reflecting differences in library preparation. Despite these distinct size distributions, median telomere length estimates were similar (7,186 bp for HiFi and 8006 bp for ONT).

When comparing DNA sources, telomeres derived from blood consistently fell within the expected range for human blood samples as seen in (Karimian et al. 2024). In contrast, telomere lengths from LCLs were systematically longer for all individuals except for the Grandmother. This finding is consistent with the biology of EBV-transformed lymphoblastoid cell lines, where telomeres shorten following infection but survivor clones emerge that maintain length through telomerase upregulation or alternative lengthening of telomeres (ALT) (Counter et al. 1994).

Taken together, these analyses show that both PacBio HiFi and ONT ultra-long sequencing can be used for bulk telomere length estimation, though ONT protocols better preserve extremely long telomeric molecules. More critically, they demonstrate that LCLs sometimes yield artificially elongated telomeres relative to blood, underscoring the need for caution when interpreting telomere length from LCLs. For accurate telomere length measurements, blood remains the more appropriate source of DNA.

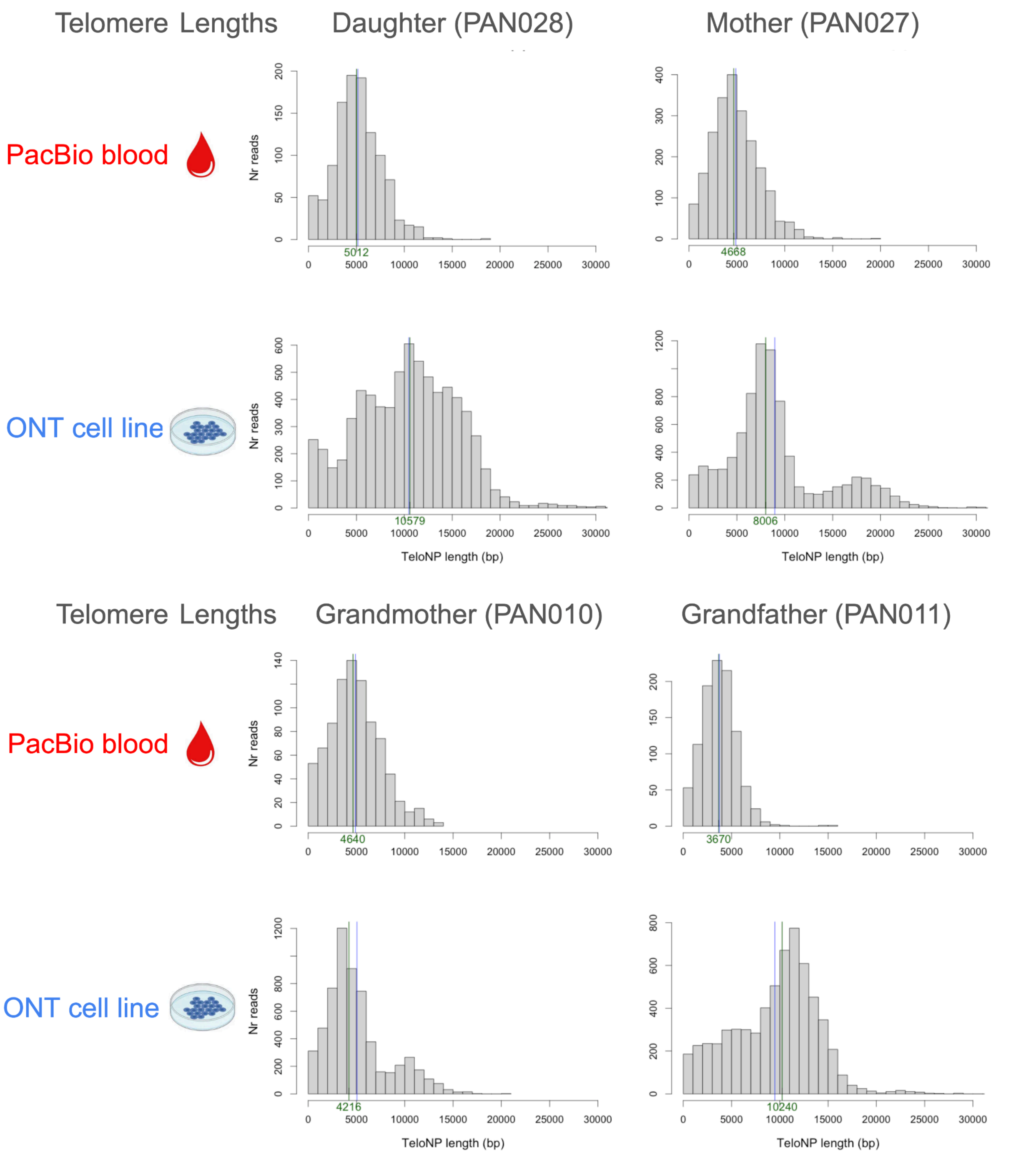


**Figure S5.3. A comparison of telomere lengths between Blood and LCLs for all four family members.**
