## Supplementary material for "Complete genomes of a multi-generational pedigree to expand studies of genetic and epigenetic inheritance": 2_WashU_Consent_Pangenome Project WUSM ICF 3.10.20.pdf

Washington University in St. Louis

FOR IRB USE ONLY
IRB ID #: 201905060
APPROVAL DATE: 05/11/21
RELEASED DATE: 05/13/21
EXPIRATION DATE: 05/10/22

### INFORMED CONSENT DOCUMENT

**Project Title: The Pangenome Project: Improving the diversity of the human reference genome**

**Principal Investigator: Nathan Stitziel, MD, PhD**

**Research Team Contact: Teresa Roediger, CCRP, CCRM, 314-362-4502**

This consent form describes the research study and helps you decide if you want to participate. It provides important information about what you will be asked to do during the study, about the risks and benefits of the study, and about your rights and responsibilities as a research participant. By signing this form you are agreeing to participate in this study.

- You should read and understand the information in this document including the procedures, risks and potential benefits.
- If you have questions about anything in this form, you should ask the research team for more information before you agree to participate.
- You may also wish to talk to your family or friends about your participation in this study.
- Do not agree to participate in this study unless the research team has answered your questions and you decide that you want to be part of this study.

This is a research study conducted by Nathan Stitziel, MD, PhD having to do with improving the human reference genome**.** You should carefully consider the information in this consent document and discuss it with the research team. Be sure you understand why you might want to participate, or why you might not want to participate. You may choose to participate or not.

If you agree and sign this consent, you will be volunteering to participate in the research study. All of the information below will be explained and is listed in more detail in the consent document below. The research team must give you a copy of this signed consent document.

**How will this study affect me?**

- The purpose of this study is to improving the human reference genome, specifically for individuals of African American ancestry.
- As a voluntary participant, you will be asked to spend approximately 30 minutes for a one-time blood draw.
- You were selected because you are of African American ancestry.
- You will be in this study for the amount of time it takes to obtain consent and a blood specimen.
- You will need to come to The Clinical and Translational Research Unit (CTRU) located on the 5th floor of The Center for Outpatient Health.
- The main risks to you are bruising, pain, or bleeding at the site of the needle stick. Rarely, people feel lightheaded or faint when their blood is drawn.

- You will be paid $75 gift card for participating in this study. You will not have costs for participating.
- If you withdraw from the study, the research team may continue to use information already collected about you in this study.

**WHAT IS THE PURPOSE OF THIS STUDY?**

Genes are the basic "instruction book" for the cells that make up our bodies and are made out of DNA. Although the DNA of a person is more than 99% the same as the DNA of any other person, no two people have exactly the same DNA except identical twins. Difference in DNA is called genetic variation. Genetic variation explains some of the physical differences among people, and partially explains why some people get diseases like cancer, diabetes, asthma, and depression, while others do not. Such diseases may also be affected by factors like diet, exercise, smoking, and pollution in the environment, which makes it hard to figure out which genes affect the diseases.

To study genetic variation, researchers routinely access the human reference genome. The current reference genome (available on the internet at genome.ucsc.edu along with other websites) was mostly (70%) made from reading the DNA of a single individual which causes problems when researchers are trying to interpret genetic variation from diverse populations.

The purpose of "The Pangenome Project" is to create a reference genome that is more representative of the genetic variation that exists in humans. We will do this by studying the DNA/RNA in blood samples collected from many people in the St. Louis area who self-identify as African American, and then putting all of this information in scientific databases on the internet.  These scientific databases will be kept for a long time, and many future researchers around the world will use them to help find genes and genetic variants related to health and disease.

The scientific databases we develop for this project will not include any medical information, but they will still be useful to help future researchers learn about health and disease. In the future, for a disease (such as diabetes), researchers will study different sets of samples—some from people who have the disease and some from people who do not, and look for areas in the DNA where the patterns of variation differ between the two groups. This will give them a clue that those areas might contain genes that affect the disease. We will also measure how much of each gene is being made by your body. They will then use the scientific databases we develop for this project to look at the genetic variants in those regions, to help figure out which genes might affect the disease. They can then study how the genes work and eventually find better ways to prevent, diagnose, and treat the disease.

Researchers will also use the scientific databases to learn more about how different people respond to different drugs, and about how traits (like long life), or behaviors (like addiction), differ between people. Other future studies that use scientific databases and the samples themselves will help researchers understand even more about human genetic variation and other important biological questions.

This is a research project, not medical care. You should see your health care provider for any scheduled visits or if you have a health problem or medical question.

#### WHAT WILL HAPPEN DURING THIS STUDY?

We will first ask you a few questions to determine whether you are eligible to give a sample, such as your age, race and whether any of your relatives already gave samples for this project.

If you agree to participate in this study we will ask for a sample of blood (4 tubes including one for preserving white blood cells and one PAX tube for preserving RNA). We will obtain a blood sample of approximately 35 milliliters or 3 tablespoons. We will use this blood to sequence all of your DNA/RNA. We will not collect your name or any medical information about you.

As part of this study, we are obtaining blood samples from you. These may be used for commercial profit (even if we remove your identifiable information.) There are no plans to provide financial compensation to you should this occur. By allowing us to use your blood or data you give up any property rights you may have in the blood samples.

#### WILL YOU SAVE MY RESEARCH INFORMATION AND/OR BIOSPECIMENS TO USE IN FUTURE RESEARCH STUDIES?

As part of this study, we are obtaining blood and/or data from you. We would like to use the blood and/or data we are obtaining in this study for studies going on right now as well as studies that are conducted in the future. Your blood and/or data may also be used for broad sharing throughout the research community. This means your blood and/or data may be used for any sort of research including research to develop investigational tests, treatments, drugs or devices that are not yet approved by the U.S. Food and Drug Administration. These researchers may be at Washington University, at other research centers and institutions, or commercial sponsors of research. It is unlikely that what we learn from these studies will have a direct benefit to you. There are no plans to provide financial compensation to you should this occur. By allowing us to use your blood and/or data you give up any property rights you may have in the blood and/or data.

We will remove all identifiers from your private information and your blood and data and then use the information and your blood and data for future research studies or share them with other researchers for their future research. If this occurs we will not ask you for additional consent for these uses of your information or blood and data.

We will send your blood cells to a research repository to be processed and stored. The only information we will include with the sample is your sex and race.

The research repository will make a cell line from your cells. This means that cells from your blood will be grown in culture indefinitely. This is to ensure that researchers can get an unlimited amount of your DNA/RNA for a long time, perhaps forever. The cells derived from your blood sample can also be transformed into different kinds of cells. For example, blood cells in this study could be transformed into nerve cells or muscle cells.

The repository will share cell lines and DNA from the samples with other researchers around the world so they can be used in many future studies. These future researchers may include researchers in universities, hospitals, non-profit groups, companies, and government laboratories. These researchers will follow the laws and guidelines that apply to biomedical research.

One way in which we may share your data with others is by putting it into a large database of information, called a data repository. If your data is placed in one of these repositories it will be placed in the "unrestricted access" or "open" portion of the repository. This means the information will be widely available not only to researchers, but to any member of the general public. There are no monitors or controls placed on who can or cannot access and download these data and there are no permissions needed for anyone to use the data. Once your data is placed in the database, it may not be possible to remove it. Even if it can be removed from the database, it will not be possible to retrieve any information that has already been accessed and/or downloaded. Although these data will not have your name or other identifying information associated with it, it is still possible that someone may be able to trace these data back to you because genetic information is unique.

Individuals accessing the data might read all of the genetic information in your DNA/RNA by sequencing, which will provide a detailed description of your DNA/RNA and is sometimes called whole genome sequencing. They may compare it to other DNA/RNA to study questions that relate to the biology of DNA/RNA, how DNA/RNA variation arise, how evolution works, the composition and size of human groups, how people from different populations are related to each other, and how DNA/RNA variation is related to health and disease.

Your blood and data will be stored without your name or any other kind of link that would enable us to identify which sample(s) or data are yours. Therefore, it will be available for use in future research studies indefinitely and cannot be removed.

#### HOW MANY PEOPLE WILL PARTICIPATE?

Approximately 30 people will take part in this study conducted by investigators at Washington University.

#### HOW LONG WILL I BE IN THIS STUDY?

If you agree to take part in this study, your involvement will include a one-time blood draw, though data and/or cell lines from your participation will last indefinitely.

#### WHAT ARE THE RISKS OF THIS STUDY?

You may experience one or more of the risks indicated below from being in this study. In addition to these, there may be other unknown risks, or risks that we did not anticipate, associated with being in this study.

Risk of Blood Draw
We are drawing a small amount of blood though occasionally there can be bruising, pain, or bleeding at the site of the needle stick. Rarely, people feel lightheaded or faint when their blood is drawn. Very rarely, the vein may become red and swollen, or infected. If this happens, we can treat the problem. This is comparable to a blood draw that you may have at your doctor's office.

Risk of Genetic Testing

Although we will not collect any names or medical information, and we will take many measures to protect your privacy (see, "How will you keep my information confidential?") we will generate lots of genetic information about each person whose sample is studied. This information will be put in open access scientific databases, available on the Internet to anyone who wants to look at it. Although only experts will know how to interpret this information, there is a small chance that somebody could figure out how to connect you with the information from the study of the sample you give; the information could then be used to discriminate against you or your family members. Currently, we believe this could happen only if somebody knew that you had given a sample to be studied for this project and:

- got another sample from you, found an expert to test that sample, and then compared the genetic information from that test with the genetic information in the scientific databases;
- found an expert to compare the genetic information about you in the scientific databases with information known to have come from you (or from a family member) included in some other database developed by someone else for some other purpose; or
- found an expert to look in the scientific databases for a particular genetic variation known (or someday found) to be associated with a disease or trait that you have or carry, that others know about or can see, and that is very rare.

Any of these things would require that the person trying to link the information to you knew that you participated in the project. For this reason, to minimize these risks, you may wish to limit the number of people you tell about your participation.

There could be risks to your group or community. The name of the ethnic group the samples came from will be included with the samples and in the scientific databases. In future studies, researchers may find that certain genetic variations appear more often in people from your group than in people from other groups, and that these variations are more common in people with a certain disease. This may make some people look down on your group unfairly.

Some people may use the information from the scientific databases, or from future studies using the scientific databases, to exaggerate differences between groups for prejudiced or other bad reasons.

Others may use the information to downplay differences between groups, to say that all people's genes are about the same, so we don't need to respect the special concerns of different groups. Biology does not provide a reason for prejudice, but discrimination does exist.

We will work to make sure that the ethnic or geographic identity of your community is described as carefully as possible--in the sample collection, in the scientific databases, and in articles that project researchers write based on this research, but we cannot completely control how this information is described in publications that others write.

As technology advances, there may be new ways of linking information back to you that we cannot foresee now.

**Re-identification from a genetic sample**
While the data developed for this study is being stored without traditional identifiers (stored only with coded ID numbers, no names), there may be ways of linking the genetic materials back to you. DNA does directly identify you, so it is possible that someone could compare information in our database with information from you in another database and be able to identify you. Because more people are uploading and openly sharing their genetic information on health and genealogy websites, it is possible that someone could identify that your DNA was used in this study. It is also possible that law enforcement could use this same approach to identify you or one of your relatives.

**Genetic Research**
There is a federal law called the Genetic Information Nondiscrimination Act (GINA). In general, this law makes it illegal for health insurance companies, group health plans and employers with greater than 15 employees to discriminate against you based on your genetic information (if someone were to identify that you participated in this study, for example). However, it does not protect you against discrimination by companies that sell life insurance, disability insurance or long term-care insurance.

**Breach of Confidentiality**
One risk of participating in this study is that confidential information about you may be accidentally disclosed. We will use our best efforts to keep the information about you secure. Please see the section in this consent form titled *"How will you keep my information confidential?"* for more information.

#### WHAT ARE THE BENEFITS OF THIS STUDY?

You will not benefit from being in this study.

However, we hope that, in the future, other people might benefit from this study because it will increase our knowledge about the diversity of human genetic variation, and to enable improved interpretation and study of all human genomes.

**WILL IT COST ME ANYTHING TO BE IN THIS STUDY?**

You will not have any costs for being in this research study.

**WILL I BE PAID FOR PARTICIPATING?**

You will be paid for being in this research study. You will need to provide your social security number (SSN) in order for us to pay you. You may choose to participate without being paid if you do not wish to provide your social security number (SSN) for this purpose. You may also need to provide your address if a check will be mailed to you. If your social security number is obtained for payment purposes only, it will not be retained for research purposes.

You will receive a $75 gift card for completing the one time blood draw.

**WHO IS FUNDING THIS STUDY?**

The National Institutes of Health (NIH) is funding this research study. This means that Washington University is receiving payments from the NIH to support the activities that are required to conduct the study. No one on the research team will receive a direct payment or increase in salary from the NIH for conducting this study.

**WHAT IF I AM INJURED AS A RESULT OF THIS STUDY?**

Washington University investigators and staff will try to reduce, control, and treat any complications from this research. If you feel you are injured because of the study, please contact the investigator at Dr. Nathan Stitziel at (314) 747-8394 and/or the Human Research Protection Office at 1-(800)-438-0445.

Decisions about whether payment for medical treatment for injuries relating to your participation in research will be made by Washington University**.** If you need to seek medical care for a research-related injury, please notify the investigator as soon as possible.

**HOW WILL YOU KEEP MY INFORMATION CONFIDENTIAL?**

Other people such as those indicated below may become aware of your participation in this study and may inspect and copy records pertaining to this research. Some of these records could contain information that personally identifies you.

- Government representatives (including the Office for Human Research Protections) to complete federal or state responsibilities
- The U.S. Food and Drug Administration
- The National Institutes of Health
- People who use the registry
- Hospital or University representatives to complete Hospital or University responsibilities
- Information about your participation in this study may be documented in your health care records

and will be available to anyone with access to your health care record, including your health insurance company. This information may also be released as part of a release of information request.

- The last four digits of your social security number may be used in hospital or University systems to track billing information for research procedures.
- Washington University's Institutional Review Board (a committee that oversees the conduct of research involving human participants) and the Human Research Protection Office. The Institutional Review Board has reviewed and approved this study.
- Any report or article that we write will not include information that can directly identify you. The journals that publish these reports or articles require that we share your information that was collected for this study with others to make sure the results of this study are correct and help develop new ideas for research. Your information will be shared in a way that cannot directly identify you.

To help protect your confidentiality, we will store your signed consent form in a locked file; only members of the study team at Washington University School of Medicine will have access to this file. We will not collect your name or any other identifying information (such as address, birth date, or Social Security number) or give your sample a code number that could identify you. We will collect more samples than we will use, so that nobody - not even you or us - will know for sure whether your sample was used or if any of the information in the scientific databases came from your sample. Samples that are not used will be destroyed. Because of these measures, it will be very hard for anyone who looks at any of the scientific databases to know which information came from you, or even that any information in the scientific databases came from you.

To further protect your privacy, this research is covered by a Certificate of Confidentiality from the federal government. This means that the researchers can refuse to disclose information that may identify you in any legal or court proceeding or to anyone who is not connected with the research except if:

- there is a law that requires disclosure, such as to report child abuse and neglect, or harm to self or others;
- you give permission to disclose your information, including as described in this consent form; or
- it is used for other scientific research allowed by federal law.

You have the right to share your information or involvement in this study with anyone at any time. You may also give the research team permission to disclose your information to a third party or any other person not connected with the research.

#### **Are there additional protections for my health information?**

Protected Health Information (PHI) is health information that identifies you. PHI is protected by federal law under HIPAA (the Health Insurance Portability and Accountability Act). To take part in this research, you must give the research team permission to use and disclose (share) your PHI for the study as explained in this consent form. The research team will follow state and federal laws and may share your health information with the agencies and people listed under the previous section titled, "How will you keep my information confidential?"

Once your health information is shared with someone outside of the research team, it may no longer be protected by HIPAA.

The research team will only use and share your information as talked about in this form or as permitted or required by law. When possible, the research team will make sure information cannot be linked to you (de-identified). Once information is de-identified, it may be used and shared for other purposes not discussed in this consent form. If you have questions or concerns about your privacy and the use of your PHI, please contact the University's Privacy Officer at 866-747-4975.

Although you will not be allowed to see the study information, you may be given access to your health care records by contacting your health care provider.

**If you decide not to sign this form, it will not affect**

- your treatment or the care given by your health provider.
- your insurance payment or enrollment in any health plans.
- any benefits to which you are entitled.

However, it will not be possible for you to take part in the study.

**If you sign this form:**

- You authorize the use of your PHI for this research
- This authorization does not expire.
- You may later change your mind and not let the research team use or share your information (you may revoke your authorization).
  - To revoke your authorization, complete the withdrawal letter, found in the Participant section of the Human Research Protection Office website at https://hrpo.wustl.edu/participants/withdrawing-from-a-study/ or you may request that the investigator send you a copy of the letter.
    - **If you revoke your authorization:**
      - The research team may only use and share information already collected for the study.
      - Your information may still be used and shared as necessary to maintain the integrity of the research, for example, to account for a participant's withdrawal from the research study or for safety reasons.
      - You will not be allowed to continue to participate in the study.

#### IS BEING IN THIS STUDY VOLUNTARY?

Taking part in this research study is completely voluntary**.** You may choose not to take part at all. If you decide to be in this study, you may stop participating at any time. Any data that was collected as part of your participation in the study will remain as part of the study records and cannot be removed.

If you decide not to be in this study, or if you stop participating at any time, you won't be penalized or lose any benefits for which you otherwise qualify.

#### WHAT IF I DECIDE TO WITHDRAW FROM THE STUDY?

You may withdraw by telling the study team you are no longer interested in participating in the study or you may send in a withdrawal letter. A sample withdrawal letter can be found at https://hrpo.wustl.edu/participants/withdrawing-from-a-study/ under Withdrawing from a Research Study. However, your blood and data will be stored without your name or any other kind of link that would enable us to identify which sample(s) or data are yours so it will not be possible to remove your data in future research studies.

#### WHAT IF I HAVE QUESTIONS?

We encourage you to ask questions. If you have any questions about the research study itself, please contact: Nathan Stitziel, MD, PhD, (314) 747-8394**.** If you experience a research-related injury, please contact: Nathan Stitziel, MD, PhD, (314) 747-8394.

If you have questions, concerns, or complaints about your rights as a research participant, please contact the **Human Research Protection Office 1-(800)-438-0445**, or. General information about being a research participant can be found on the Human Research Protection Office web site, http://hrpo.wustl.edu. To offer input about your experiences as a research participant or to speak to someone other than the research staff, call the Human Research Protection Office.

---

This consent form is not a contract. It is a written explanation of what will happen during the study if you decide to participate. You are not waiving any legal rights by agreeing to participate in this study. As a participant you have rights and responsibilities as described in this document and including:

- To be given enough time before signing below to weigh the risks and potential benefits and decide if you want to participate without any pressure from the research team or others.
- To understand all of the information included in the document, have your questions answered, and receive an explanation of anything you do not understand.
- To follow the procedures described in this document and the instructions of the research team to the best of your ability unless you choose to stop your participation in the research study.
- To give the research team accurate and complete information.
- To tell the research team promptly about any problems you have related to your participation, or if you are unable to continue and wish to stop participating in the research study.

Your signature indicates that this research study has been explained to you, that your questions have been answered, and that you agree to take part in this study. You will receive a signed and dated copy of this form.

**Do not sign this form if today's date is after EXPIRATION DATE: 05/10/22.**

________________________________________ ______________________________
(Signature of Participant) (Date)

__________________________________________
(Participant's name – printed)

**Statement of Person Who Obtained Consent**

The information in this document has been discussed with the participant or, where appropriate, with the participant's legally authorized representative. The participant has indicated that they understand the risks, benefits, and procedures involved with participation in this research study.

________________________________________ ______________________________
(Signature of Person who Obtained Consent) (Date)

_________________________________________
(Name of Person who Obtained Consent - printed)
