## Supplementary material for "Complete genomes of a multi-generational pedigree to expand studies of genetic and epigenetic inheritance": HG06807C P4.pdf

CORIELL INSTITUTE FOR MEDICAL RESEARCH

### Cytogenomics Chromosome Analysis Report

| | |
|---|---|
| **Coriell Case ID:** | **HG06807*C** |
| **Cell Line ID:** | **MGISTL-PAN027** |
| **Passage:** | **4** |
| **Specimen Type:** | **Lymph** |
| **Species:** | **Human** |
| **Date Received:** | **09/21/2021** |
| **Banding Technique:** | **G-Banding** |
| **Cells Counted:** | **20** |
| **Cells Analyzed:** | **5** |
| **Cells Karyotyped:** | **5** |

**ISCN: 46,XX[20]**

**Additional Information:** N/A

*Small chromosome anomalies and mosaicism may not be detectable using the standard methods employed. Chromosome analysis was performed at a level of 400 bands or greater.*

*This analysis was performed for research purpose only.*

-----------------------------------------------------------------------------------------------

Reviewed by:
Access Genomics, LLC by Yoshiko Mito, PhD, FACMG Yoshiko Mito Date 10/5/2021
Cytogenomics Consultant for Coriell Institute for Medical Research

Form 0600-14 Rev F-110917

CORIELL INSTITUTE
FOR MEDICAL RESEARCH

**Cell Line ID: HG06807*C**

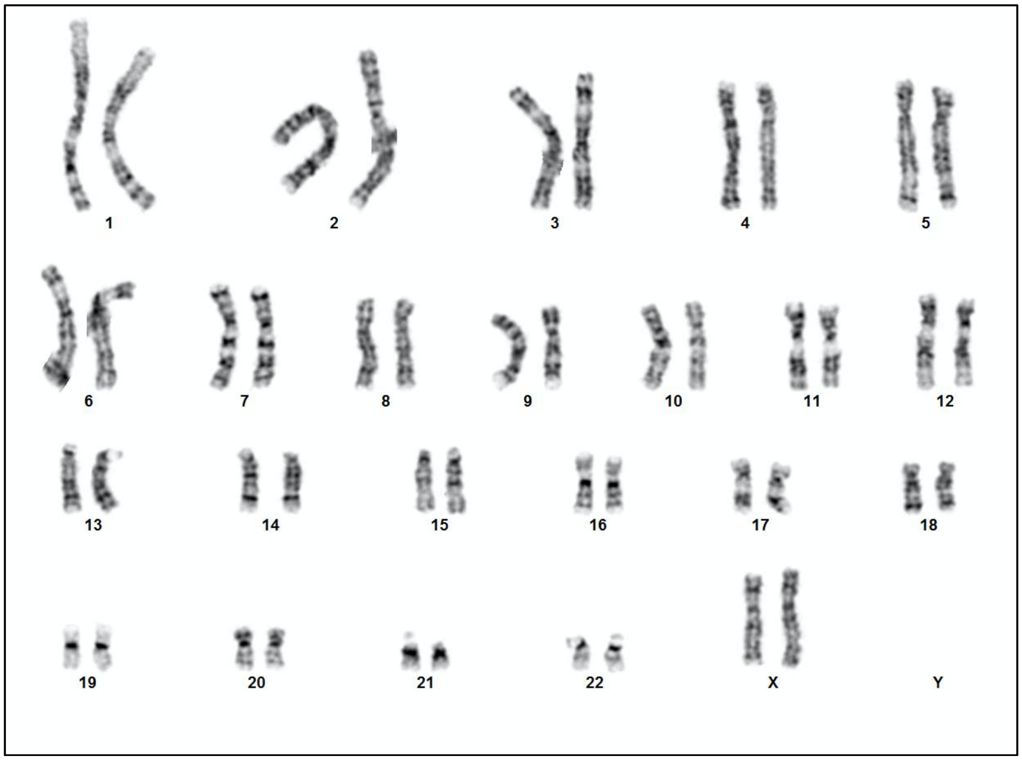


46,XX

**Cell Line ID: HG06807*C**

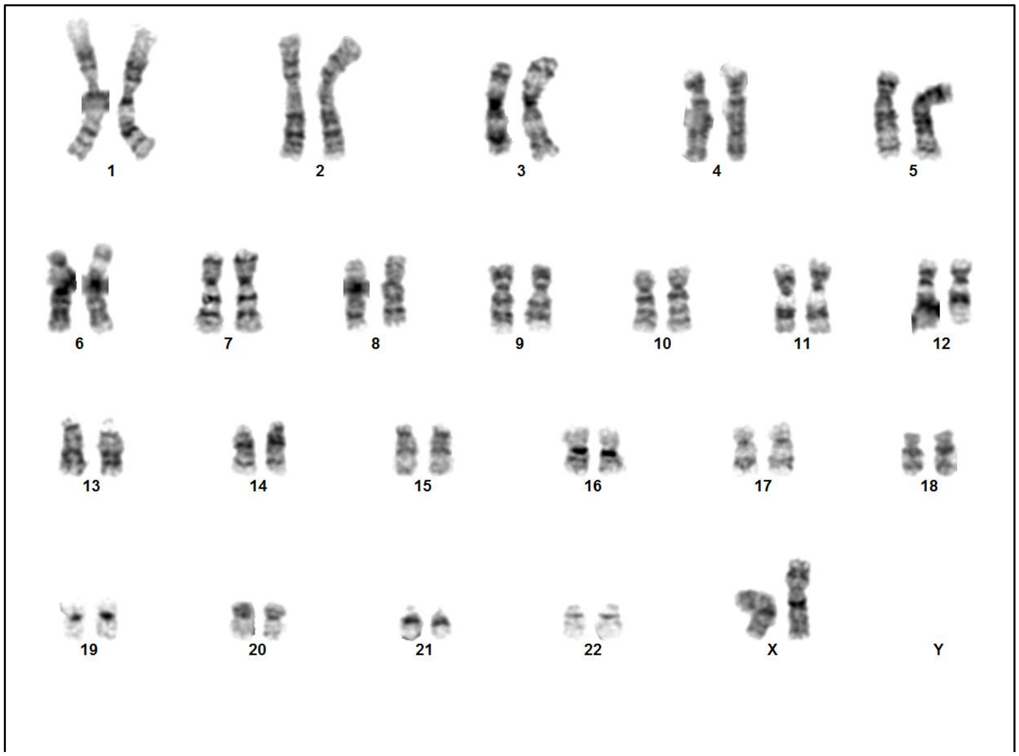


46,XX[20]
