## Supplementary material for "Complete genomes of a multi-generational pedigree to expand studies of genetic and epigenetic inheritance": HG06808B P4.pdf

### Cytogenomics Chromosome Analysis Report

| | |
|---|---|
| **Coriell Case ID:** | **HG06808*B** |
| **Cell Line ID:** | **MGISTL-PAN028** |
| **Passage:** | **4** |
| **Specimen Type:** | **Lymph** |
| **Species:** | **Human** |
| **Date Received:** | **08/17/2021** |
| **Banding Technique:** | **G-banding** |
| **Cells Counted:** | **20** |
| **Cells Analyzed:** | **9** |
| **Cells Karyotyped:** | **8** |

**ISCN: 45,X,-X[5]/46,XX[15]**

**Additional Information:** N/A

*Small chromosome anomalies and mosaicism may not be detectable using the standard methods employed. Chromosome analysis was performed at a level of 400 bands or greater.*

*This analysis was performed for research purpose only.*

-----------------------------------------------------------------------------------------------------
Reviewed by:
Access Genomics, LLC by Yoshiko Mito, PhD, FACMG Date 9/12/2021
Cytogenomics Consultant for Coriell Institute for Medical Research



**Cell Line ID: HG06808*B**

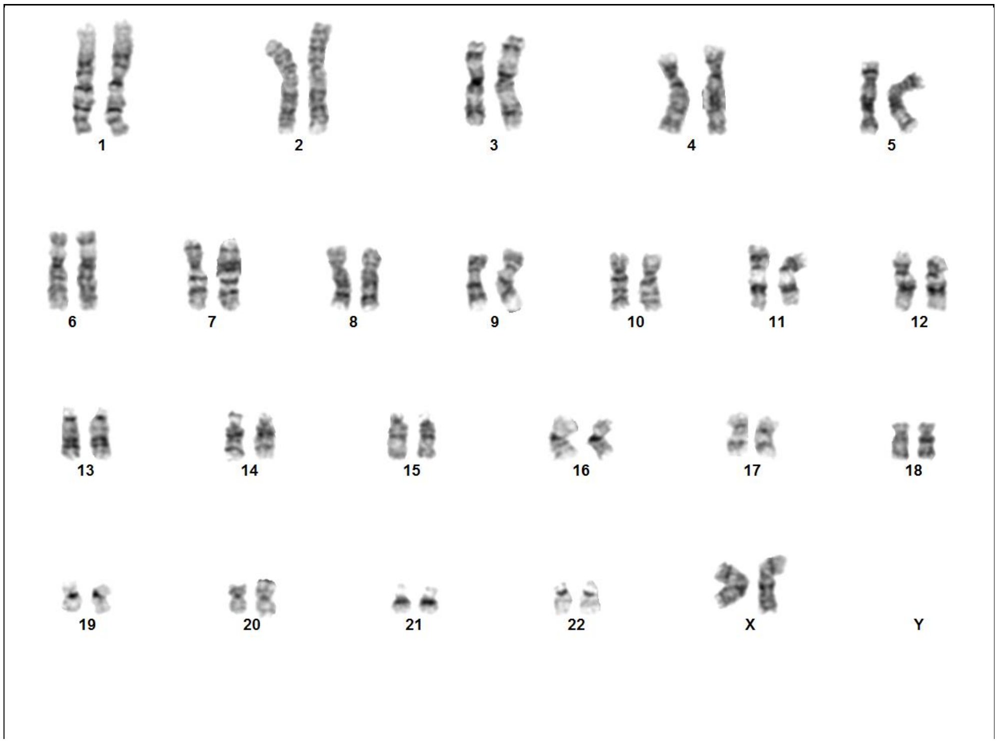


46,XX

**Cell Line ID: HG06808*B**

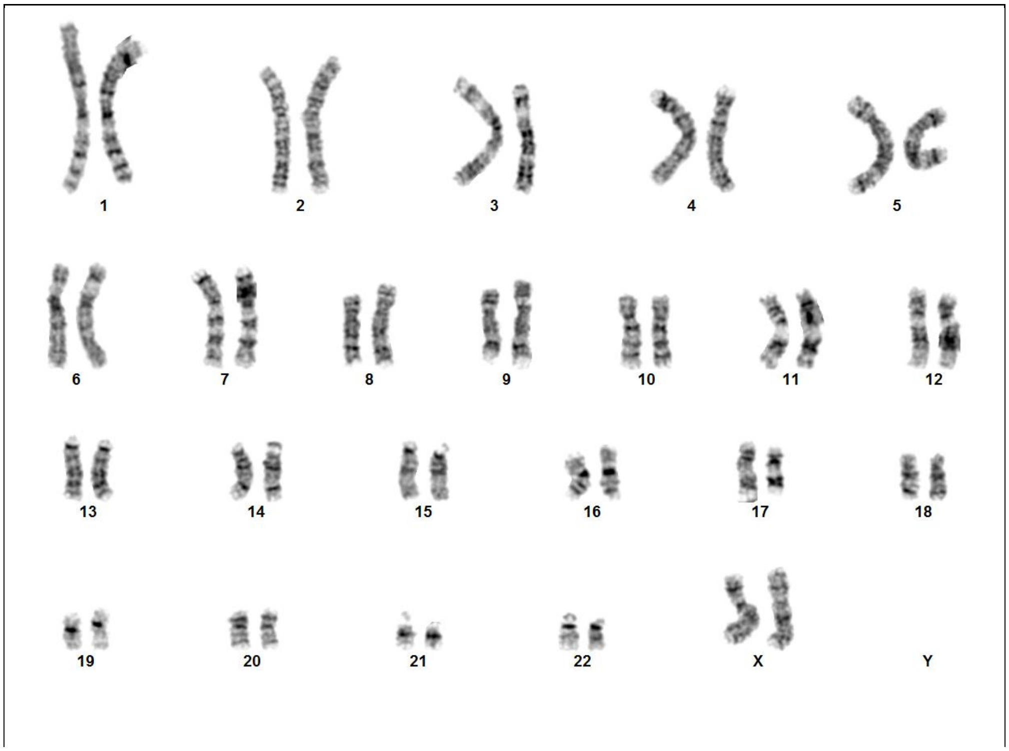


46,XX



CORIELL INSTITUTE FOR MEDICAL RESEARCH

**Cell Line ID: HG06808*B**

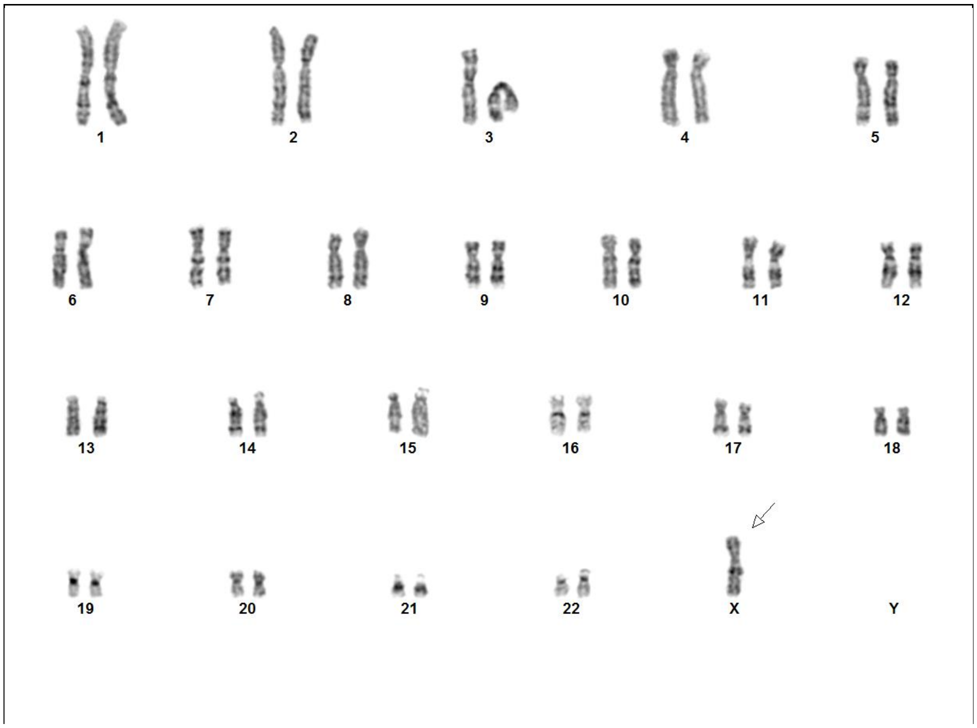


45,X,-X
